# Optimization of the experimental parameters of the ligase cycling reaction

**DOI:** 10.1101/510768

**Authors:** Niels Schlichting, Felix Reinhardt, Sven Jager, Michael Schmidt, Johannes Kabisch

## Abstract

The ligase cycling reaction (LCR) is a scarless and efficient method to assemble plasmids from fragments of DNA. This assembly method is based on the hybridization of DNA fragments with complementary oligonucleotides, so-called bridging oligos (BOs), and an experimental procedure of thermal denaturation, annealing and ligation. In this study, we explore the effect of molecular crosstalk of BOs and various experimental parameters on the LCR by utilizing a fluorescence-based screening system. The results indicate an impact of the melting temperatures of BOs on the overall success of the LCR assembly. Secondary structure inhibitors, such as DMSO and betaine, are shown to negatively impact the number of correctly assembled plasmids. Adjustments of the annealing, ligation and BO-melting temperature further improved the LCR. The optimized LCR was confirmed by validation experiments. Based on these findings, a step-by-step protocol is offered within this study to ensure a routine for high efficient LCR assemblies.

## INTRODUCTION

One of the aims of synthetic biology is to specify, design, build and test genetic circuits, and this goal requires rapid prototyping approaches to facilitate assembly and testing of a wide variety of circuits. To this end, many assembly methods were used in the last decades to build DNA constructs, e.g., Gibson assembly (1), Golden Gate assembly (2, 3), circular polymerase extension cloning (CPEC, 4), biopart assembly standard for idempotent cloning (BASIC, 5) and yeast homologous recombination (YHR, 6). Some of these methods require specific modifications of the DNA parts such as overhangs or restriction sites, which hamper the reusability, while other methods leave scars. Additionally, DNA part standardization approaches, e. g. the BioBrick-system, result in sequence redundancies which have a negative impact on assembly efficiencies (7).

By contrast, the ligase cycling reaction (LCR) fulfills the prerequisites for automated assemblies and uses phosphorylated DNA parts (8, 9, 10, 11). The assembly order is determined by single-stranded oligonucleotides building a bridge (so-called bridging oligonucleotides, BOs) between adjacent parts. Bridging oligos are typically designed based on general rules provided by the literature (8, 9). One important parameter for the LCR-assembly is the melting temperature (*T*_*m*_) of the BOs at around 70 °C for each half to facilitate optimal hybridization of template and oligonucleotide for given cycling parameters. Closely related is the free energy Δ*G*, which is assumed as the more important quantity for oligonucleotide-based biological experiments (12, 13). In the LCR, its impact is counteracted by using dimethyl sulfoxide (DMSO) and betaine to increase Δ*G* and thus to reduce secondary structures (8, 9). Nevertheless, the roles of Δ*G*-related crosstalk and the potential of Δ*G*-optimized BOs in the LCR have not been investigated so far.

The literature offers several tools regarding LCR optimization. Nowak et al. (14) provides a tool for the assembly of DNA that codes for a protein, where they return both the DNA fragments as well as the BOs to minimize unwanted effects. The tool considers codon mutations, as long as they encode the same amino acid, and is intended to be applied for LCR-based gene synthesis. Bode et al. (15) offers similar functionality. Another web-application includes the design of primers and performs *T*_*m*_ and Δ*G* cross-checks for the oligonucleotide sequences against themselves, their DNA probes and whole genomes (13) but is not applied for the LCR. Robinson et al. (16) use a BO-design-tool with an adjustable target melting temperature but without optimizing the crosstalk. An experimental perspective is given by de Kok et al. (8), where a design-of-experiment approach and multivariate data analysis were used to assess the impact of a wide range of parameters including the concentrations of the secondary structure inhibitors DMSO and betaine. The following study starts with these baseline-conditions. The LCR is investigated with the scope on the impact of the choice of BOs, their intramolecular and intermolecular crosstalk and the context of the experimental temperatures. For this, a toy-model plasmid and fluorescence-based readout is utilized (graphical abstract: Figure 1) to detect and validate the influence of all parameters and to generate new rules for an optimized LCR-assembly. Finally, the new LCR-conditions are used to assemble two additional plasmids to verify the findings.

**Figure 1.**
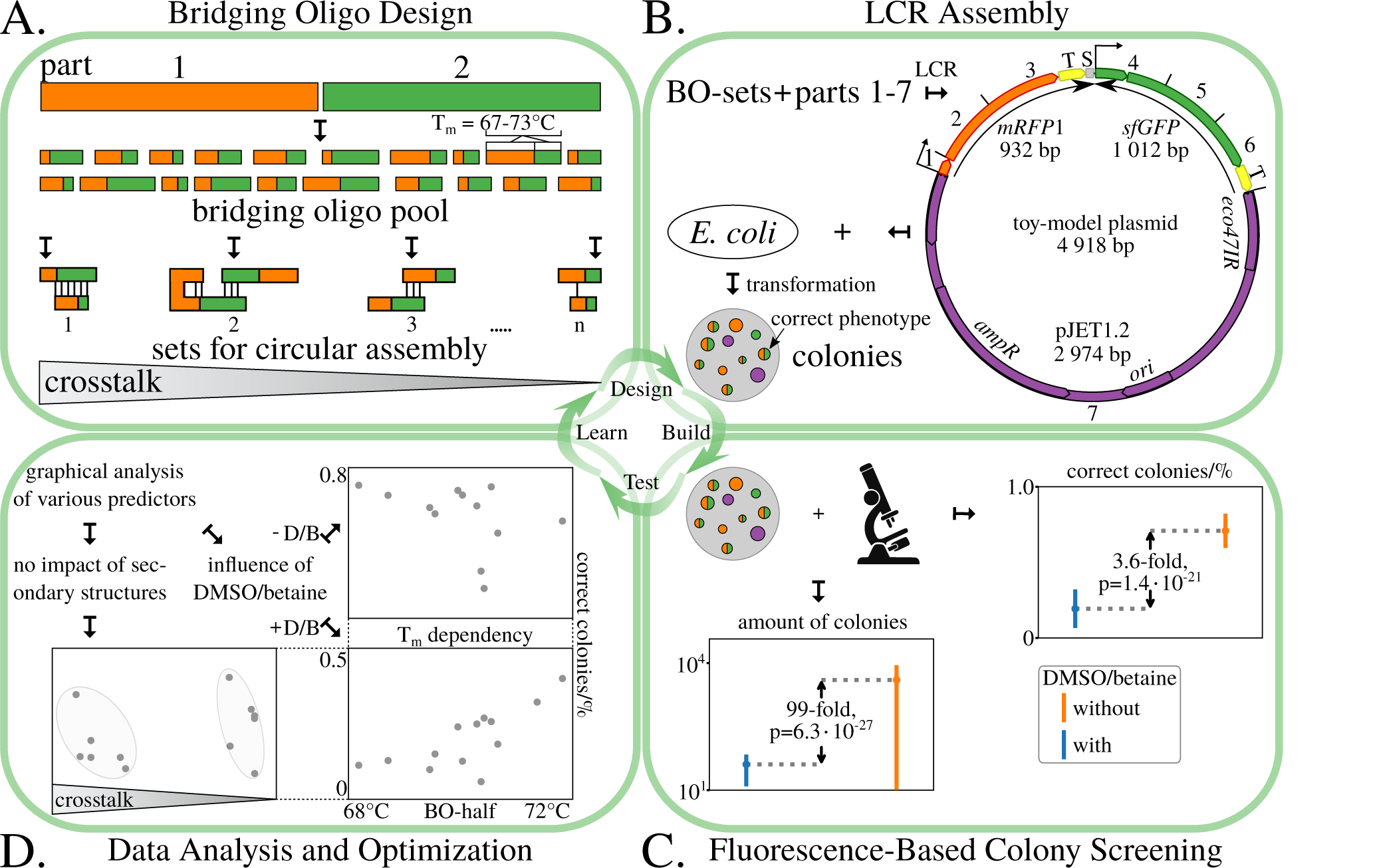
Workflow for the LCR optimization. **A**. Bridging oligo-sets (BO-sets) were designed using general design rules with the focus on Δ*G*-related BO-crosstalk while maintaining a *T*_*m*_ of 70±3°C. **B**. In total, 25 sets were designed, including one manually designed set using the software `Primer3`, and utilized for the LCR-assembly of a toy-model plasmid. The plasmid consists of seven parts with lengths in the range of 79 bp to 2974 bp and a total length of 4918 bp. It contains two genes for fluorescent proteins, *mRFP1* and *sfGFP*, and a vector. Both fluorescent protein genes were split into three subparts (*mRFP1*: parts 1-3, *sfGFP*: parts 4-6). The same terminator *BBa B0015* (T) was used twice to simulate sequence redundancies. A DNA-spacer S (37 bp) was added at the end of part 3 to avoid hybridization of the BO utilized for the ligation of parts 6 and 7. The sequences of all parts are shown in Table 1. **C**. The toy-model plasmid enables a fast and reliable fluorescence-based readout to observe the LCR efficiency and the total amount of colonies to investigate various LCR conditions. Based on this method, a significant negative impact by using the baseline LCR conditions (8 %v*/*v DMSO, 0.45 M betaine) was revealed for the seven-part toy-plasmid. The p-values were derived from a Kolmogorov-Smirnov test between the two sets of data shown in Supplementary Figures 4 and 5. **D**. With the focus on all BO-sets, no Δ*G*-related impact was detected in the baseline and DMSO/betaine-free LCR conditions. Further graphical analysis of the BO-sets revealed an association of the average BO-*T*_*m*_, the efficiency and total amount of colonies. Finally, the optimizations were applied on another split of the toy-plasmid (three-part design: *mRFP1*, 2. *sfGFP*, 3. vector) and two validation plasmids (Supplementary Figure 18 and 19). *ampR*: gene for ampicillin resistance, ±*D/B*: with or without DMSO and betaine, *eco47IR*: gene for a restriction enzyme (reduction of religations of the vector), *mRFP1*: monomeric red fluorescent protein 1, *ori*: origin of replication, *sfGFP*: superfolder green fluorescent protein, *T*_*m*_: melting temperature.

## MATERIALS AND METHODS

### Toy-model plasmid for the optimization experiments

All LCRs for the optimizations were performed with a toy-model plasmid made of seven fragments (Figure 1B), with a total length of 4918 bp. This plasmid consists of six inserts and the plasmid cloning vector CloneJet pJET1.2/blunt (2974 bp; Thermo Fisher Scientific, Massachusetts, USA). For the inserts, two genes of fluorescent proteins, *sfGFP, mRFP1*, were ordered at Addgene (www.addgene.org: pYTK001, pYTK090; 17) and split into three subparts each to increase size heterogeneity, so that the plasmid fragment length ranges from 79 bp to 2974 bp. Additionally, the same terminator *BBa B0015* was used in both fluorescent protein genes to further increase assembly complexity. A spacer sequence “S” of 37 bp was added to prevent the ligation of part 3 with part 7 (Figure 1B). For all *in silico* cloning, the software `Geneious` was utilized (v. 11.0.5, http://www.geneious.com, 18). A GenBank-file of the toy-model plasmid is available in the supplement and at www.gitlab.com/kabischlab.de/lcr-publication-synthetic-biology.

**Table 1.** Summary of the LCRs for the validation experiments using the toy-model plasmid and the validation plasmids 1 and 2 (Figure 1B, Supplementary Figure 18 and 19). Each plasmid was split into three and seven parts which were assembled by using the baseline and improved LCR protocol (Table 1). The LCRs were transformed in chemically competent cells by a heat-shock. For results indicated by an *, the LCRs were transformed by electroporation due to low or no colonies when chemical transformation was used. BO: bridging oligo, CFU: colony forming unit, DMSO: dimethyl sulfoxide, *T*_*m*_: melting temperature of a BO-half using the calculation of SantaLucia (20).

| Factor | Unit | LCR protocol |  |  | Protocol | Toy-model plasmid (TP) |  | Validation plasmid 1 (VP1) |  | Validation plasmid 2 (VP2) |  |
| --- | --- | --- | --- | --- | --- | --- | --- | --- | --- | --- | --- |
|  |  | Baseline | 75 °C | Improved |  | 3 parts | 7 parts | 3 parts | 7 parts | 3 parts | 7 parts |
| DMSO | %v/v | 8 | 8 | / |  |  |  | Efficiency (%) |  |  |  |
| Betaine | M | 0.45 | 0.45 | / | Baseline | 37±6 | 0±0* | 88±4 | 17±29* | 98±2 | 93±12 |
| Annealing temperature | °C | 55.0 | 55.0 | 66.0 | Improved | 47±6 | 17±7* | 96±1 | 98±3* | 95±0 | 93±3 |
| Target $T_m$ for each bridging oligo half | °C | 70.0 | 74.8 | 67.8 | | | | Total CFUs (per 3 µL LCR) | | | |
|  |  |  |  |  | Baseline | 250±46 | 6±3* | 125±8 | 1±1* | 365±63 | 12±2 |
|  |  |  |  |  | Improved | 559±33 | 30±14* | 493±19 | 323±70* | 2023±331 | 189±26 |

### Part amplification

Primers for the amplification (Eurofins Genomics, Ebersberg, Germany) were phosphorylated by the T4-polynucleotidekinase/-buffer (New England Biolabs, Ipswich, USA) prior to amplification *via* PCR (Q5^®^ High-Fidelity Polymerase, New England Biolabs, Ipswich, USA). Forward and reverse primers were phosphorylated separately in 50 µL total volume with 4 µM primer, 4 mM ATP and 10 U of T4-PNK for 1 h at 37 °C and 20 min at 65 °C for the denaturation. The low primer concentration was chosen because it is beneficial for the enzymatic phosphorylation. The 79 bp promoter of *mRFP1* (part 1, Figure 1B) was ordered as forward and reverse strand (lyophilized, salt-free). Both strands were phosphorylated separately as described for the amplification primers, followed by an annealing procedure to obtain double-stranded DNA (3 min at 95 °C and 70 cycles of 20 s with an incremental decrease of 1 °C). The backbone-part pJET1.2/blunt was PCR-amplified using a template containing a lacZ coding sequence (pJET1.2/blunt-*lacZ*). Choosing primers which omit *lacZ* during PCR allows to screen for plasmid-carryover through blue-white screening. The *lacZ* does not exist in the final sequence of the toy-model plasmid. Afterwards, all PCR products were DpnI-digested (60 min at 37 °C; inactivation: 20 min at 80 °C), purified (column-based NEB Monarch^®^PCR & DNA Cleanup Kit; New England Biolabs, Ipswich, USA) and the DNA quantity/quality was measured by photometry (Spectrophotometer UV5Nano, Mettler Toledo, Columbus, USA). Afterwards, the phosphorylated primers and DNA parts were stored at −20 °C. The sequences of the amplification primers are shown in Supplementary Table 2. The sequences of the toy-model parts are shown in Supplementary Table 1.

### Bridging oligo design

Bridging oligos were predicted according to the rules given in de Kok et al. (8): they were all orientated in forward direction and designed at specific concentrations of 10 mM Mg^2+^, 50 mM Na^+^, 3 nM plasmid parts, 30 nM BOs and 0 mM dNTPs. `Geneious` asks for a concentration of dNTPs but they are not utilized in the LCR. Though, the concentration has to be adjusted to 0 mM. All BOs were ordered lyophilized from Eurofins Genomics (Ebersberg, Germany) as salt-free custom DNA oligonucleotides. Quality was checked by Eurofins Genomics *via* matrix-assisted laser desorption ionization-time of flight (MALDI-TOF) or capillary electrophoresis (CE).

One BO-set was designed manually with the melting temperature tool of `Primer3` (19), which is distributed with the software suite `Geneious`. For the *T*_*m*_ calculation and salt correction, the nearest-neighbor algorithm and the corresponding salt correction by SantaLucia (20) were utilized to design a BO-set with melting temperatures of 70 °C for each half-BO. `Primer3` only accepts a single DNA concentration, despite the experiment using different concentrations of parts and BOs. As prompted by `Geneious`, only the BO concentration of 30 nM was inputted. This manual set is denoted by an “M”.

To investigate the crosstalk of BOs, additional sets were designed by minimizing or maximizing Δ*G*-dependent crosstalk between oligonucleotides while maintaining a *T*_*m*_ between 67 °C and 73 °C. Crosstalk is based on secondary structure and among others defined as the sum of all minimum free energies (MFEs) when cofolding each oligonucleotide of a BO-set with each oligonucleotide in that set and with itself. As a reference temperature for the crosstalk calculations, the annealing temperature of the multi-step LCR protocol was used (55 °C, 8). As with the manual set, the SantaLucia parameters were utilized for the *T*_*m*_ calculations. Additionally, the DNA part concentration was adjusted to 3 nM to match the experiment. The impact of DMSO and betaine were not considered for the calculations of the MFEs. Sets with minimized crosstalk are denoted by “L” for low crosstalk and sets with maximized crosstalk by a “H” for high crosstalk. All BO-sets, sequences and melting temperatures are provided in Supplementary Table 3 and are available at www.gitlab.com/kabischlab.de/lcr-publication-synthetic-biology. Further details on the BO design are given in the supplement (21, 22, 23, 24, 25, 26, 27).

### Ligase cycling reaction

For the assembly, the purified PCR products were mixed with supplements with the following concentrations: 1×-Ampligase^®^ ligase buffer, 0.5 mM NAD^+^, 3 nM of toy-plasmid parts 1-6, 0.45 M betaine and 8 %v*/*v DMSO. In contrast to de Kok et al. (8), the concentration of the vector pJET1.2/blunt (part 7) was reduced to 0.3 nM to achieve fewer religations of the vector. Positive effects of increasing the molar insert-to-vector ratio were already described for other cloning methods (28, 29) and were confirmed in preliminary LCRs (data not shown). LCRs using these experimental concentrations, i.e. the manually designed BOs and the cycling conditions described in the next paragraph, are referred to as baseline-conditions. A 10×-Ampligase^®^reaction buffer was self-made with the concentrations described by the manufacturer of the Ampligase^®^(Lucigen, Wisconsin, USA). NAD^+^ was supplied separately by using a self-prepared 10 mM stock solution. Bridging oligo sets were premixed in nuclease-free water with 1.5 µM of each BO, heated up for 10 min at 70 °C and cooled down on ice before adding them to the split master-mixes. Subsequently 7.5 U of Ampligase^®^ (Lucigen, Wisconsin, USA) was added. After mixing by inverting and centrifugation, each LCR was split typically in three (*n* = 3) reactions with a cycling-volume of 3 µL. LCR-quintuplets (*n* = 5) were used to investigate the impacts of crosstalk and utilizing or omitting DMSO/betaine, which are presented in Figure 2, 5 and 12. After the cycling, all LCRs were cooled down on ice to recondense evaporated liquid in the PCR-tubes, and centrifuged.

**Figure 2.**
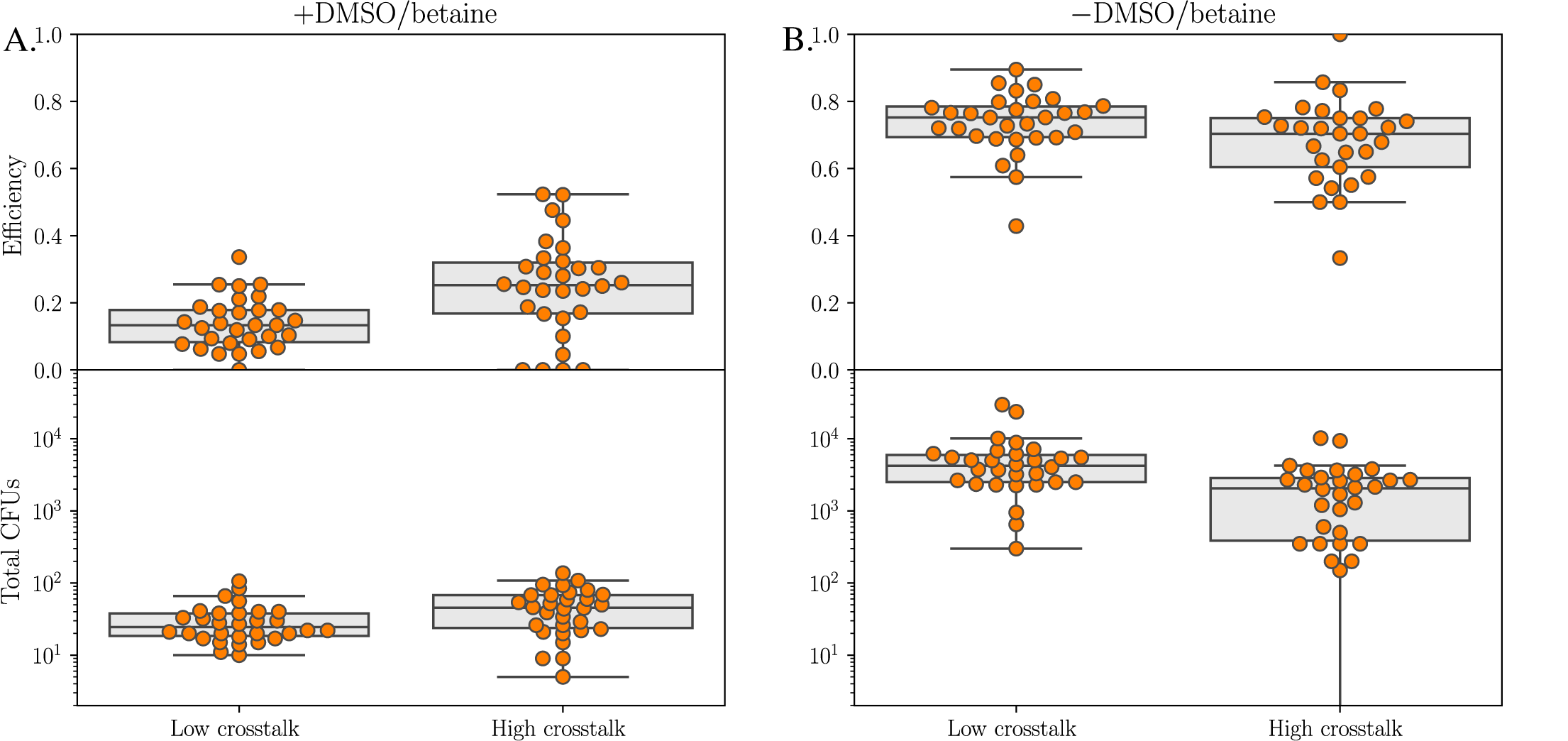
Overview of the seven-part LCR of the toy-model plasmid by utilizing six bridging oligo-sets (BO-sets) with low crosstalk and six BO-sets with high crosstalk. Each BO-set was used five times to assemble the toy-model plasmid (Figure 1B) by using the baseline-conditions with 8 %v*/*v DMSO/0.45 M betaine and without both detergents. For all LCRs, 3 µL were transformed by electroporation in 30 µL NEB^®^10-*β E. coli* cells. For more detailed results of each BO-set refer to Supplementary Figures 4 and 5. **A.** The baseline-conditions with DMSO and betaine resulted in low efficiencies and low amounts of colonies. No correlation between crosstalk and BO performance was found. **B.** LCRs without DMSO and betaine resulted in more colonies and higher efficiencies in comparison to the baseline-conditions. The sequences of all BO-sets are shown in Supplementary Table 3. BO: bridging oligo, CFU: colony forming unit, DMSO: dimethyl sulfoxide.

For the cycling, a DNA Engine Tetrad^®^2 thermal cycler, 96-well Alpha™Unit cycling block and low-profile PCR stripes (Bio-Rad Laboratories GmbH, Muenchen, Germany) were utilized. The speed of ramping used for all LCRs was 3 °C s^−1^. The cycling was initiated by a denaturation step at 94 °C for 2 min, followed by 25 cycles at 94 °C for 10 s, 55 °C for 30 s and 66 °C for 1 min. In contrast to the published protocols (8, 9), 25 cycles instead of 50 cycles were used due to the low total LCR volume of 3 µL. Afterwards, either 30 µL electro-competent or chemically-competent NEB^®^ 10-*β E. coli* cells (New England Biolabs, Ipswich, USA) were transformed as described in the following paragraph.

Electroporations were performed serially for each single LCR. For this, each LCR condition was transformed in a non-batch-wise manner to reduce workflow-derived influences, so that, e.g., each condition was transformed once before proceeding with the second replicates. 1. cells were mixed from a master-aliquot for each experiment with one LCR-replicate of one LCR condition, 2. suspension was transferred to a cuvette, 3. an electric pulse was applied (2.5 kV), 4. 450 µL SOC-medium were added, 5. the cell-suspension was transferred to a tube, 6. the tube was put for 1 h in a thermal cycler for recovery and 7. finally the suspension was plated on agar plates. After the recovery, an appropriate volume of the transformation mix was plated on Lysogeny Broth (Miller) plates containing 1 %m*/*v agar and 100 µgmL^−1^ ampicillin to get a total amount in the range of 10-200 CFUs per plate. The plates were grown for 15 h at 37 °C.

For chemical transformations, the experiments were performed in a parallel manner: 1. cells were mixed from a master-aliquot for each experiment with all replicates of all LCRs in a 96-well PCR-plate using a multi-channel pipette, 2. the suspensions were incubated on ice for 30 min, 3. the heat-shock of 30 sec was applied in a PCR-cycler by using 96-well PCR-plates, 4. the suspensions were placed on ice for 10 min, 5. 170 µL SOC-medium were added to all LCR-cell-mixes followed by recovery of 1 h and plating as described for the electroporations.

### Colony screening and plasmid sequencing

By transforming the LCRs into *E. coli*, the toy-model plasmid enables a fluorescence-based discrimination of correct (red and green-fluorescent CFUs) and misligated plasmids (red or green or non-fluorescent CFUs). To investigate the LCR assembly, the efficiencies of all LCRs were determined by observing the phenotype of ampicillin-resistant CFUs *via* fluorescence microscopy (microscope: Axio Vert.A1, Carl Zeiss Microscopy GmbH, Jena, Germany, 50×-magnification; LEDs for sfGFP/mRFP1: 470*/*540−580 nm). To calculate the LCR efficiency, the phenotypes of all CFUs per LCR were screened. For plates with more than 100 CFUs, 100 colonies were screened randomly by first picking the spot of interest macroscopically followed by the observation with the microscope. The efficiency was obtained by dividing the number of correct CFUs by the total number of microscopically observed CFUs. The phenotypes of the CFUs were determined by visual observation only. An example image of all phenotypes is shown in 3.

The amount of CFUs per 3 µL was obtained from the total CFUs per plate and the used plating volume and dilution.

To validate this fluorescence-based system, plasmids from 120 CFUs with different phenotypes were isolated with the Monarch^®^Plasmid Miniprep Kit (New England Biolabs, Ipswich, USA) and analyzed *via* Sanger-sequencing (Eurofins Genomics, Ebersberg, Germany) to correlate them with the genotypes.

### Graphical analysis

Unless specified otherwise, *T*_*m*_ calculations in the following sections were performed with the thermodynamic parameters and salt correction by SantaLucia (20) and the divalent salt conversion by von Ahsen et al. (30). The shown *T*_*m*_s are of half-BOs. Each half-BO is complementary to the DNA part that it is designed to attach to. A BO is the direct concatenation of two half-BOs. Dangling ends and coaxial stacking were not included in the calculation. Error bars are the standard deviation with Bessel’s correction. The average *T*_*m*_ of a BO-set is obtained from all half-BOs of the set.

### Validation experiments

For the validation experiments two additional plasmids-constructs were split in three and seven parts (Supplementary Figures 18 and 19) and were assembled using the baseline and two optimized conditions (Table 1) derived by the results of the toy-model plasmid. Both plasmids and splits are similar in size range in comparison to the toy-model plasmid with *sfGFP* and *mRFP1*. GenBank-files of the validation plasmids are available in the supplement and at www.gitlab.com/kabischlab.de/lcr-publication-synthetic-biology. In the validation plasmids the *lacZ* and two or one antibiotic resistance genes were split into subparts (Supplementary Figures 18 and 19). This enables a blue-white screening of correct assembled plasmids (blue colonies) and misassembled ones (white) by plating them on agar with the antibiotics and X-Gal/IPTG. For both plasmids, 3 µL per LCR were transformed in 30 µL NEB^®^ 10-*β E. coli* cells.

For the validation plasmid 1, the transformed cells was transferred to LB agar plates with 100 µgmL^−1^ spectinomycin after one hour of recovery followed by an overnight incubation for 16 h at 37 °C. 40 CFUs were transferred to agar plates containing spectinomycin, 50 µgmL^−1^ kanamycin, 40 µgmL^−1^ X-Gal and 200 nM IPTG by using sterile tooth picks to identify colonies with correct assembled and misassembled plasmids. After 16 h, all transferred colonies were checked for a blue/white phenotype and empty spots were regarded as kanamycin sensitive. Blue colonies represented correct assembled plasmids. White colonies and empty spots were treated as misassembled plasmids. To calculate the efficiency for each LCR the amount of blue colonies was divided by the amount of transferred CFUs. Ten plasmids of blue, spectinomycin and kanamycin resistant colonies were sequenced *via* Sanger sequencing. Further, four plasmids of white, spectinomycin and kanamycin resistant colonies were analyzed.

For the validation plasmid 2, the transformed cells were transferred to LB agar plates containing 100 µgmL^−1^ spectinomycin, 40 µgmL^−1^ X-Gal and 200 nM IPTG. After the incubation of 16 h at 37 °C, all colonies per plate were checked for a blue/white phenotype. For the calculation of each LCR efficiency, the amount of blue colonies were divided by the sum of blue and white colonies. As described for the validation plasmid 1, Sanger sequencing of plasmids from ten colonies with the correct phenotype “blue” and plasmids from four colonies with the wrong phenotype “white” was used to validate the results.

## RESULTS AND DISCUSSION

### Toy-Model plasmid offers robust system to investigate LCR assemblies

To simulate a challenging SynBio-construct for the investigations of the LCR, a toy-model plasmid was designed (Figure 1B). This plasmid consists of seven parts of varying size (79 bp to 2974 bp) and the same BioBrick terminator sequence *BBa B0015* in both fluorescent protein genes. To investigate the LCR different bridging oligo sets, experimental parameters, a toy-model plasmid and fluorescence-based analysis were used. For the analysis and optimization, ∼100 CFUs/LCR (if available) were screened *via* visual observation by using fluorescence microscopy to calculate the efficiency of each assembly reaction. The four observable phenotypes “non-fluorescent”, “green fluorescent”, “red fluorescent” and “green + red fluorescent” are demonstrated in Supplementary Figure 3. The amount of CFUs per 3 µL LCR was determined by macroscopic counting of all colonies per agar plate and extrapolation by using the dilution-factor. In total, 61 different experimental conditions were tested and more than 15000 colonies were screened by fluorescence-microscopy to obtain the assembly efficiencies. In contrast to the vector concentration of 3 nM in the literature (8, 9), the concentration was decreased to 0.3 nM to counteract religations.

No fluorescent CFUs were obtained in the control reactions without the ligase (Supplementary Figure 5). Additionally, blue-white screening of about 1000 non-fluorescent CFUs revealed no carry-over of the template used for the amplification of the vector pJET1.2/blunt. A carry-over of the templates pYTK001 or pYTK090 used for the amplification of *sfGFP* and *mRFP1* was not possible due to a change of the antibiotic resistance marker gene.

To validate the fluorescence-based readout, the observed phenotypes of 120 CFUs with different phenotypes were correlated with the corresponding plasmids *via* Sanger sequencing. The analyzed plasmids from 60 CFUs with a bicolored fluorescence (red and green) contained all seven DNA parts in the correct order/orientation when compared with the *in silico* sequence of the toy-plasmid. Sequencing results of 60 plasmids with a different phenotype (20 plasmids from only green fluorescent CFUs, 20 plasmids from only red fluorescent CFUs and 20 plasmids from non-fluorescent CFUs) indicated that they lacked at least one *sfGFP*-subpart or *mRFP1*-subpart (still red or green) or were religated vector (no fluorescence). The latter phenotype was also observed for plasmids with missing subparts of both fluorescent protein genes in preliminary experiments. Point-mutations in the LCR products were regarded as errors introduced by amplification primers, PCRs or *E. coli* and were not treated as LCR-misassemblies since they occurred outside of the ligation areas.

In our experiments, the misassembled plasmids from green fluorescent colonies lacked at least one subpart of the *mRFP1*. Interestingly, about 10 bp to 100 bp of both ends of the missing subparts were still existent. For the 20 analyzed CFUs with only the red fluorescent phenotype, the plasmids lacked the spacer sequence at the 3’-end of the *mRFP1* and the entire *sfGFP*. This suggests that *E. coli* can recognize the 129 bp *BBa B0015* terminator that is used for both *sfGFP* and *mRFP1* and delete it by *recA*-independent recombination (31, 32). Thus repetitive sequences in the LCR assembly cause misassemblies. Related to this, the BO that spans parts 6 and 7 can partly hybridize with the terminator in part 3 and may negatively influence the ligation. This suggests that duplicates at the end of DNA parts negatively influence LCR efficiency.

Another issue is related to the heterogeneous LCR-mixture, which contains the desired DNA fragments, debris of these fragments from PCRs, amplification primers and PCR templates (even if they are digested and purified). *E. coli* may circularize any linear DNA in the mixture. The subsequent transformation of *E. coli* enables a ring closure by endogenous mechanisms and the growth of CFUs containing the religated vector or plasmids with missing parts. This was observed in the control reaction without ligase in Supplementary Figure 5. Thus the carry-over of DNA in combination with the ability of *E. coli* to ligate linear DNA contributes both to the amount of misassembled plasmids and correct plasmids.

In summary, the fluorescence-based screening of colonies employed here is a valid and fast (manually: ~ 500 CFUs h^−1^) method to determine the assembly efficiencies and to investigate the influence of changing parameters in the LCR. A correlation of the phenotype and genotype is a useful tool without the need for next generation sequencing and offers an objective true-false readout by microscopy (as utilized for these studies) or photometric analysis of images (using e.g. `OpenCFU`, 33; `CellProfiler`, 34).

### Influence of DMSO, betaine and the *T*_*m*_ of bridging oligos on LCR assemblies

Thirteen different BO-sets, with sequences given in Supplementary Table 4, were used to assemble the toy-model plasmid made of seven fragments. Two separate experiments were performed with different conditions. The baseline LCR used 8 %v*/*v DMSO and 0.45 M betaine, whereas the crosstalk-increased LCR did not use any DMSO or betaine (Figure 2).

On average, the LCRs with both detergents revealed 3.6× lower efficiencies and 99× fewer total CFUs per 3 µL in comparison to assemblies without DMSO and betaine (*p*< 0.001, Figure 1C; raw data in Supplementary Figure 4). Further confirmation of these results can be seen in Supplementary Figures 12 and 14 in more consistent experiments using the same batches of DNA parts and competent cells. Graphical analysis of various predictors revealed no Δ*G*-related effects for assembling the toy-plasmid for both experimental setups (Figure 2, Supplementary Figure 6A+B; more predictors in Supplementary Figures 7, 8, 9 and 10). Guanine or cytosine at the 3’-end of BOs were not found to affect the LCR. Notably, the conditions without DMSO and betaine resulted in less red fluorescent phenotypes for all utilized bridging oligo sets (Supplementary Figure 4). Apparently, this phenotype is mainly related to *recA*-independent deletion by *E. coli*. Probably, the omission of both detergents is beneficial for assemblies with repetitive sequences.

For LCRs with DMSO and betaine, BO-related differences were detected, e.g., the LCR using BO-set L1 was less efficient than using H1 (Supplementary Figure 5A). Further analysis of the used BO-sets revealed an impact of the melting temperature despite the *T*_*m*_s of all sets being similar ranging from 68 °C to 72 °C (Supplementary Figure 6C). The set with a BO-*T*_*m*_ of 68 °C had an efficiency of 10% whereas the set with a BO-*T*_*m*_ of 72 °C showed an efficiency of 50%. This suggests that a melting temperature higher than 72 °C may result in even better assemblies for LCRs with DMSO and betaine. This is confirmed by recalculating the melting temperatures of the set of sequences designed to have a *T*_*m*_ of 70 °C by de Kok et al. (8), who used the SantaLucia parameters with the salt correction by Owczarzy et al. (35). We have found the average *T*_*m*_ of these half-BOs to be 72.2 °C when calculated using these parameters. When using the SantaLucia parameters and SantaLucia salt correction as has been done so far in this study, the *T*_*m*_ is found at 74.8 °C. The impact of different salt corrections is illustrated in Supplementary Figure 11. Overall, for LCRs with 8 %v*/*v DMSO and 0.45 M betaine, a BO target temperature above 70 °C was found to be beneficial.

Investigation of the `Primer3` source code used for the design of the manual BO-set and comparison with SantaLucia (20) revealed that the code expects both concentrations to be identical and to sum up to the input amount. Thus the prompted BO concentration of 30 nM corresponds to part and BO concentrations of 15 nM. Due to this, the manual BO-set had an average *T*_*m*_ of 71.5 °C (Supplementary Figure 6) when evaluated in full accordance with the SantaLucia formula instead of the targeted 70 °C.

An alternative approach to obtain higher assembly efficiencies without rising costs due to synthesizing longer oligonucleotides is to omit DMSO and betaine. This omission also aids automated liquid handling approaches because both detergents have unfavorable properties like extreme viscosities, hygroscopic characteristics and acting as surfactants. These new experimental conditions greatly improved both the efficiency and total number of CFUs (Supplementary Figure 5). Consistent with the results of the LCRs with DMSO/betaine, the melting temperature is the most influencing parameter. As a result of the omission of DMSO and betaine, BO-sets with lower *T*_*m*_s became favorable, yielding the greatest total amount of colonies at high efficiency (Supplementary Figure 6D). This behavior is also observed by comparing the total CFUs derived by utilizing the BO-sets L2, H1 and the manual one (Supplementary Figure 12). Similar to this seven-part toy-plasmid, another LCR-design with a *T*_*m*_-independent BO-design (20 bp for each BO-half) was published by Roth et al. (10) where six parts in a range of 79 bp to 2061 bp were assembled with high efficiencies. The *Taq*-ligase and a two-step LCR-protocol with annealing and ligation at the same temperature of 60 °C were utilized. This supports our findings that the omission of DMSO and betaine and to increase the annealing temperature can be helpful for ligation-based DNA assemblies.

An explanation for low efficiencies in the baseline LCR is related to the large difference between the annealing temperature of 55 °C and ligation temperature of 66 °C in combination with the utilization of DMSO and betaine. Both detergents reduce the energies required for strand separations (36, 37). Together with the experimental temperatures, this may result in an extensive reduction of template for the ligase by separating already hybridized BO/probe double strands. Bridging oligo-sets with lower *T*_*m*_s are theoretically more affected than sets with higher melting temperatures and should benefit from decreasing the temperature interval. Related to this, a reduction of the ligation temperature is expected to be advantageous. This was validated by using three BO-sets with different melting temperatures (Supplementary Figure 13) despite the ligation temperature of 60 °C being lower than the optimum temperature of the ligase (according to the manufacturer: 70 °C). Increasing the annealing temperature, which would also decrease the interval, was assumed to be disadvantageous due to accelerated BO/template-separation and was not investigated.

Several mechanisms are suspected to cause the total CFU decrease in LCRs with DMSO/betaine. First, lower LCR efficiencies result in fewer fully assembled plasmids and fewer colonies. The effects of DMSO and betaine were also investigated separately to prove these results (Supplementary Figure 14). Second, DMSO/betaine negatively influence the electroporation process. This can also be seen in Supplementary Figure 14, where an LCR without DMSO/betaine was mixed with both detergents before the electroporation and 3-4× fewer CFUs were obtained in comparison to DMSO/betaine-free controls. In contrast to the literature (8, 9), a lower volume-ratio of LCR and cells was used for the transformations (1:10; 2× higher concentration of DMSO/betaine). These conditions are not toxic for the *E. coli* strain NEB^®^ 10-*β* because chemical transformations of the same strain with the plasmid pUC19 revealed no negative effects of utilizing a combination of DMSO and betaine (Supplementary Figure 17). A negative impact of DMSO and a positive impact of betaine was observed. A strain-dependent influence of DMSO in chemical transformations was described in (38) and such an influence is also plausible for betaine and a combination of both reagents. To counteract the CFU-reducing effects of detergents, a 30 min dialysis using *aq. dest.* and a nitrocellulose-membrane after the LCR was found to increase the amount of colonies (data not shown).

The omission of DMSO/betaine was beneficial for the assembly of the seven-part split of the toy-model plasmid. To investigate optimal experimental temperatures for the new conditions without DMSO and betaine, the interaction of the annealing, ligation and BO-melting temperatures need to be considered.

### Increased annealing temperature is beneficial for LCRs without DMSO and betaine

To optimize the LCR without DMSO/betaine, several experiments were performed. Supplementary Figure 6D shows that the BO-sets with low *T*_*m*_s yield the most correct CFUs, which suggests that the BOs may bind too strongly to unwanted sites and be unavailable for ligation. To counteract this, the optimal annealing temperature was determined by performing a gradient-LCR, i.e. LCRs at different cycling conditions. For this, three new BO-sets were composed from the existing pool of BOs used in LCRs shown in Supplementary Figure 5 to obtain sets with average melting temperatures of 67.8 °C, 69.9 °C and 71.8 °C (sequences in Supplementary Table 4). In an annealing temperature range of 56.5 °C to 75.6 °C these sets were analyzed by chemically transforming the corresponding LCRs in chemically competent NEB^®^10-*β E. coli*. This resulted in roughly 100× lower transformation efficiency in comparison to the electrocompetent cells utilized in previous experiments.

For all sets, the efficiency was similar throughout the entire temperature range (Figure 3A). Consistent with previous results, the total amount of CFUs increased with lower BO-*T*_*m*_s (comparing the results shown in Supplementary Figure 6H and Figure 3B). Furthermore, all sets have a global CFU maximum in the annealing temperature range of ∼66-71 °C. This range contains the ligase optimum of 70 °C, which suggests positive effects of prolonging the total ligation time to 1.5 min (30 s annealing at 66-68 °C, 1 min ligation at 66 °C) in comparison to the baseline-condition of 30 s annealing at 55 °C and 1 min ligation at 66 °C.

**Figure 3.**
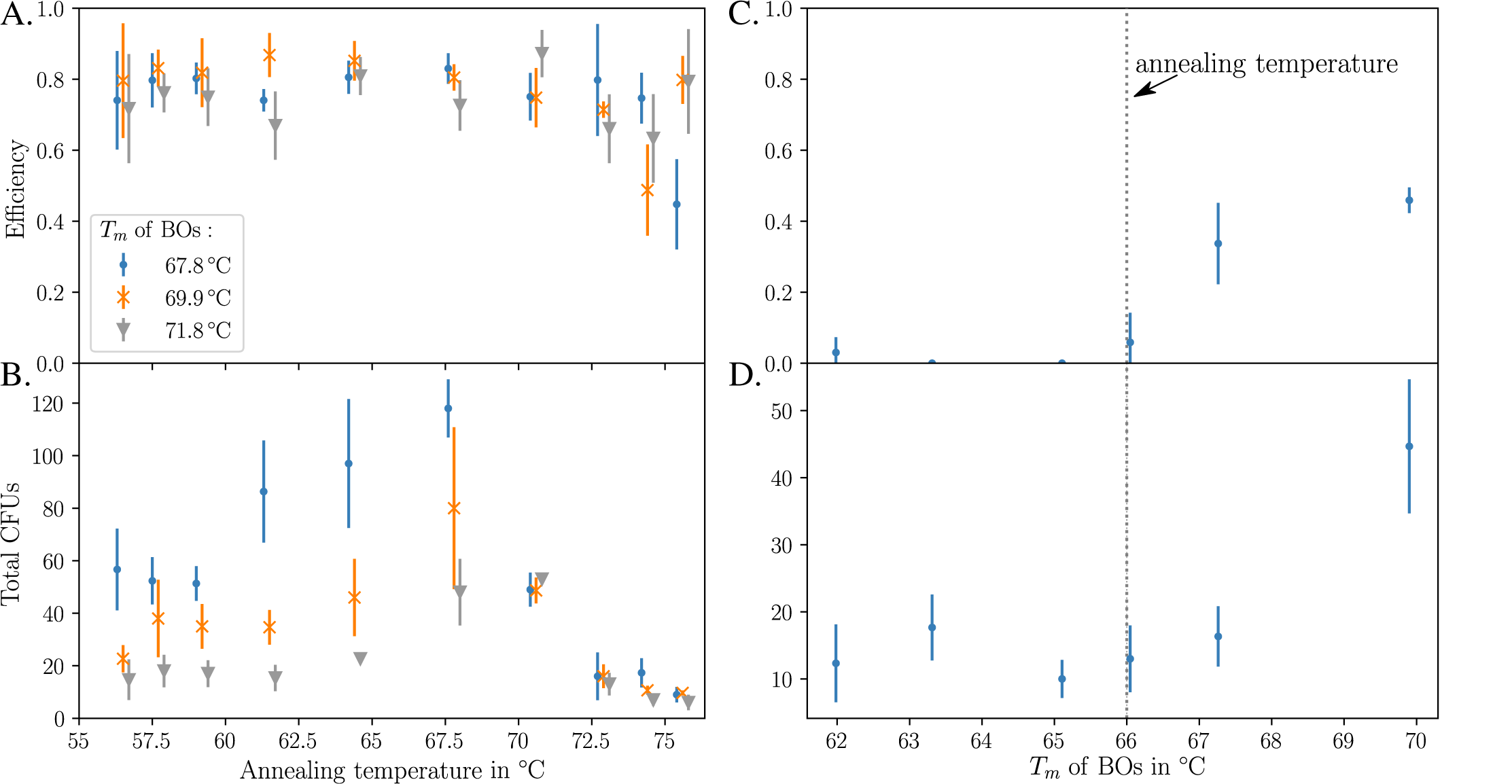
Adjustment of the annealing temperature and melting temperature of bridging oligo halves of DMSO/betaine-free LCR. The LCRs were performed as triplicates using the same DNA parts and chemically competent cells. The standard deviation for each LCR is indicated by error bars. **A.** For a better visibility the bars of the results were shifted with an offset of the annealing temperatures shown on the x-axis. Optimization of the annealing temperature of the seven-part plasmid *via* gradient-LCR. The temperature range for the annealing was 56.5 °C to 75.6 °C. Three BO-sets with different melting temperatures were used (67.8 °C, 69.9 °C and 71.8 °C). For the three BO-sets the LCR-efficiency was at a similar level. It started decreasing at an annealing temperature of more than 70.6 °C. **B.** As already observed, the total amount of colonies increased with a lower BO-*T*_*m*_. Overall, the LCRs using these BO-sets resulted in a maximum of colonies in a range of ∼66-71 °C with 2× more colonies (also shown in Supplementary Figure 15). The BO-set sequences of the three sets are shown in Supplementary Table 4. **C+D.** Based on the optimization shown in A+B, the annealing temperature of 66 °C was used to investigate the influence of BO-halves with lower *T*_*m*_s of 62.0 °C to 67.3 °C. As a reference, the BO-set “71.8 °C” from A+B was used. Further decrease of the BO-*T*_*m*_ did not improve the LCR at the optimized annealing temperature of 66 °C. In comparison to A+D, the LCR with the BO-set “69.9 °C” was less efficient due to loss of function by additional repeated freeze-thaw cycles of the DNA parts and BOs. The sequences of the BO-sets “62.0 °C” to “67.3 °C” are shown in Supplementary Table 5. BO: bridging oligo, CFU: colony forming unit, DMSO: dimethyl sulfoxide, *T*_*m*_: melting temperature of a BO-half.

In total, increasing the annealing temperature from 55 °C to 66 °C improved the CFU yield of every BO-set by a factor of two without a loss of efficiency. The BO-set “67.8 °C” performed better than the other sets. Supplementary Figure 16 shows that at the optimum annealing temperature of 67.8 °C, the BO-set “67.8 °C” also showed slightly greater efficiency than the other sets.

Lower BO melting temperatures may have additional benefits for LCR assemblies without DMSO and betaine. To validate this, new BO-sets with lower melting temperatures of 62 °C to 67.3 °C were designed and applied for ligations at 66 °C (Figure 3C+D, sequences in Supplementary Table 5). Surprisingly, a drop-off in efficiency and CFUs was observed for BO-sets with equal or lower melting temperatures than 66 °C. In comparison to the results shown in Figure 3A the efficiency of the 69.9 °C-set is 30% lower. This result is most likely related to a loss of function by additional freeze-thaw cycles or contamination of the DNA parts, BO-sets and/or supplements. Negative impacts of repeated freeze-thaw cycles on the DNA parts and BOs were already mentioned for the LCR (16) and for single-stranded oligonucleotides (39). Nevertheless, the total amount of CFUs was further optimized by using BOs with a *T*_*m*_ of 67.8 °C for each half and by increasing the annealing temperature in the range of the ligase optimum without decreasing the efficiency. Together with previous optimizations by performing the LCR without DMSO and betaine the assembly of the seven-part was highly improved. For the following experiments, we defined an improved LCR-protocol (Table 1).

### New LCR protocol improves the assembly of two different toy-model plasmid splits

In order validate the positive effects of omitting DMSO/betaine and to use a higher annealing temperature of 66 °C in DMSO/betaine free conditions, the sequence of the seven-part plasmid was used for a three-part split consisting of *mRFP1* as part 1, *sfGFP* as part 2 and pJET1.2/blunt as part 3. The BOs for this assembly were newly ordered and new aliquots of all LCR supplements were utilized. This plasmid was assembled by using the baseline and improved LCR conditions (Table 1). For the baseline condition, the LCR was performed using the manual BO-set with a *T*_*m*_ of 71.4 °C for each half, 8 %v*/*v DMSO, 0.45 M betaine and the annealing at 55 °C. For the improved LCR condition, the BO-set had a *T*_*m*_ of 67.8 °C for each half (set was already used for the gradient-LCR in Figure 3), no DMSO/betaine were used and the annealing was at 66 °C. The sequences of both BO-sets are shown in Supplementary Table 10. In general, the three-part plasmid was built by the two different LCR-protocols with no differences in the total plasmid size (4918 bp), sequence and BOs in comparison to the seven-part version. Further, the seven-part split was also assembled by using both protocols.

For both splits of the toy-model, the omission of DMSO and betaine was beneficial (plasmid “TP” in Figure 4 and Table 1) although the differences are lower in comparison to the optimization experiments. The low efficiencies of the seven-part split without DMSO/betaine in comparison to the high efficiencies obtained in the LCRs shown in Figure 2 are likely related to the repeated freeze-thaw cycles of the DNA parts. Further, the three-part LCR split was transformed in chemically and electrocompetent cells (Supplementary Figure 20). The results indicate that the LCR efficiencies are independent of the transformation method. Interestingly, the amount of colonies are similar for the chemical transformation and electroporation although the pUC19-transformation efficiency of the electrocompetent cells was ∼100× higher. We assume that the negative effects of DMSO/betaine in the electroporation and the higher transformation efficiency (Supplementary Figure 14) compensate each other. For the LCRs without DMSO and betaine the amount of CFUs was higher when the electroporation was used.

**Figure 4.**
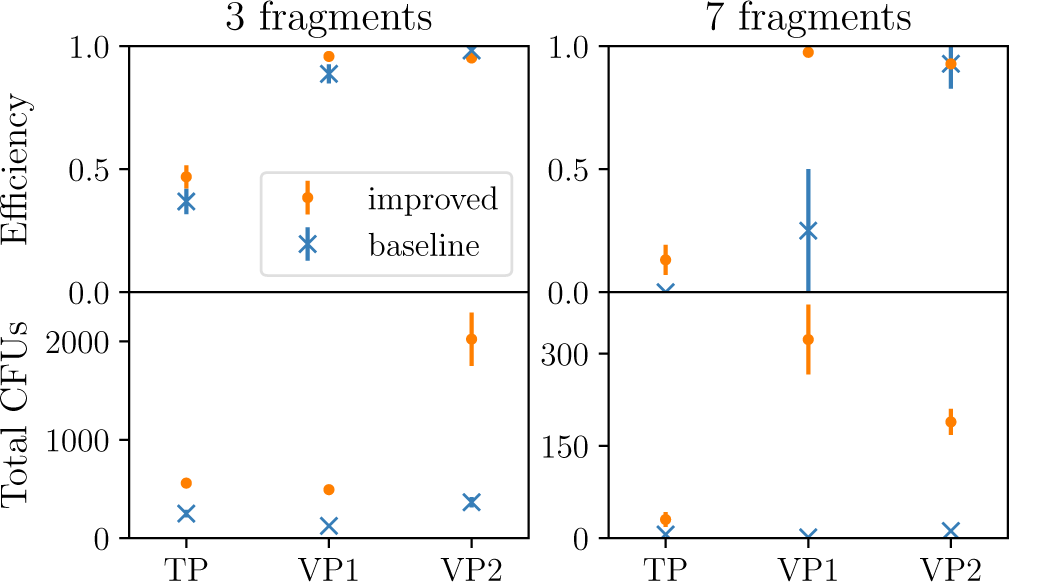
Comparison of the baseline and the improved LCR protocol for assembling the toy-model plasmid (“TP”) and the validation plasmids 1 (“VP1”) and 2 (“VP2”). The improved protocol increased the total amount of (correct) colonies for all assemblies. The efficiencies were also improved, although the LCRs of the three-part splits were similar for both protocols. LCRs were performed as triplicates and were transformed by chemical transformation. For the seven-part split LCRs of the toy-model plasmid “TP” and validation plasmid “VP1” the LCRs were transformed by electroporation due to low or no colonies when chemical transformation was used. CFU: colony forming unit.

### Validation of the improved protocol

To further verify the improved LCR protocol, we designed two validation plasmids based on the reporter genes *lacZ* and one or two antibiotic resistance genes (Supplementary Figures 18 and 19). For both plasmids a three-part and seven-part split were designed and three protocols were applied: the baseline, improved and a third protocol using BO-sets with a higher target melting temperature of 74.8 °C. According to the results shown in Supplementary Figures 6C and 11A, BOs with a higher *T*_*m*_ may improve the efficiency.

As shown by the optimizations for the toy-model plasmid, the improved LCR protocol is beneficial for the assembly of both splits of the validation plasmids “VP1” and “VP2” (Figure 4 and Table 1). Due to high efficiencies when the baseline protocol was applied, the effect of an increased efficiency by the improved protocol was only seen for the seven-part split of the toy-model plasmid “TP” and validation plasmid “VP1”. Nevertheless, the optimized protocol without DMSO/betaine increased the total amount of CFUs in comparison to the baseline protocol. According to the results of the third protocol, higher BO melting temperatures improved the LCR with DMSO and betaine but with a low impact (Supplementary Figure 21 and 22). In addition, longer BOs are used in this protocol which are typically more expensive. For automated assemblies the usage of DMSO and betaine would also be disadvantageous.

For both seven-part splits, Sanger sequencing of ten plasmids from CFUs with the correct phenotypes “blue” and resistance to the antibiotic(s) was performed for each validation plasmid. No false-positives were revealed. Control reactions without BOs and Ampligase^®^were performed for all splits and revealed no colonies. Additionally, four plasmids from colonies with the wrong phenotype “white” were analyzed for each validation plasmid. For the validation plasmid 1, the sequences had point-mutations in the 80 bp part of the *lacZ*. This part was ordered as oligonucleotides. This suggests that the mutations are due to the oligonucleotide synthesis. For the validation plasmid 2, three out of four plasmids had a point-mutation in the same *lacZ* part mentioned for the validation plasmid 1. One plasmid lacked the *lacZ*-parts 1 and 2 (parts 4 and 5 of the seven-part split shown in Supplementary Figure 18). Nevertheless, the validation experiments support the improved LCR protocol because the false-negative rate only affects the efficiencies and not the total amount of CFUs.

Besides general factors like the genetic context, total plasmid size, amount of parts, purification grade and freeze-thaw-cycles, the successful optimization of the LCR depends on BOs with melting temperatures which fit the kinetic conditions in the reaction. According to this study, the assembly of two splits of three plasmids showed a negative impact when DMSO and betaine are supplied (Figures 2 and 4) although it is considered to be beneficial in the literature (8).

## CONCLUSION

To our knowledge, the presented assembly of a plasmid with *sfGFP* and *mRFP1* is the first documented experimental LCR-design that includes a direct and fast readout to investigate the influence of various plasmid designs and experimental conditions. The utilized toy-model plasmid in combination with the fluorescence-based analysis enables a robust and easy-to-adapt *in vivo* system to get valuable insights into the LCR and is also adaptable for investigations of other assembly techniques.

Based on this workflow, the impact of intramolecular and intermolecular crosstalk between BOs is assumed to be negligible for the assembly of a seven-part toy-plasmid, whereas a strong interdependence in regards to the addition of DMSO/betaine, the BO-*T*_*m*_, the annealing temperature and ligation temperature was observed. Those findings were validated by assembling two validation plasmids by comparing the baseline conditions with an improved LCR protocol without DMSO/betaine, a target *T*_*m*_ of 67.8 °C for each BO-half and an annealing temperature of 66 °C. Related to this, it is of crucial importance to be consistent in the choice of the algorithms for the *T*_*m*_ calculation. Sets with a melting temperature of 67.8 °C when using the formula of SantaLucia (20) are recommended. A melting temperature of 65.2 °C is recommended when using the algorithms of SantaLucia (20) and the salt correction of Owcarzy et al. (35). Using guanine or cytosine at the 3’-end of the BOs was not found to be beneficial.

In total, we used a new toy-model plasmid to investigate an improved LCR protocol without DMSO/betaine and successfully applied it for the assembly of two validation plasmids. We conclude that our improved LCR protocol will help to achieve more efficient assemblies and to enable a simpler way for automated LCR assemblies by omitting DMSO/betaine and adjustments of the thermal conditions. To ensure reproducible conditions we provide a step-by-step LCR-protocol. You can download it from the digital supplement. We also offer a beta-version of a `Geneious`-plugin to design LCR-constructs and bridging oligos with the improved parameters at www.gitlab.com/kabischlab.de/lcr-publication-synthetic-biology. In the future, we will provide a web-application and a stand-alone tool for the automated design of LCR-constructs and bridging oligos. To accelerate the screening process flow cytometry could applied to use this validated method for future applications to e.g., investigate more rules for the LCR or other assembly methods. Another interesting approach would include the combination of *in vitro* studies using cell-free systems (40) and the toy-model plasmid.

## ACKNOWLEDGEMENTS

We thank Carolin Dombrowsky for her technical support to perform the transformations of *E. coli* and the fluorescence-based screening. The authors also acknowledge the scientific discussions and advices with members of the LOEWE CompuGene-group. We further thank Brigitte Held and Dunja Sehn for their organizational support and Alexander Rapp for providing the fluorescence microscope and technical assistance.

## SUPPLEMENTARY DATA

Supplementary data in a pdf file. A step-by-step LCR protocol with the improved conditions is also provided. All GenBank-files, bridging oligos sequences and the `Geneious`-plugin for the design of bridging oligos are available at www.gitlab.com/kabischlab.de/lcr-publication-synthetic-biology. The GenBank-files also contain the manually designed bridging oligos, amplification primers and sequencing primers.

## SUPPLEMENT

### Bridging oligo optimization details

Crosstalk-optimized sets of bridging oligonucleotides (BOs) were obtained by quantifying the crosstalk with a score derived from secondary structure and then sampling with simulated annealing from the search space.

### Search space

The oligo search space is restricted because each half-oligo (henceforth called primer) must be complementary to the end of a DNA part and the melting temperature between them must be slightly above the experimental ligation temperature. While perfect complementarity is technically not required because thermodynamic data exists for mismatches, melting temperature calculations are too inaccurate for such an approach. *In silico*, changing the set of thermodynamic parameters often has a bigger impact on the melting temperature than inserting a mismatch, and so sensible design of non-complementary oligos is not feasible.

The oligo candidates were created by concatenating two primers, each of which is complementary to one end of one DNA part. For each end of each DNA part, a list of primers was created by taking substrings of varying lengths, where one end of the substring had to coincide with an end of the part. Taking the complementary was not necessary because the experiments contained double-stranded DNA. Each such primer had to satisfy the conditions that its melting temperature to the part is in the range 67°C ≤ *T*_*m*_ ≤ 73°C. The list of BOs for a pair of DNA parts was obtained by taking the Cartesian product of the set of primers for the 3’ end of one part with the set of primers of the 5’ end of the next part, and finally concatenating the individual elements of the product. This yielded a list *q*_*i*_ of all possible BOs for every bridging site 1 ≤ *i* ≤ *N*, which were used for the subsequent calculations.

### Scoring

The score to be minimized was defined in terms of the Hamiltonian 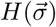 with 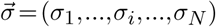 and *σ*_*i*_ ∈ {*q*_1_,…,*q*_*M*_} being the BO at site *i*. We implemented three different choices for 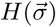, all of which are closely related to secondary structure and the minimum free energy (MFE). Throughout, we have used the dna mathews2004.par file that ships with the `ViennaRNA`-package (25).

1. Sum of the MFEs of pairwise BO-binding. For this, the Hamiltonian takes the form

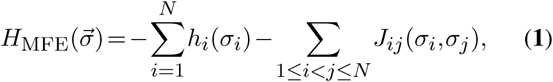

with *h*_*i*_(*σ*_*i*_) being the MFE of cofolding between two *σ*_*i*_ and *J*_*ij*_ (*σ*_*i*_,*σ*_*j*_) being the MFE of cofolding between *σ*_*i*_ and *σ*_*j*_. The cofolding MFE of every pairwise BO combination is obtained with an implementation of the Zuker algorithm (21). This Hamiltonian choice makes intuitively sense because higher MFEs correspond to less interaction between oligos.
2. Consideration of melting temperature *T*_*m*_ between pairwise BO-binding. The Hamiltonian 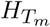 takes the same form as in (**1**) with *h*_*i*_(*σ*_*i*_) now being the negative *T*_*m*_ between two *σ*_*i*_ and *J*_*ij*_ (*σ*_*i*_,*σ*_*j*_) being the negative *T*_*m*_ of cofolding between *σ*_*i*_ and *σ*_*j*_. This negative sign takes into account that the (positive) *T*_*m*_ of a pairwise BO-binding should be as low as possible. The melting temperatures are computed by an MFE calculation with a modified traceback scheme. Instead of a secondary structure, the traceback returns entropy change Δ*S* and enthalpy change Δ*H*. The melting temperature follows as

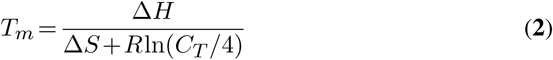

with the gas constant *R* and the BO concentration *C*_*T*_.
3. Consideration of unbound (free) BO concentrations. For the calculation of free BO concentrations we use a similar approach as described in Bernhart et al. (22), where the authors were able to calculate the concentrations of bound and unbound RNA-strands by obtaining equilibrium constants from the McCaskill algorithm (23) and subsequently solving a quadratic equation system. Within their calculations, they considered a two-species model. Here, we adapt their code to an arbitrary number of species. According to the literature (22, 24), the mass action equilibrium defines the concentration for two cofolded BOs [*σ*_*i*_,*σ*_*j*_] as

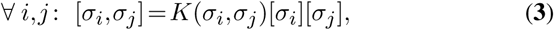

where [*σ*_*i*_],[*σ*_*j*_] are the monomer concentrations. The equilibrium constant *K*(*σ*_*i*_,*σ*_*j*_)= *Z*(*σ*_*i*_,*σ*_*j*_)*/*(*Z*(*σ*_*i*_) · *Z*(*σ*_*j*_)) is defined by the partition function of the bound state *Z*(*σ*_*i*_,*σ*_*j*_) and the free states *Z*(*σ*_*i*_),*Z*(*σ*_*j*_). The partition functions represent the thermodynamic configurations of the corresponding state (bound or unbound) and are obtained from the `ViennaRNA`-package. In analogy to Bernhart et al. (22), we introduce the total particle concentration 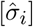 for a BO as

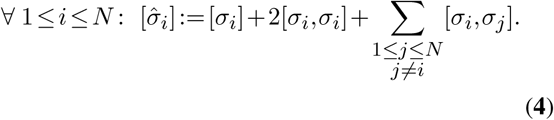 For a specific sequence 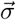, eqs. (**3**) and (**4**) yield the following quadratic system of equations due to mass conservation (22):

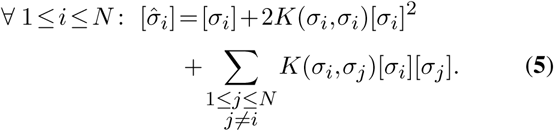 The Jacobian follows as

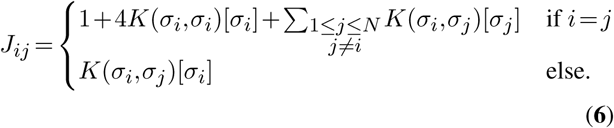 In eqs. (**5**), *K*(*σ*_*i*_,*σ*_*j*_) and 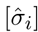 are calculated or part of the user input, so the system of equations can be solved for [*σ*_*i*_] to obtain the equilibrium concentrations 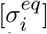. Within our implementation, we use the root finder of the `GNU Scientific Library` (26). Finally, we define our Hamiltonian by

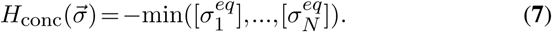 This Hamiltonian choice ensures a sufficient amount of the BO with the minimum equilibrium concentration.

### Sampling

We used the simulated annealing algorithm (27), where we define

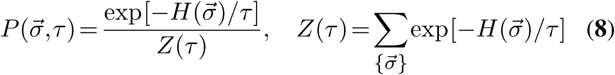

with the artificial temperature *τ*. A single mutation step consists of randomly picking *i* ∈{1,…,*N*} and 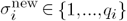 and changing 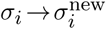.

For each Hamiltonian, three runs of simulated annealing were performed to obtain three sets with minimized crosstalk. Out of these three, the two sets with the greatest edit distance to each other were selected. The same procedure with opposite-sign Hamiltonians was done for a optimization towards the worst case. In total, we obtained six sets with minimized crosstalk and six sets with maximized crosstalk. Supplementary Figures 1 and 2 however show that crosstalk turned out to be a much worse indicator of performance than the melting temperatures between BO and DNA part.

**Supplementary Figure 1.**
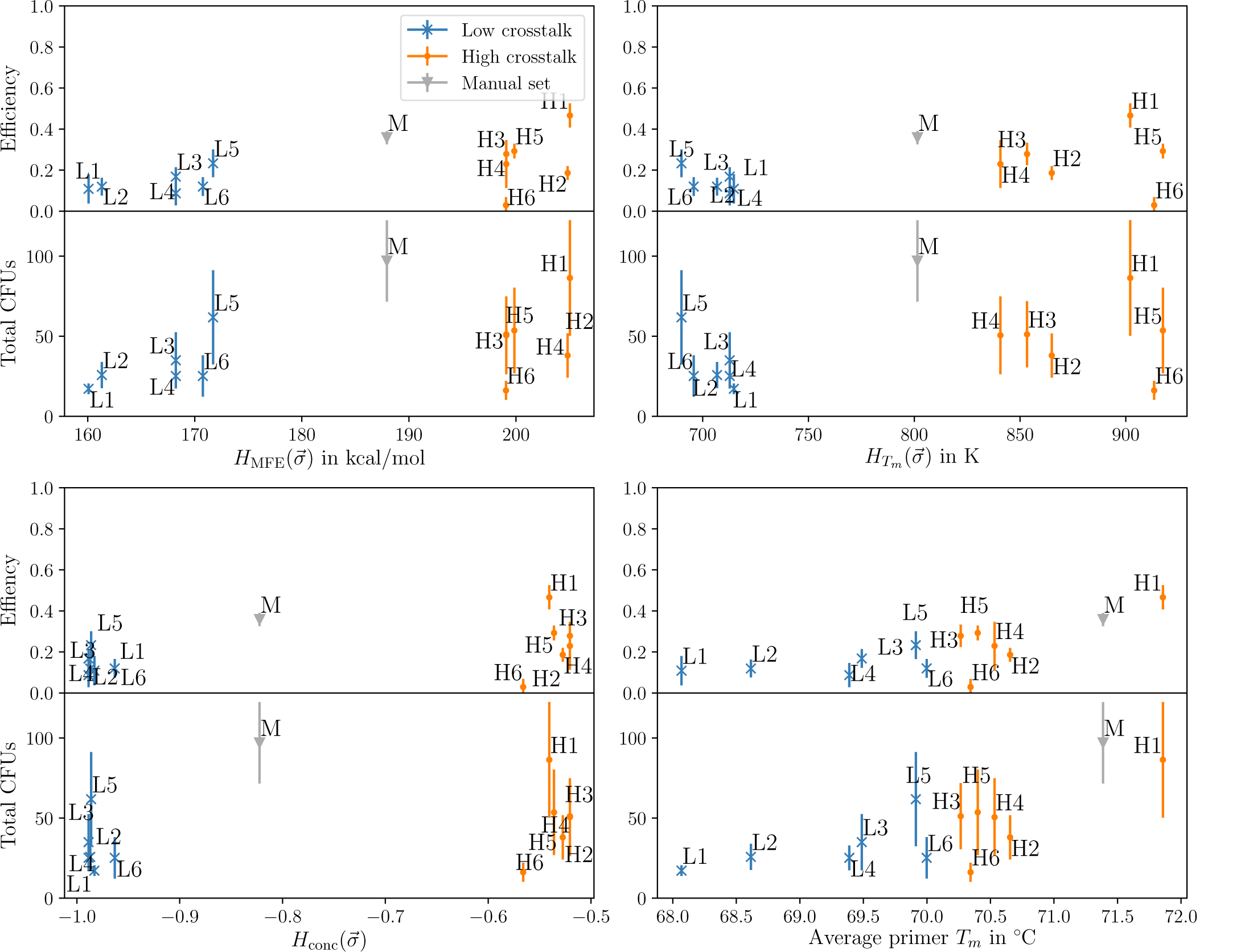
Efficiency and total number of CFUs depending on crosstalk and melting temperatures of BO sets with crosstalk optimized towards high (H) or low (L) values plus one manually designed set (M). The data is from the LCRs <u>with</u> 8 %v*/*v DMSO and 0.45 M betaine of the seven-part toy-plasmid. All LCRs were performed as quintuplets. The optimization criterion for each set can be seen in the crosstalk plots, e.g. sets L1 and L2 appear at the very left of the MFE crosstalk plot because they were optimized for low MFE crosstalk. High-crosstalk sets perform better than low-crosstalk sets because greater crosstalk is usually accompanied by greater melting temperatures; this counteracts the effect of DMSO and betaine, which lower the melting temperature. Ultimately, secondary structure was inhibited so much by DMSO and betaine that crosstalk could not occur and the BOs could barely attach to their complementary DNA parts. Thus the melting temperatures were a better criterion for performance than crosstalk.

**Supplementary Figure 2.**
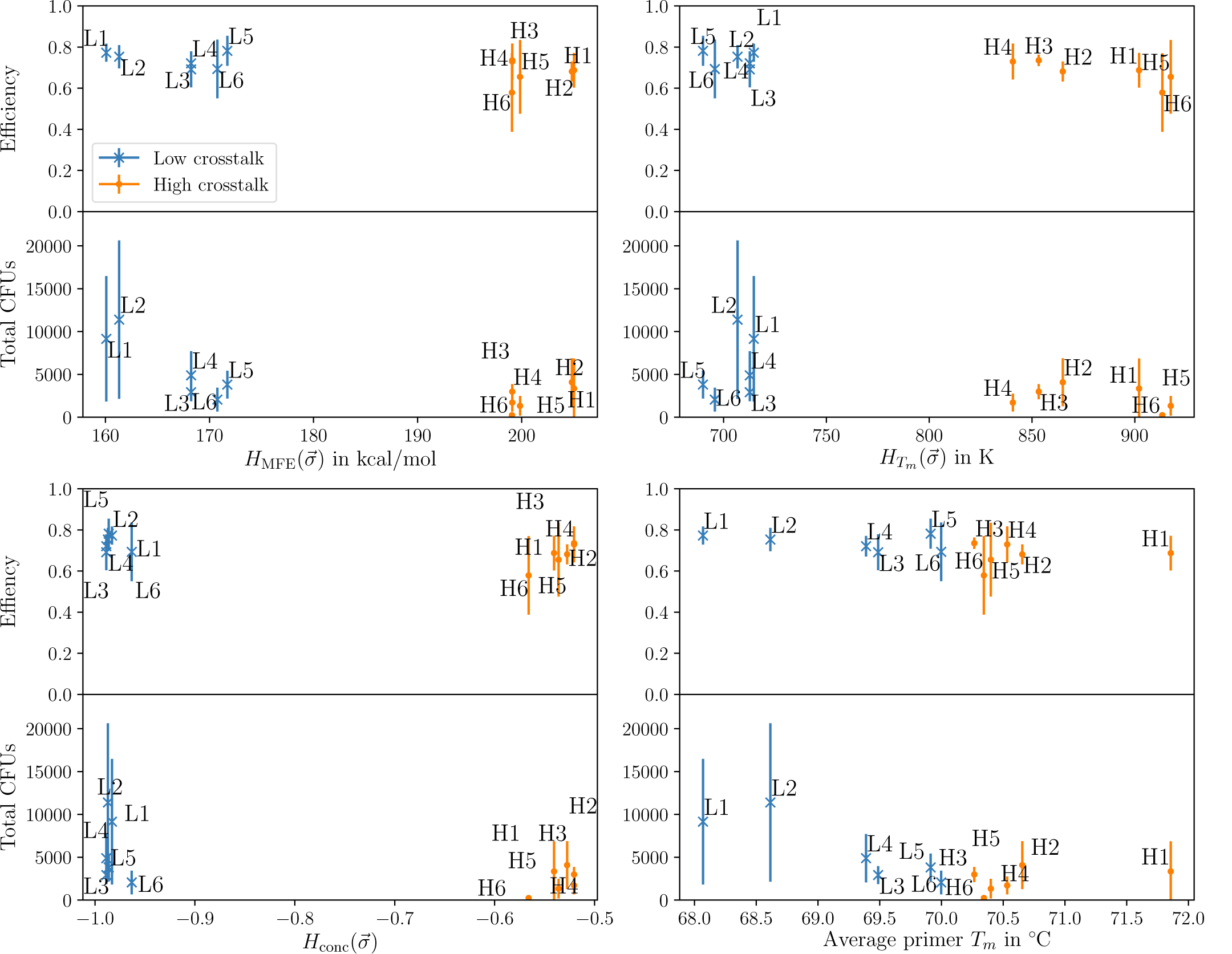
Efficiency and total number of CFUs depending on crosstalk and melting temperatures. The data is from the LCRs <u>without</u> DMSO and betaine of the seven-part toy-plasmid. All LCRs were performed as quintuplets. MFE-minimized sets perform better than the remaining sets, but the reason is likely their low melting temperature and not their low crosstalk. Their crosstalk evaluated by 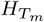 and *H*_conc_ does not differ from the other low-crosstalk sets, which further suggests that the melting temperature is the main reason for the performance.

### Phenotypes of the colonies obtained by the LCRs with the toy-model plasmid

**Supplementary Figure 3.**
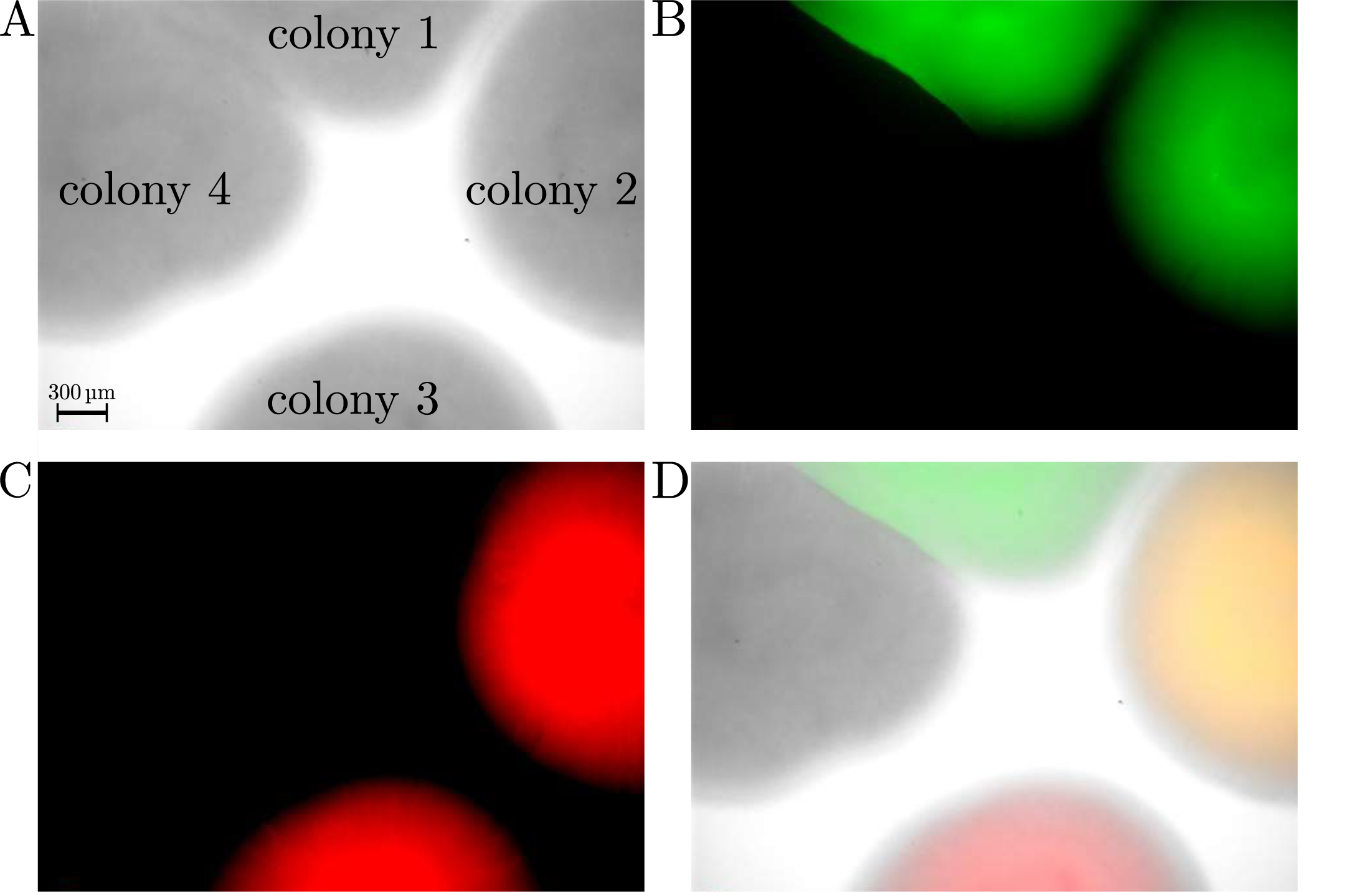
The four observed phenotypes of CFUs after the transformation of the LCRs of the toy-model plasmid. The “colony 2” with the red and green phenotype contained the correct sequence of the toy-model plasmid. The magnification of the images is 40×. **A**. Brightfield image of four colonies. **B**. Image of the colonies shown in A with (colonies 1 and 2) and without sfGFP. **C**. Image of colonies shown in A and B with and without mRFP1 (colonies 2 and 3). **D**. Overlay of the images A-C. CFU: colony forming unit, mRFP1: monomeric red fluorescent protein 1, sfGFP: superfolder green fluorescent protein.

### Optimization experiments with the toy-model plasmid

**Supplementary Figure 4.**
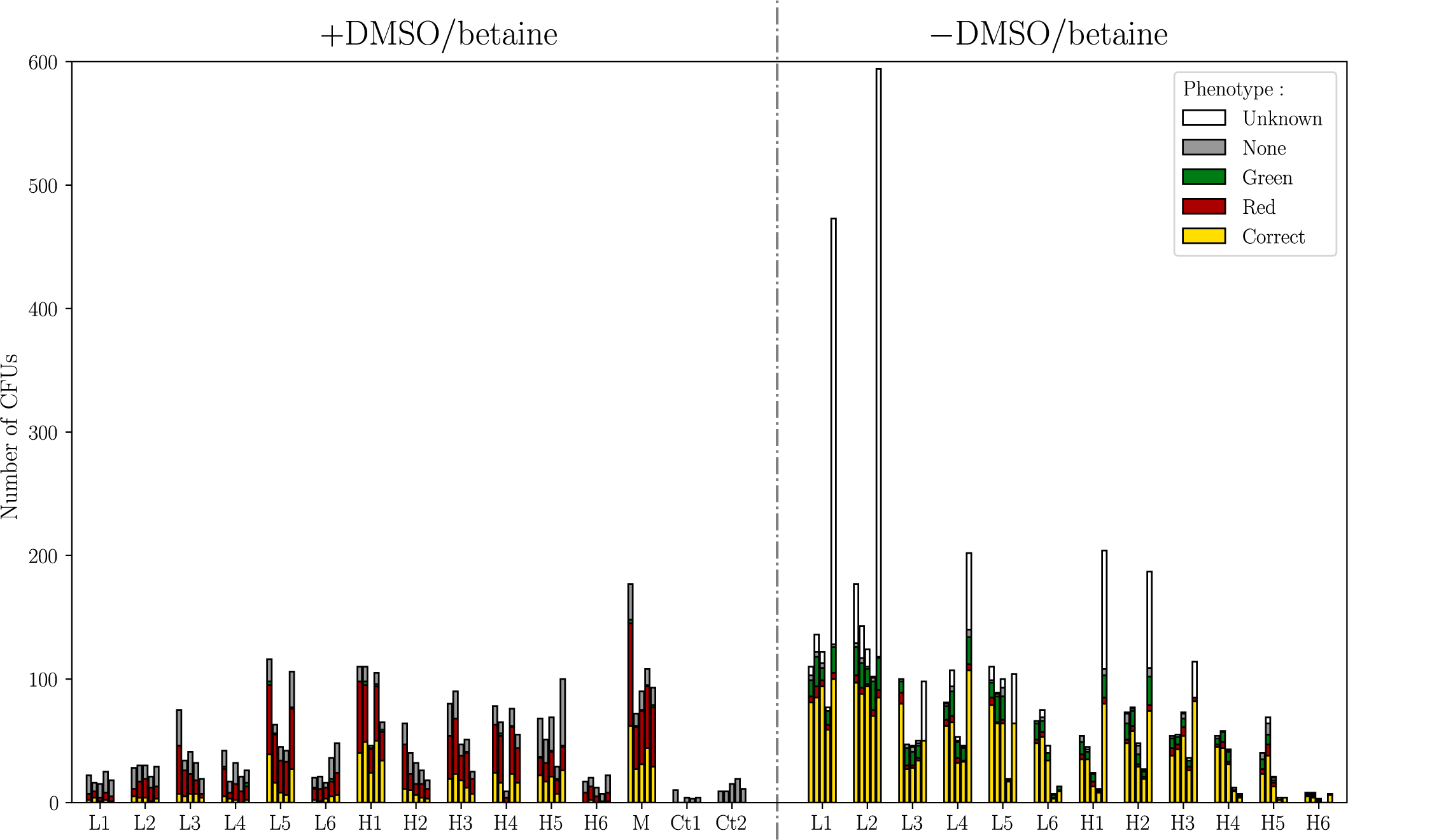
Overview of raw data of the experiments to investigate and calculate the influence/omission of DMSO and betaine of the seven-part toy-plasmid (Supplementary Figure 5). All LCRs were performed as quintuplets. Bridging oligo sets with minimized (L1-L6) crosstalk, maximized (H1-H6) crosstalk and the manually designed set (M) are presented. The negative control reactions with BOs without ligase (Ct1) and without BOs and ligase (Ct2) resulted in no fluorescent colonies and were only performed for the LCRs with DMSO/betaine. For the LCRs without DMSO and betaine the amount of CFUs presented here were corrected by multiplying them with the correction factor of 50. All CFUs were counted, but only the phenotypes of about 100 CFUs per LCR were determined. White bars indicate CFUs where the phenotype was not determined. All BO sequences are shown in Supplementary Table 3. BO: bridging oligo, CFU: colony forming unit, DMSO: dimethyl sulfoxide, *T*_*m*_: melting temperature of a BO-half.

**Supplementary Figure 5.**
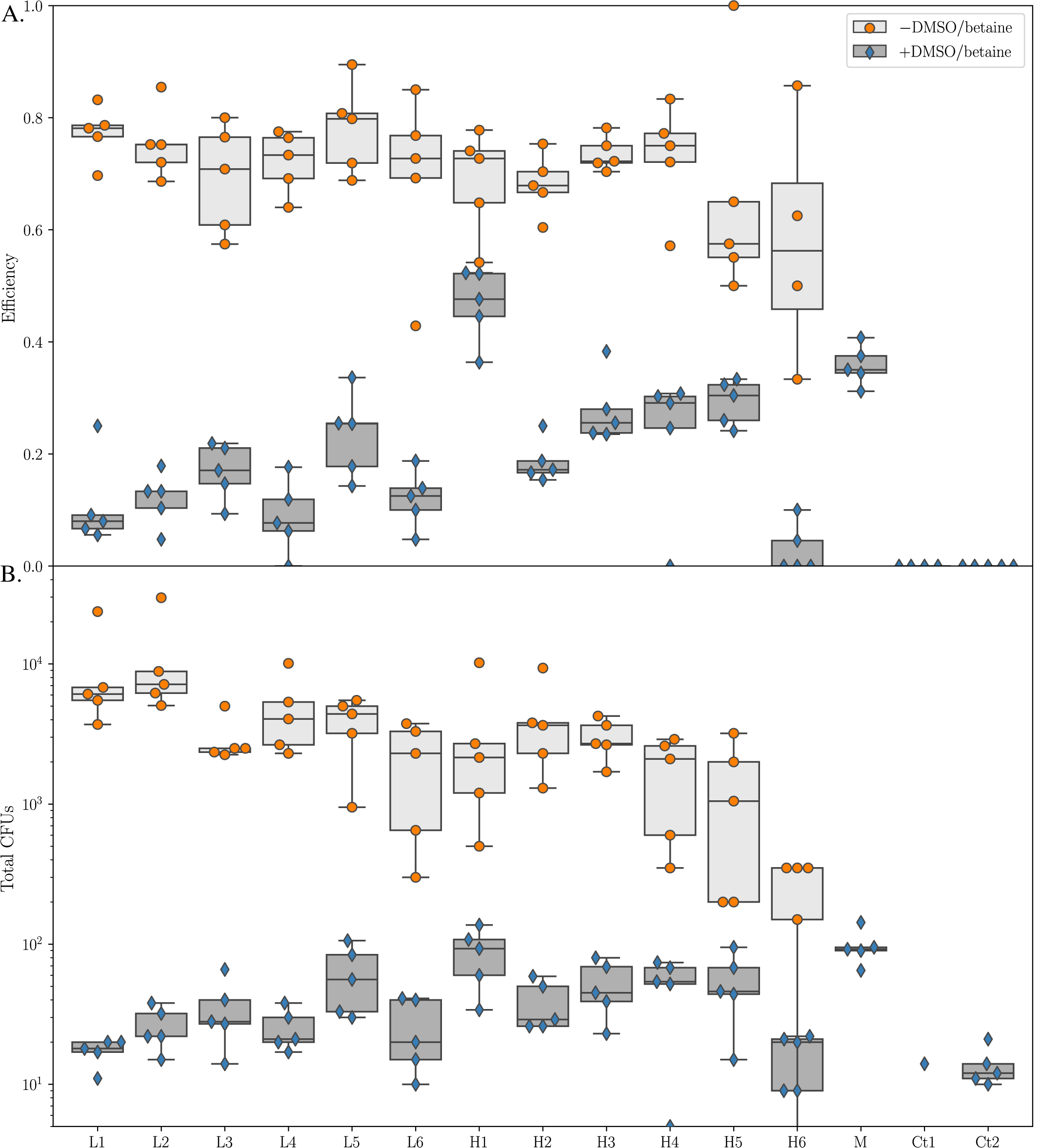
LCRs of a seven-part toy-plasmid by utilizing bridging oligo-sets (BO-sets) with high crosstalk (sets H1-H6) and low crosstalk (sets L1-L6). Each LCR was performed as a quintuplet. The standard deviation for each LCR is indicated by error bars. In addition to these twelve sets, a 13th set (“manual” set M) was utilized for the baseline LCR (8 %v*/*v DMSO and 0.45 M betaine). **A.** All LCRs without DMSO/betaine resulted in higher efficiencies in comparison to the LCRs with both detergents. No correlation between crosstalk and BO performance was found. Additionally, BO-set dependent differences were found for the baseline LCRs, e.g., BO-set H1 resulted in a higher efficiency than BO-set L1. The negative control reactions with BOs without ligase (Ct1) and without BOs and ligase (Ct2) resulted in no fluorescent colonies. **B.** LCRs without DMSO/betaine resulted in more colonies in comparison to the baseline-conditions. The raw data of the LCRs presented here are shown in Supplementary Figure 4. The sequences of all BO-sets are shown in Supplementary Table 3. BO: bridging oligo, CFU: colony forming unit, DMSO: dimethyl sulfoxide.

**Supplementary Figure 6.**
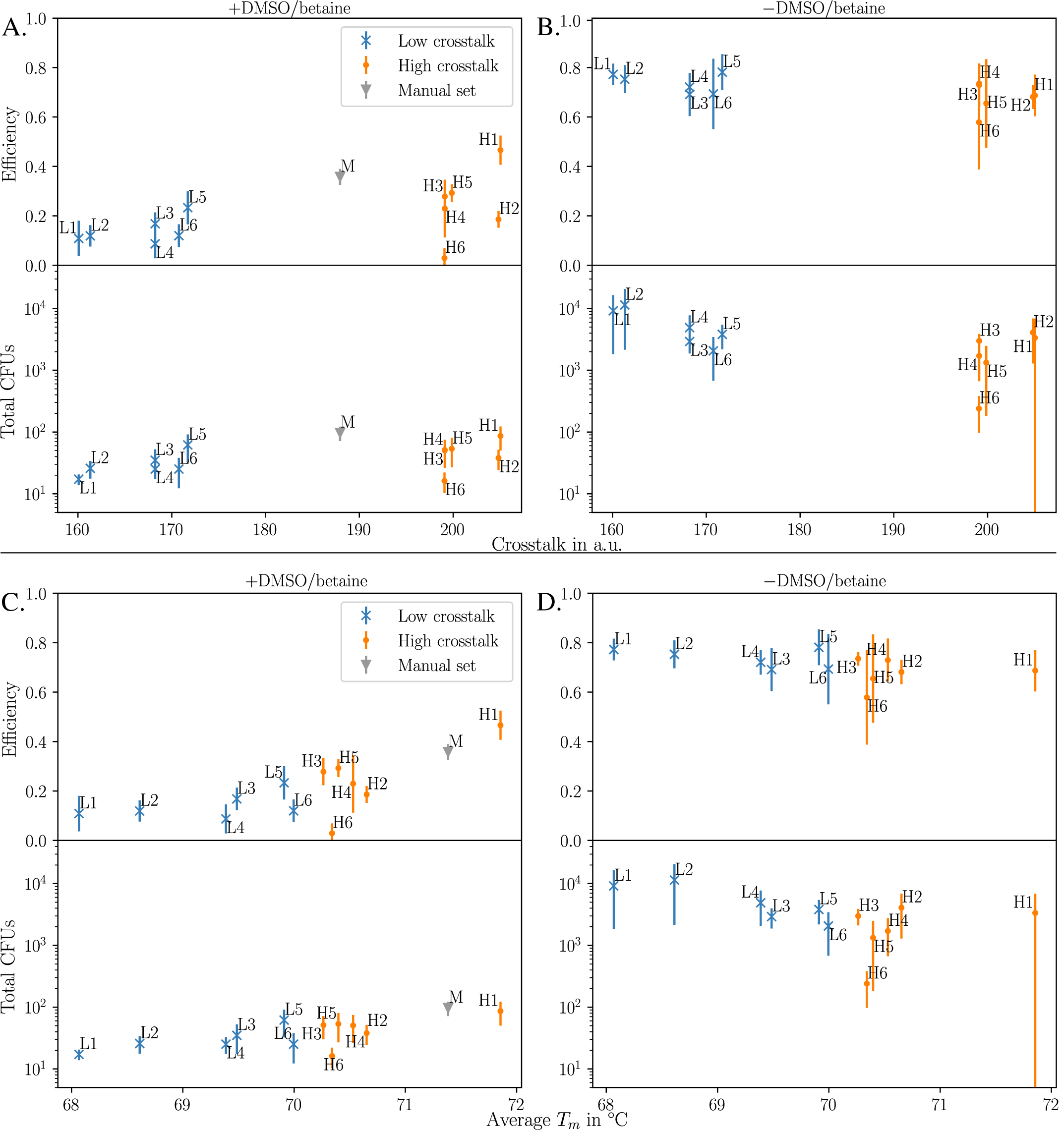
Graphical analysis of the bridging oligo sets (BO-sets) utilized for the assembly of a seven-part toy-plasmid, based on the results shown in Supplementary Figure 5. **A+B.** No crosstalk-dependent effects of the BOs were observed for the LCR with (A) and without (B) DMSO/betaine. Both clusters (L1-L6 and H1-H6) were distinguishable by the crosstalk but without affecting the LCR efficiency and total amount of colonies. Slightly higher efficiencies and more colonies were observed for the cluster of the sets H1-H6 when DMSO/betaine was used. **C+D.**: The average melting temperature of the BO-sets influenced the LCRs. Higher *T*_*m*_s resulted in higher efficiencies and more colonies when DMSO and betaine were used (C). Without DMSO and betaine, all BO-sets resulted in similar efficiencies suggesting no impact of crosstalk of BOs with the DNA parts in the LCR-assembly of the seven-part toy-plasmid. More colonies were observed for sets L1-L6. In contrast to LCRs with DMSO and betaine (C), the total amount of colonies was found to be increasing with decreasing melting temperatures. All *T*_*m*_s presented here were calculated by using the formula of SantaLucia (20) for the *T*_*m*_ calculation and salt correction. The manual set (“M”, used for LCRs with DMSO and betaine) was designed in `Geneious` with the same algorithms and a target temperature of 70 °C. The difference of ∼1.5 °C in comparison to the target *T*_*m*_ of 70 °C is due to the lack of an option to specify the DNA part concentration in the software. The raw data of the LCRs presented here are shown in Supplementary Figure 4. 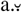.: arbitrary unit, BO: bridging oligo, CFU: colony forming unit, DMSO: dimethyl sulfoxide, M: manually designed BO-set, *T*_*m*_: melting temperature of a BO-half.

**Supplementary Figure 7.**
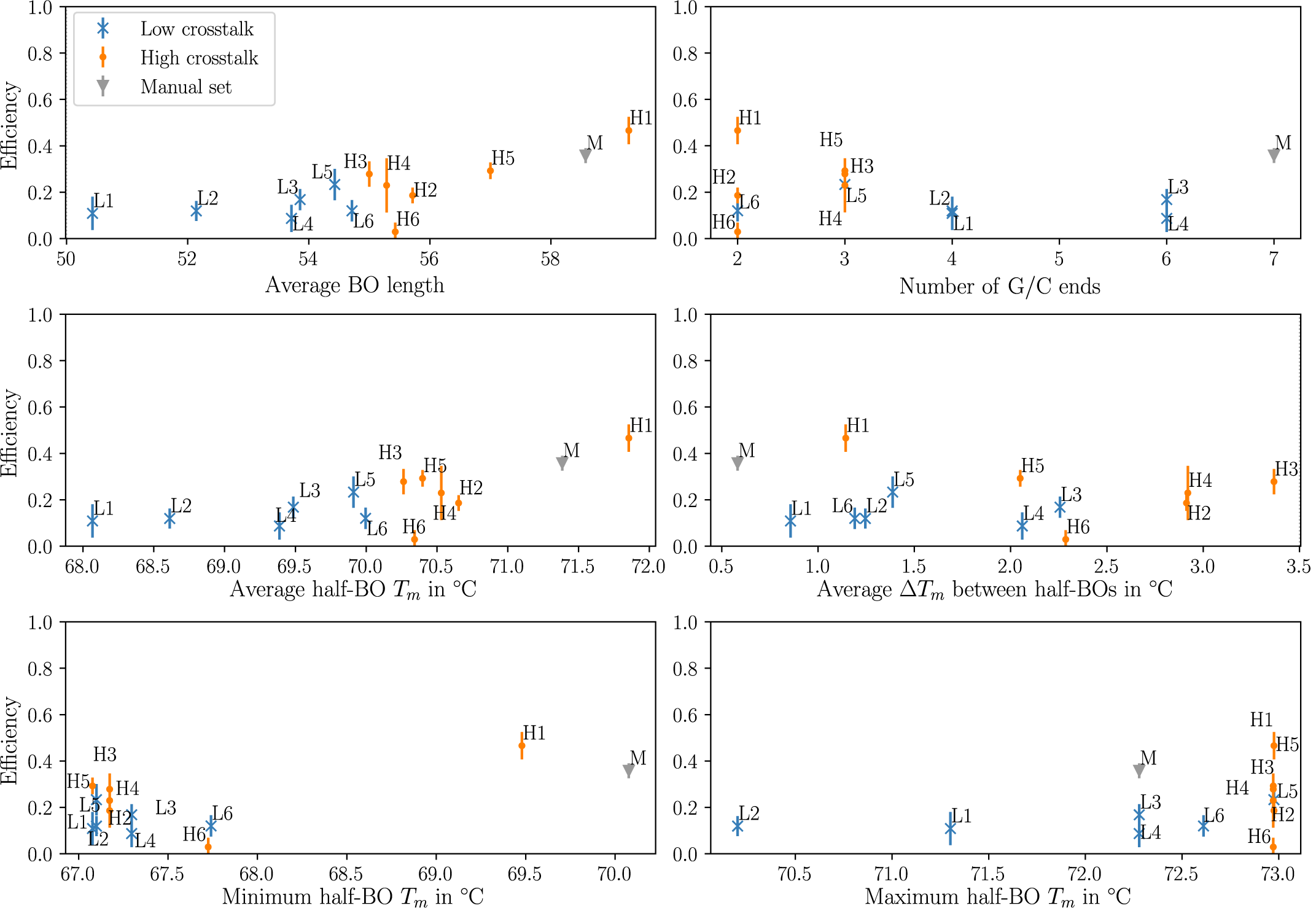
Graphical analysis of LCRs <u>with</u> 8 %v*/*v DMSO and 0.45 M betaine of the seven-part toy-plasmid and the influence of various predictors on the efficiencies by utilizing different bridging oligo sets. All LCRs were performed as quintuplets. The standard deviation for each LCR is indicated by error bars. All BO sequences are shown in Supplementary Table 3. BO: bridging oligo, DMSO: dimethyl sulfoxide, G/C end: 3’-end of one BO ending with the nucleobase guanine or cytosine, *T*_*m*_: melting temperature.

**Supplementary Figure 8.**
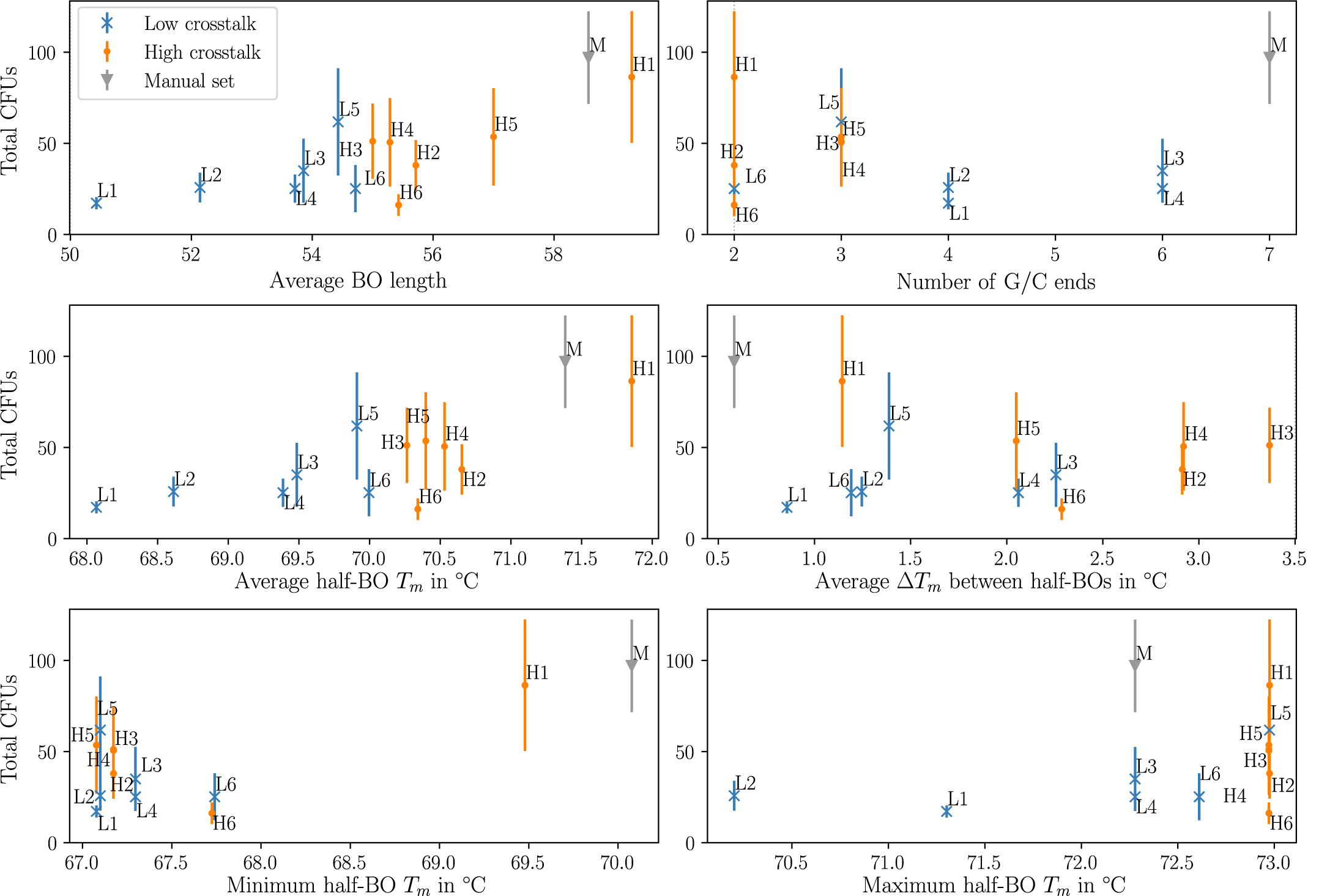
Graphical analysis of LCRs <u>with</u> 8 %v*/*v DMSO and 0.45 M betaine of the seven-part toy-plasmid and the influence of various predictors on the total amount of colonies by utilizing different bridging oligo sets. All LCRs were performed as quintuplets. The standard deviation for each LCR is indicated by error bars. All BO sequences are shown in Supplementary Table 3. BO: bridging oligo, CFU: colony forming unit, DMSO: dimethyl sulfoxide, G/C end: 3’-end of one BO ending with the nucleobase guanine or cytosine, *T*_*m*_: melting temperature.

**Supplementary Figure 9.**
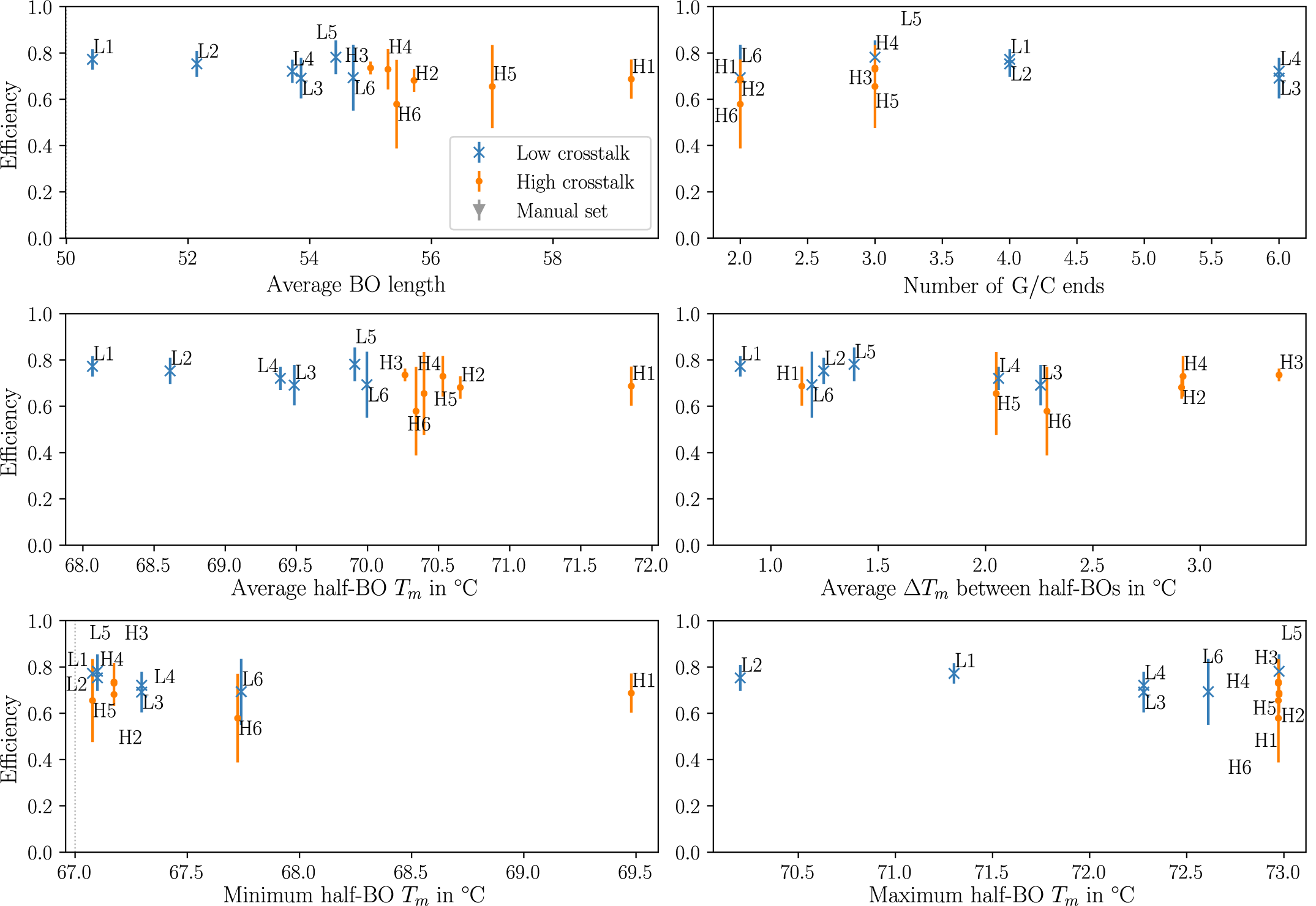
Graphical analysis of LCRs <u>without</u> 8 %v*/*v DMSO and 0.45 M betaine of the seven-part toy-plasmid and the influence of various predictors on the efficiencies by utilizing different bridging oligo sets. All LCRs were performed as quintuplets. The standard deviation for each LCR is indicated by error bars. All BO sequences are shown in Supplementary Table 3. BO: bridging oligo, DMSO: dimethyl sulfoxide, G/C end: 3’-end of one BO ending with the nucleobase guanine or cytosine, *T*_*m*_: melting temperature.

**Supplementary Figure 10.**
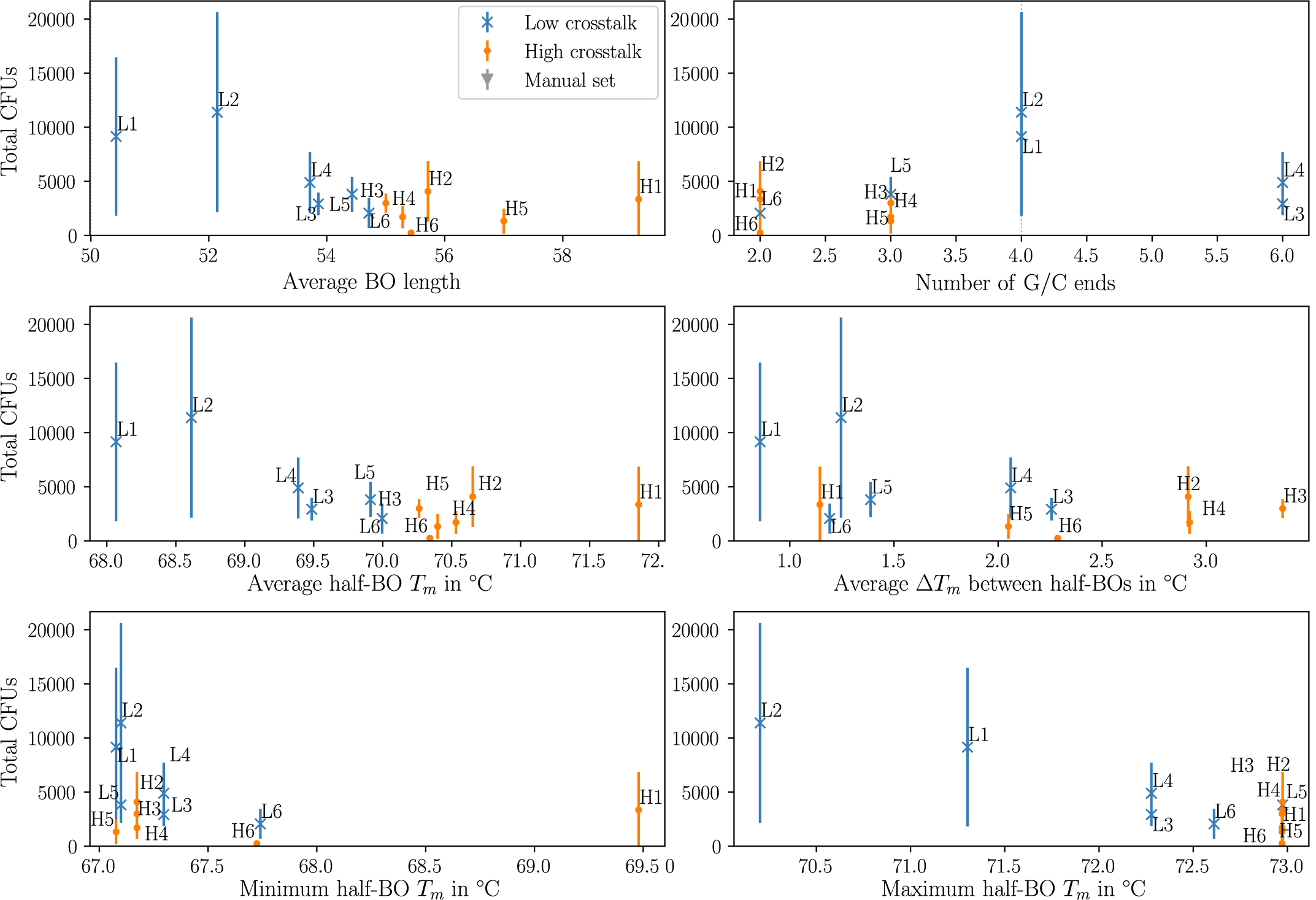
Graphical analysis of LCRs <u>without</u> 8 %v*/*v DMSO and 0.45 M betaine of the seven-part toy-plasmid and the influence of various predictors on the total amount of colonies by utilizing different bridging oligo sets. All LCRs were performed as quintuplets. The standard deviation for each LCR is indicated by error bars. All BO sequences are shown in Supplementary Table 3. BO: bridging oligo, CFU: colony forming unit, DMSO: dimethyl sulfoxide, G/C end: 3’-end of one BO ending with the nucleobase guanine or cytosine, *T*_*m*_: melting temperature.

**Supplementary Figure 11.**
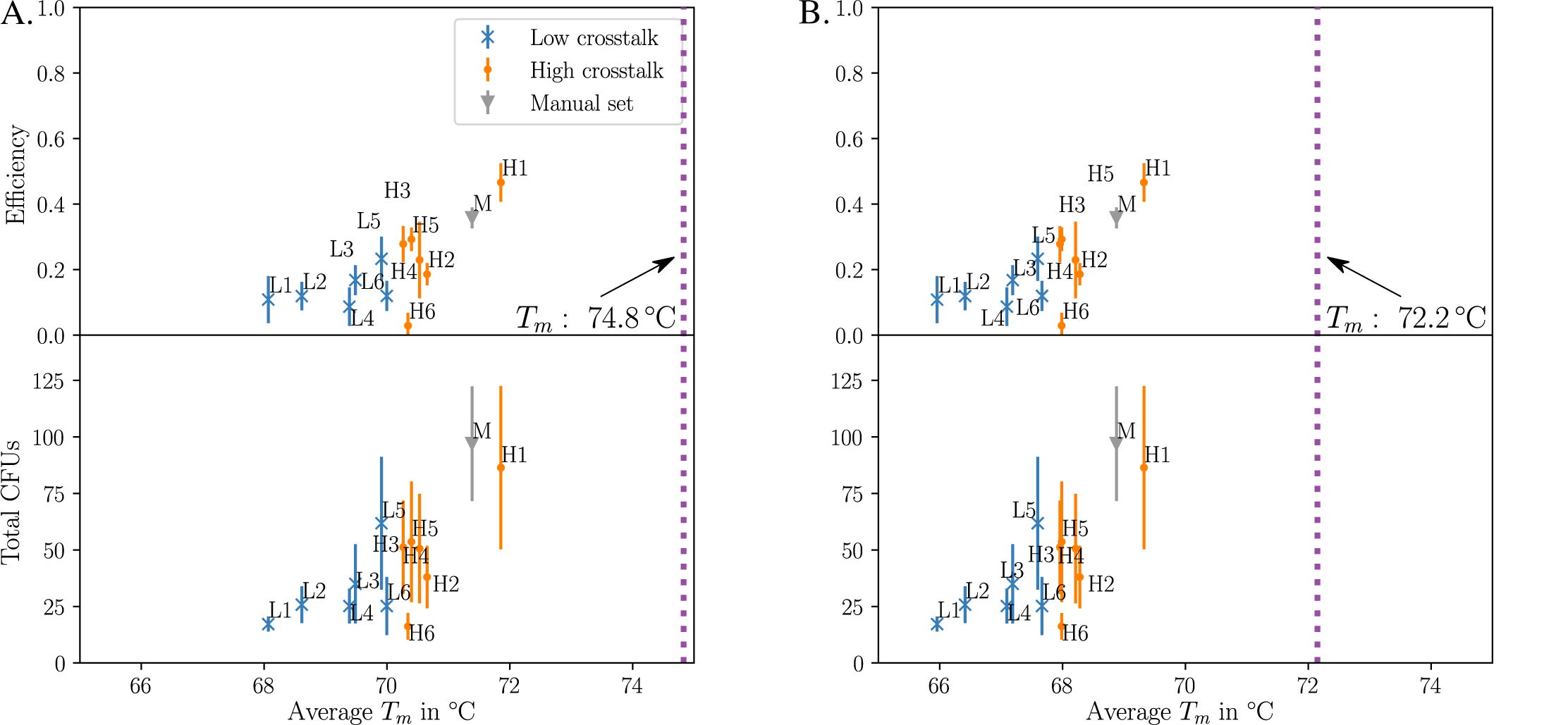
Effect of salt corrections on melting temperature. The results for the LCRs with 8 %v*/*v DMSO and 0.45 M are presented here. The thermodynamic parameters used were SantaLucia (20). The standard deviation for each LCR is indicated by error bars. **A**. SantaLucia (20) salt correction. **B**. Owczarzy (35) salt correction. The optimized melting temperature indicated by the dotted line is obtained from recalculation of the optimized BO-set of the de Kok et al. (8) study. BO: bridging oligo, CFU: colony forming unit, DMSO: dimethyl sulfoxide, *T*_*m*_: melting temperature of one BO-half.

**Supplementary Figure 12.**
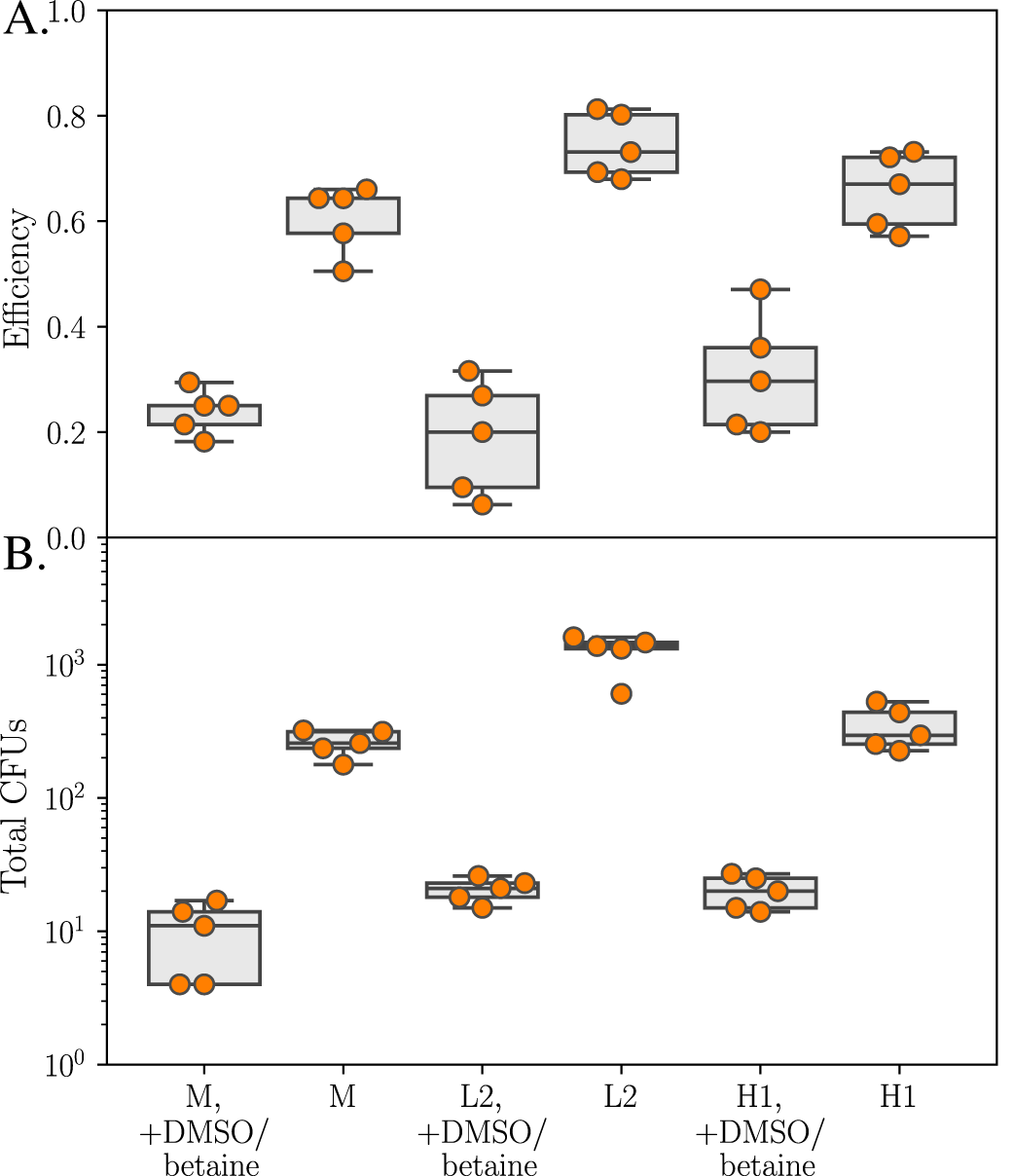
Influence of DMSO and betaine in the LCR. All LCRs were performed as quintuplets using the same batches of DNA parts, electrocompetent cells and a master-mix (excluding the BO-sets and the ligase). **A.** For all three BO-sets (H1, L2 and the manually set) the DMSO/betaine-free LCRs were more efficient. As observed in Supplementary Figure 5, the efficiency of the set “H1” was higher than the manual set “M” and set “L2” in the baseline LCR (with DMSO and betaine). **B.** The omission of DMSO/betaine highly increased the total amount of colonies. This is consistent with the results shown in Supplementary Figures 5B and 6H for DMSO/betaine-free conditions: a lower *T*_*m*_ is beneficial and resulted in more colonies. All BO sequences are shown in Supplementary Table 3.

**Supplementary Figure 13.**
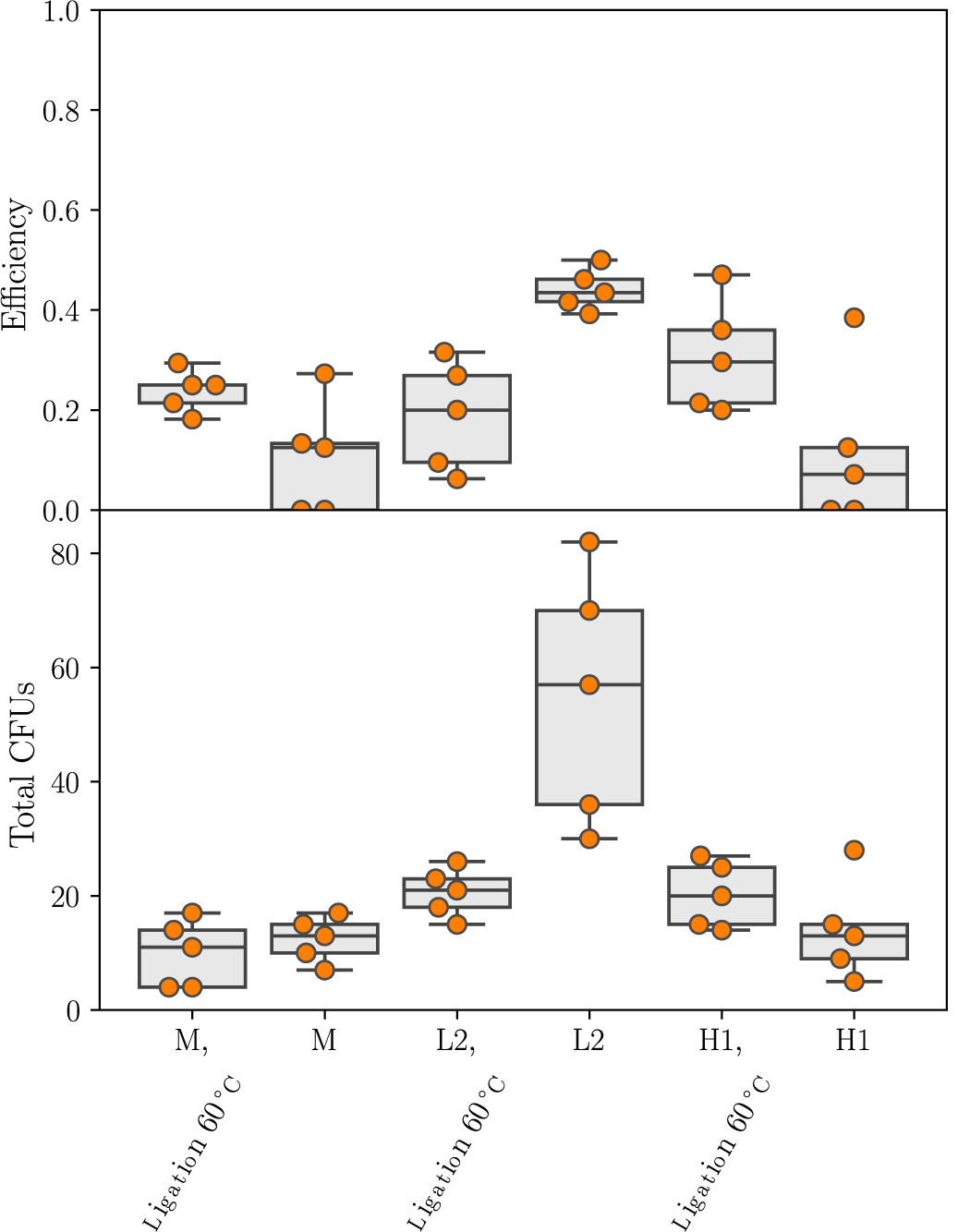
Impact of the ligation temperature on LCR performance with DMSO and betaine. All LCRs were performed as quintuplets. A lower ligation temperature is beneficial for the efficiencies and total amount of colonies for the LCR using BO-set L2. This set had a lower *T*_*m*_ of 68.6 °C compared to the manual set with a *T*_*m*_ of 71.4 °C and the set H1 with a *T*_*m*_ of 71.9 °C. Due to the lowered ligation temperature, more BOs of set L2 remained attached to the DNA parts to guide the ligase. All BO sequences are shown in Supplementary Table 3.

**Supplementary Figure 14.**
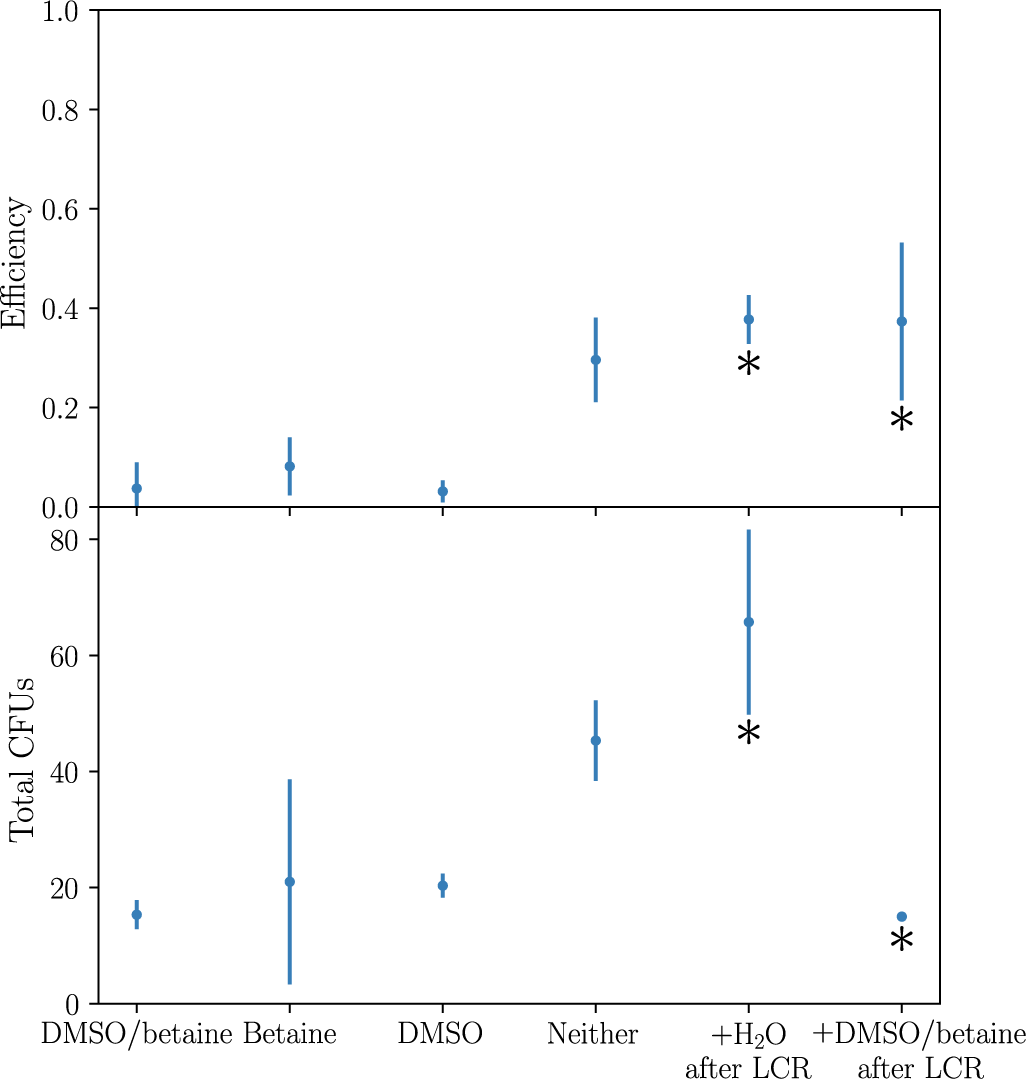
DMSO and betaine negatively affect the LCR of the seven-part plasmid. The LCRs to investigate the impact of DMSO and/or betaine were performed as triplicates. To investigate the direct influence of DMSO and betaine on the electroporation process, the LCRs were performed as quadruplets (indicated with a *). The standard deviation for each LCR is indicated by error bars and the manual set “M” was used. The combination of DMSO and betaine negatively affects the efficiency and the total amount of colonies (comparing the results for DMSO/betaine with none of both). Further investigation revealed a direct negative impact of DMSO/betaine in the electroporation. An LCR was performed without DMSO/betaine. Both detergents were added before the electroporation to simulate the transformation conditions (8 %v*/*v DMSO and 0.45 M betaine, mix of 3 µL LCR and 30 µL competent cells). As a control *dd*H_2_O was added. All BO sequences are shown in Supplementary Table 3. BO: bridging oligo, CFU: colony forming unit, DMSO: dimethyl sulfoxide.

**Supplementary Figure 15.**
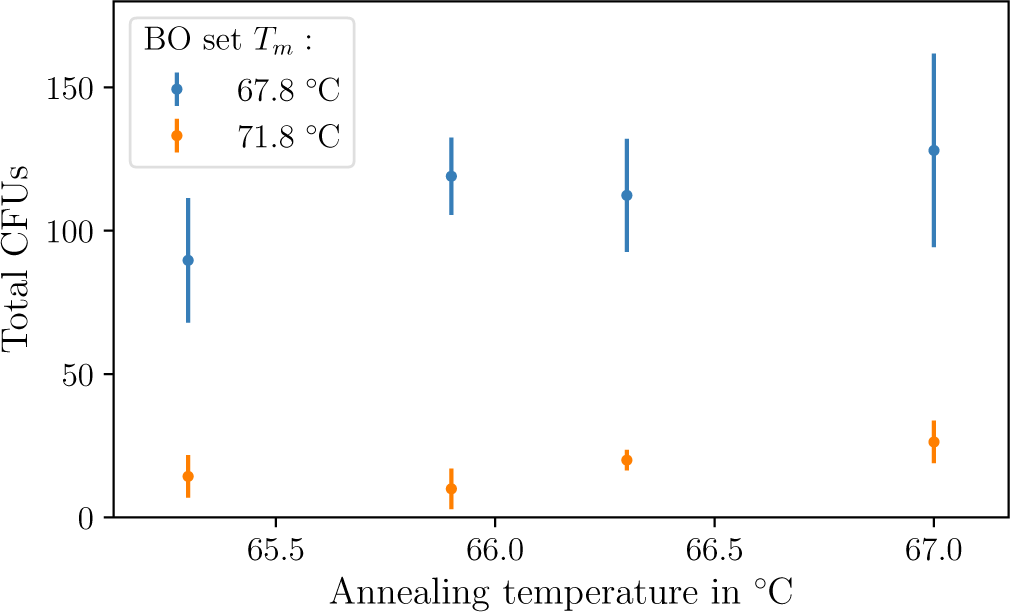
Total amount of colonies for the bridging oligo sets “67.8 °C” and “71.8 °C” in the annealing temperature range of 65-67 °C (larger temperature range shown in Figure 3A+B). All LCRs were performed as triplicates. The standard deviation for each LCR is indicated by error bars. All BO sequences are shown in Supplementary Table 4. BO: bridging oligo, CFU: colony forming unit, DMSO: dimethyl sulfoxide, *T*_*m*_: melting temperature of one BO-half.

**Supplementary Figure 16.**
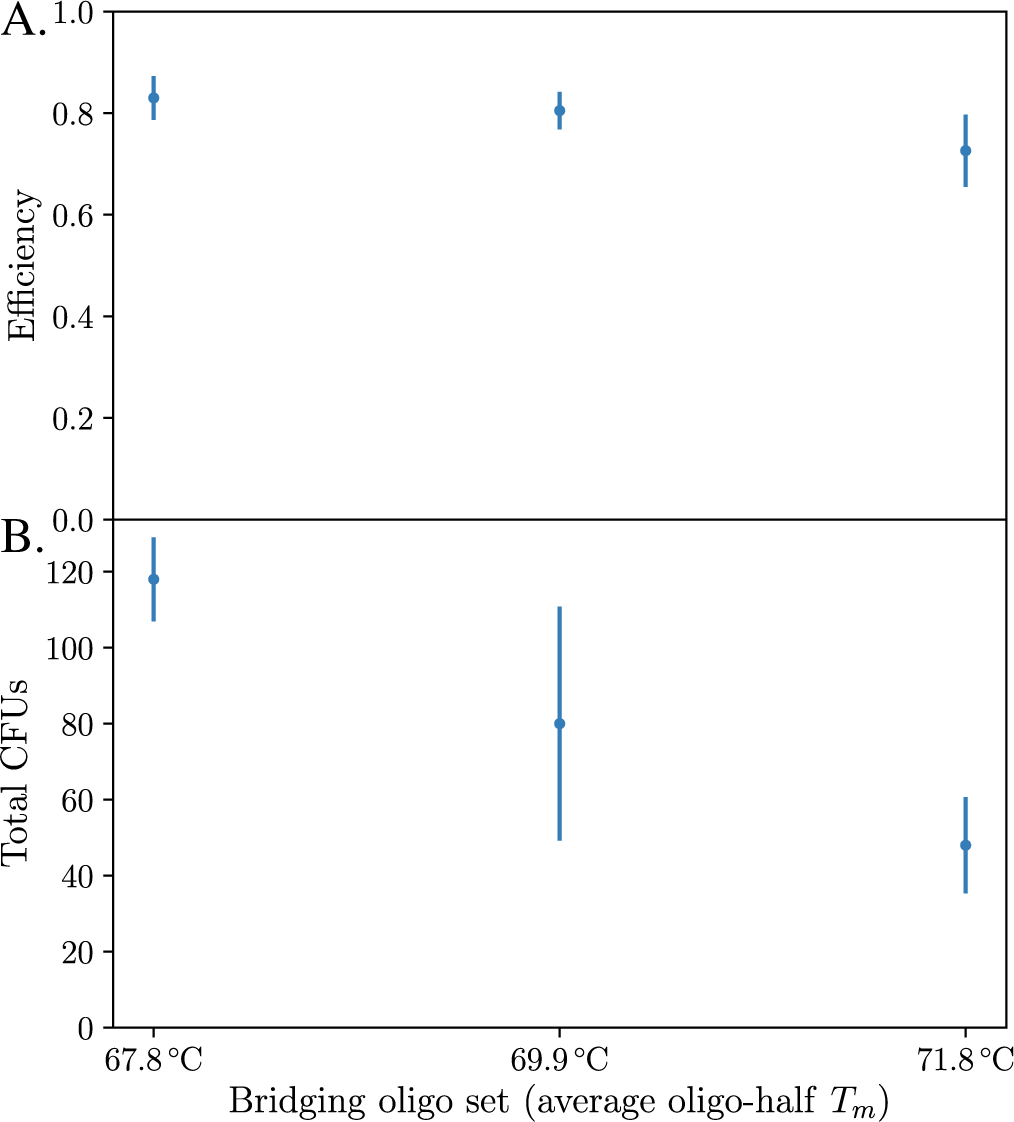
LCR without DMSO and betaine at the annealing temperature of 67.9 °C using three bridging oligo sets (*T*_*m*_ = 67.8, 69.9 and 71.8 °C). All LCRs were performed as triplicates. The standard deviation for each LCR is indicated by error bars. A larger range of the annealing temperature is shown in Figure 3A+B. **A.** The efficiencies of the LCRs were similar. **B.** The total amount of colonies increased with lower melting temperatures of the BOs. All BO sequences are shown in Supplementary Table 4. BO: bridging oligo, CFU: colony forming unit, DMSO: dimethyl sulfoxide, *T*_*m*_: melting temperature of one BO-half.

**Supplementary Figure 17.**
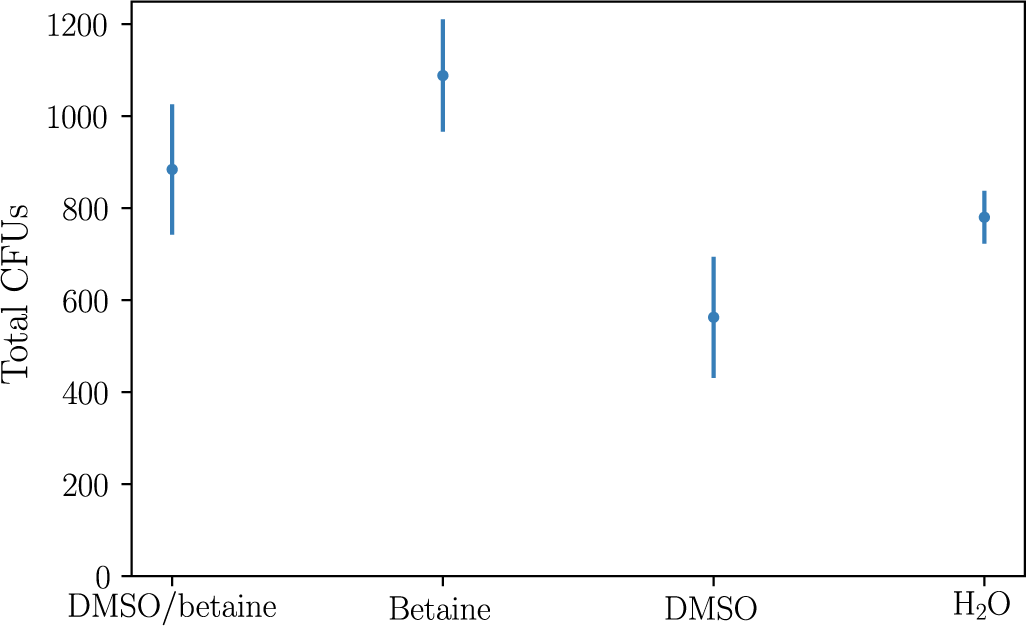
DMSO and betaine in chemical transformations. All transformations were performed as triplicates. The standard deviation for each LCR is indicated by error bars. The plasmid pUC19 was mixed with DMSO and/or betaine or *aq. dest.* followed by chemical transformation of *E. coli*. DMSO had a negative impact whereas betaine had a positive impact on the number of colonies. Adding both yielded similar results as adding neither. CFU: colony forming unit, DMSO: dimethyl sulfoxide.

### Sequences of the toy-model plasmid and the utilized amplification primers and bridging oligos

**Supplementary Table 1:**
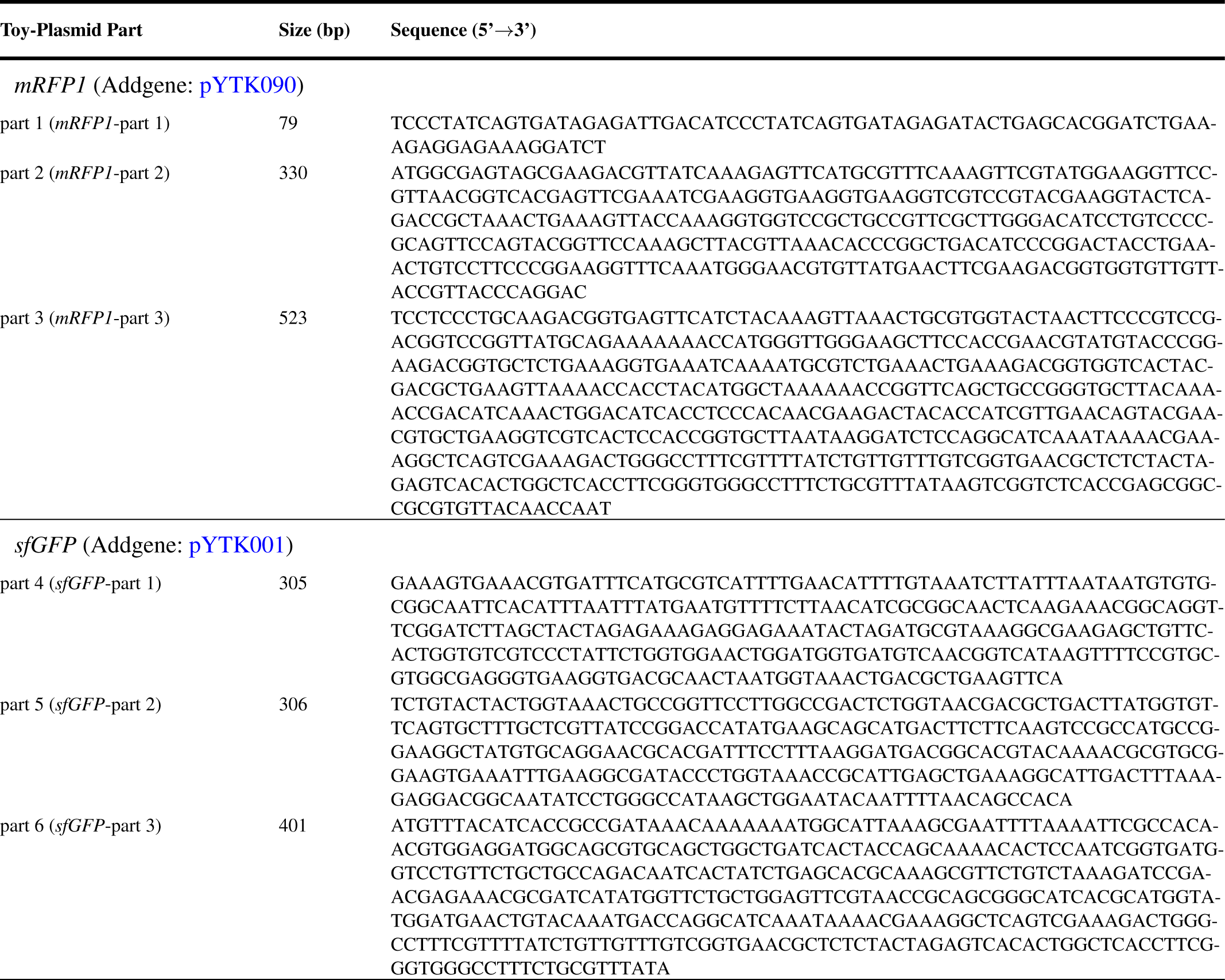
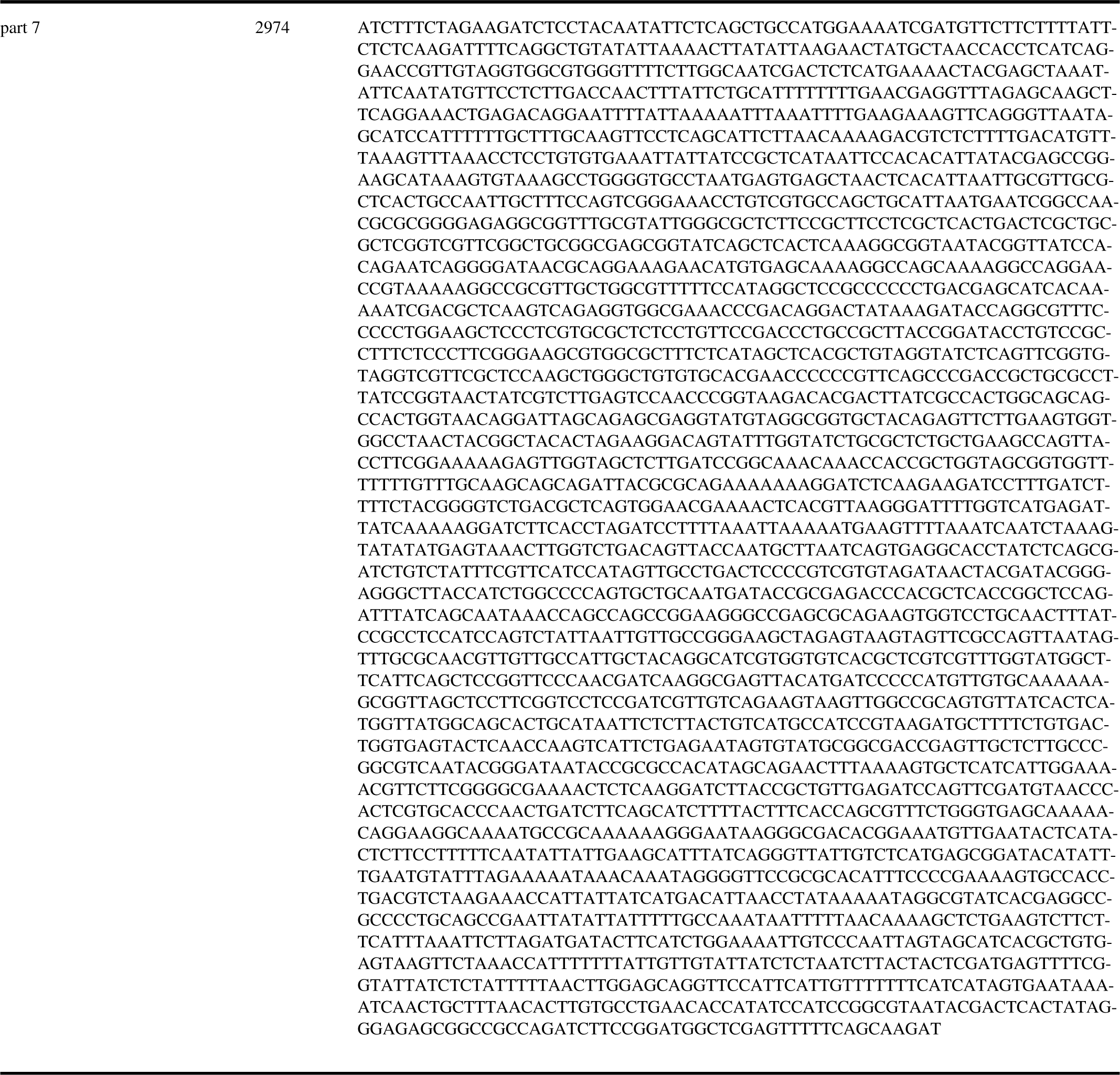
Sequences of the seven-part split of the toy-model (Figure 1B). For the three-part split parts 1-3, 4-6 and 7 were used. The bridging oligos were designed by concatenating subsequences of these parts. It was not necessary to use the reverse complement for the design because the experiments contained both strands of these parts. A GenBank-file of the toy-model plasmid is available in the digital supplement and at www.kabisch-lab.de/data/SynBioSuppBOfasta.zip. *mRFP1*: monomeric red fluorescent protein 1, *sfGFP*: superfolder green fluorescent protein.

**Supplementary Table 2:**
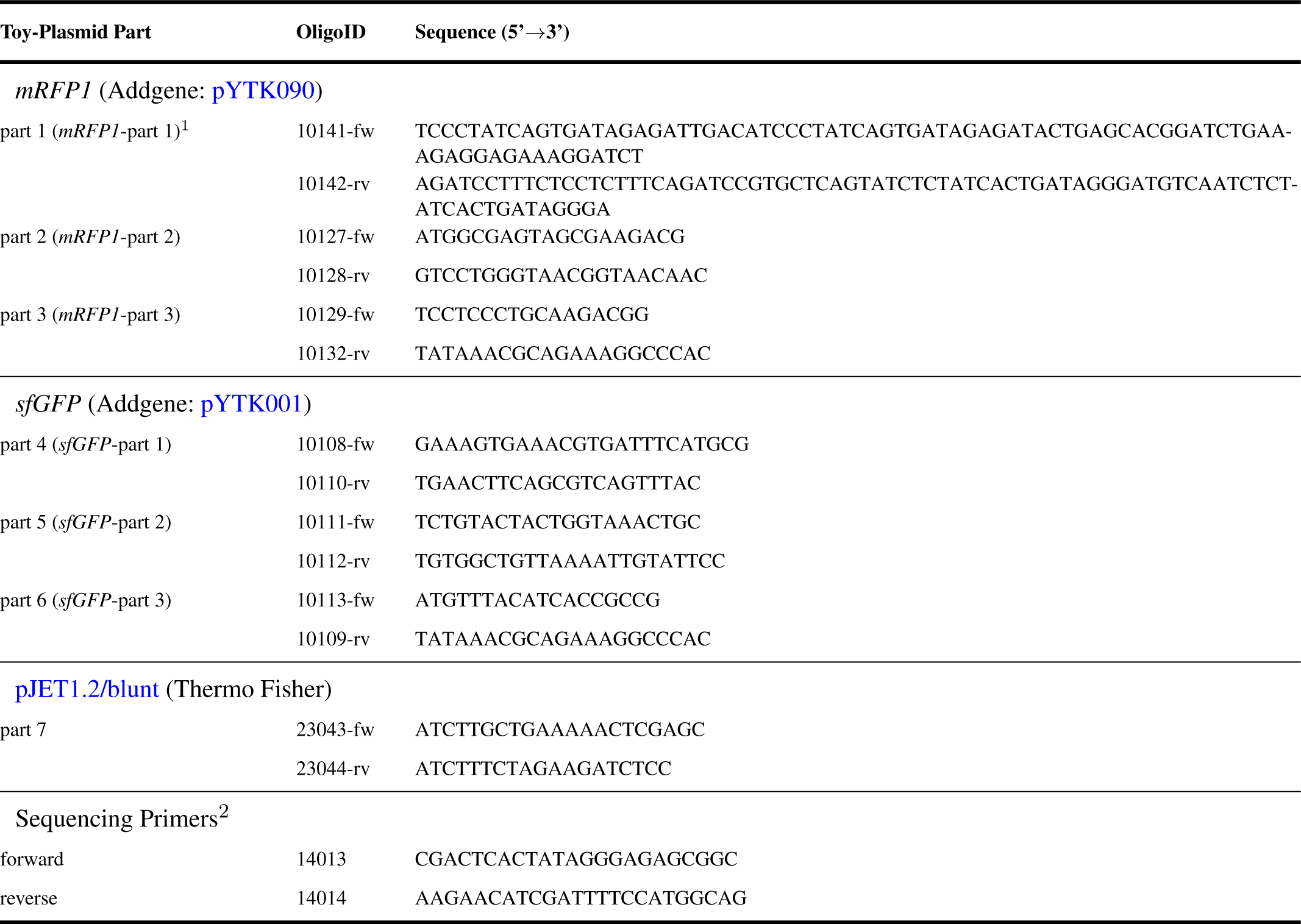
Oligonucleotides for the amplification of all toy-model plasmid parts (shown in Figure 1B). ^1^ The complete part was ordered as two oligonucleotides, which were phosphorylated and annealed to double stranded DNA before the LCR (described in the methods). ^2^ For sequencing from the vector into the inserts. fw: forward direction, *mRFP1*: monomeric red fluorescent protein 1, rv: reverse direction, *sfGFP*: superfolder green fluorescent protein.

**Supplementary Table 3:**
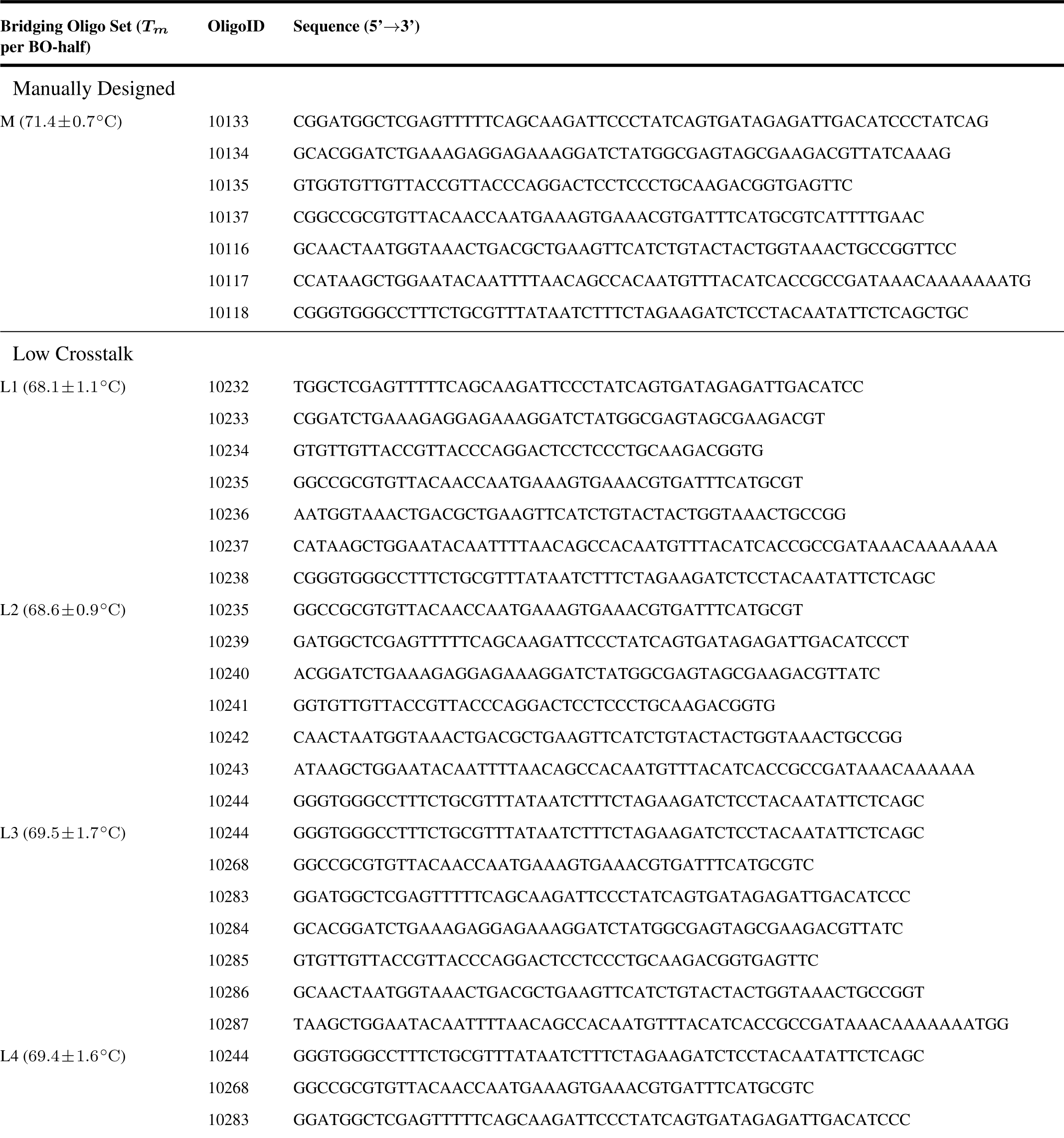
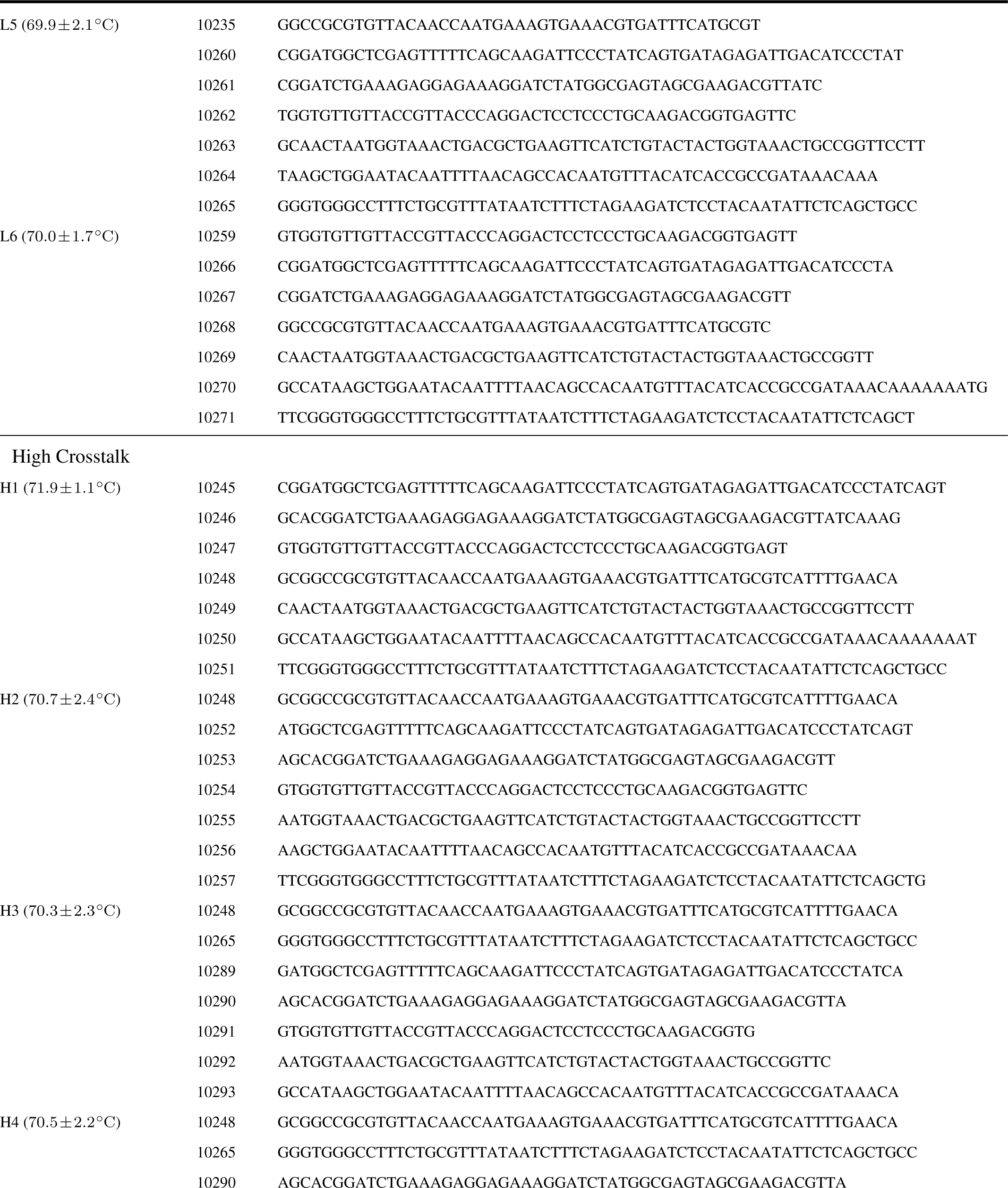
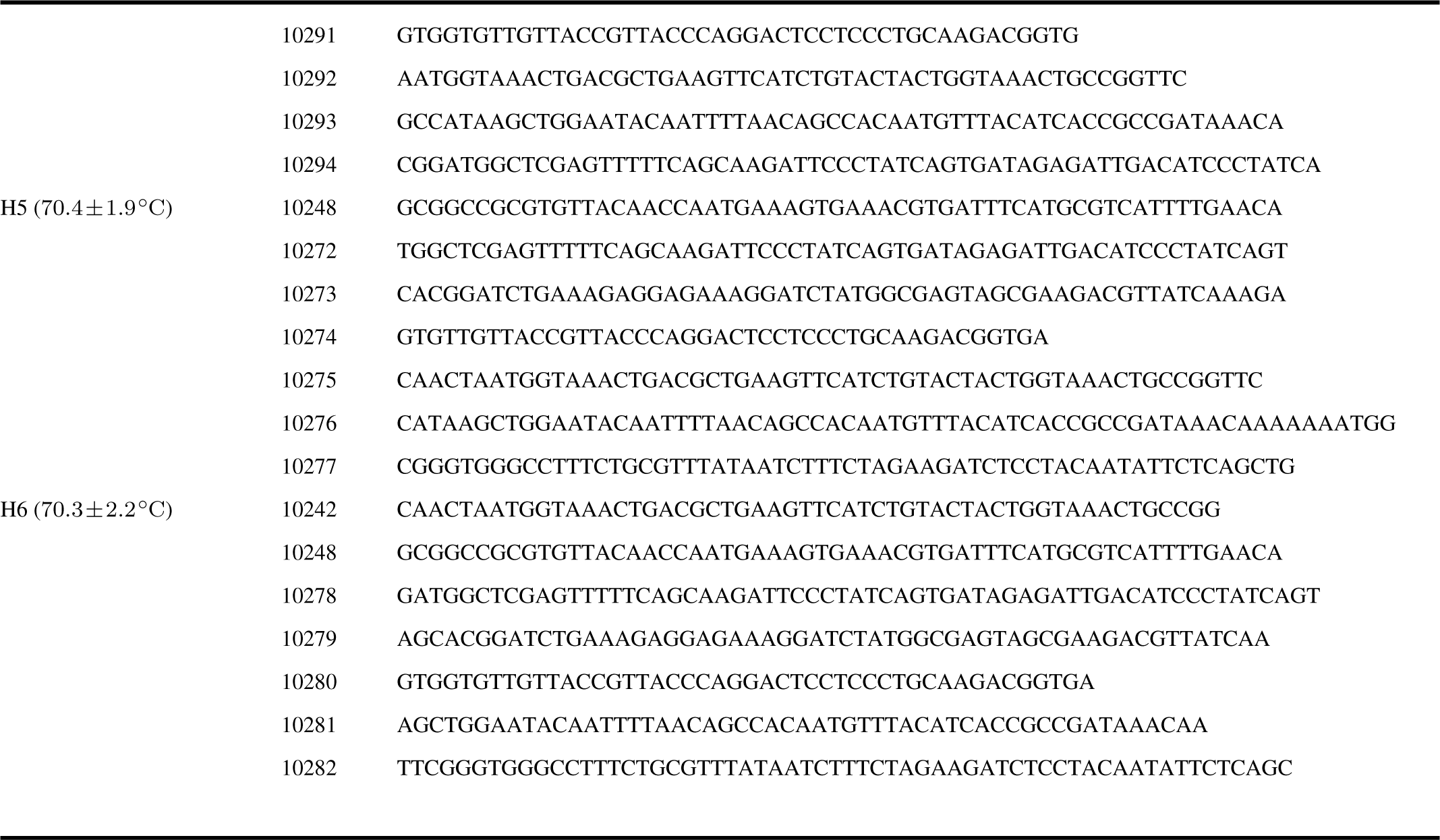
Oligonucleotides for crosstalk LCRs (results shown in Supplementary Figures 5 and 6). All melting temperatures (*Tm*s) presented here are calculated for each BO-half using the formula of SantaLucia (20) for the *Tm*-calculation and the salt correction. *Tm*: melting temperature of a BO-half.

**Supplementary Table 4:**
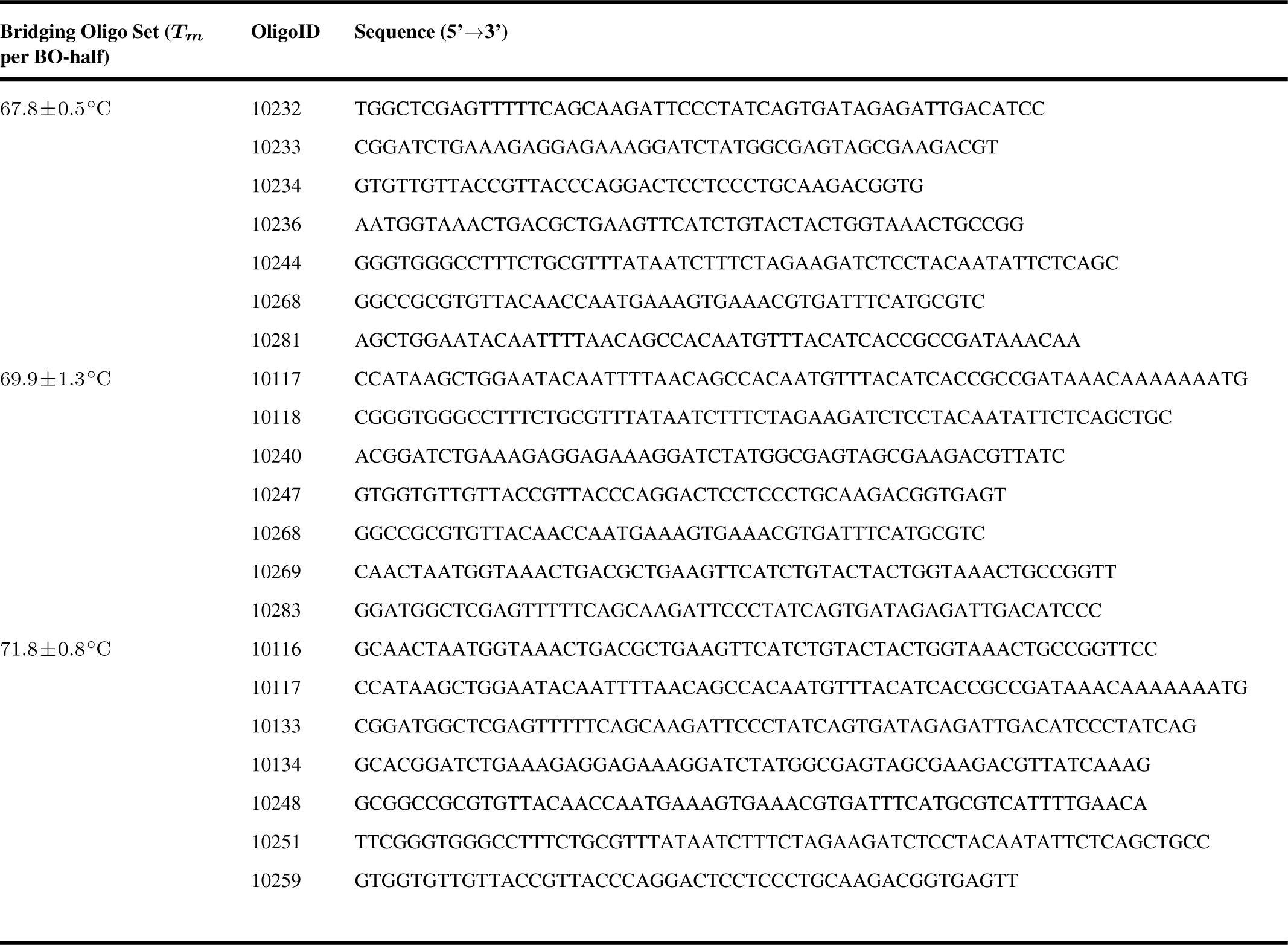
Bridging oligo sets for the gradient-LCR (composed of bridging oligos for crosstalk experiments; Supplementary Table 3). The results of the gradient-LCRs are shown in Figure 3A+B. All melting temperatures (*Tm*s) presented here are calculated for each BO-half using the formula of SantaLucia (20) for the *Tm*-calculation and the salt correction. *Tm*: melting temperature of a BO-half.

**Supplementary Table 5:**
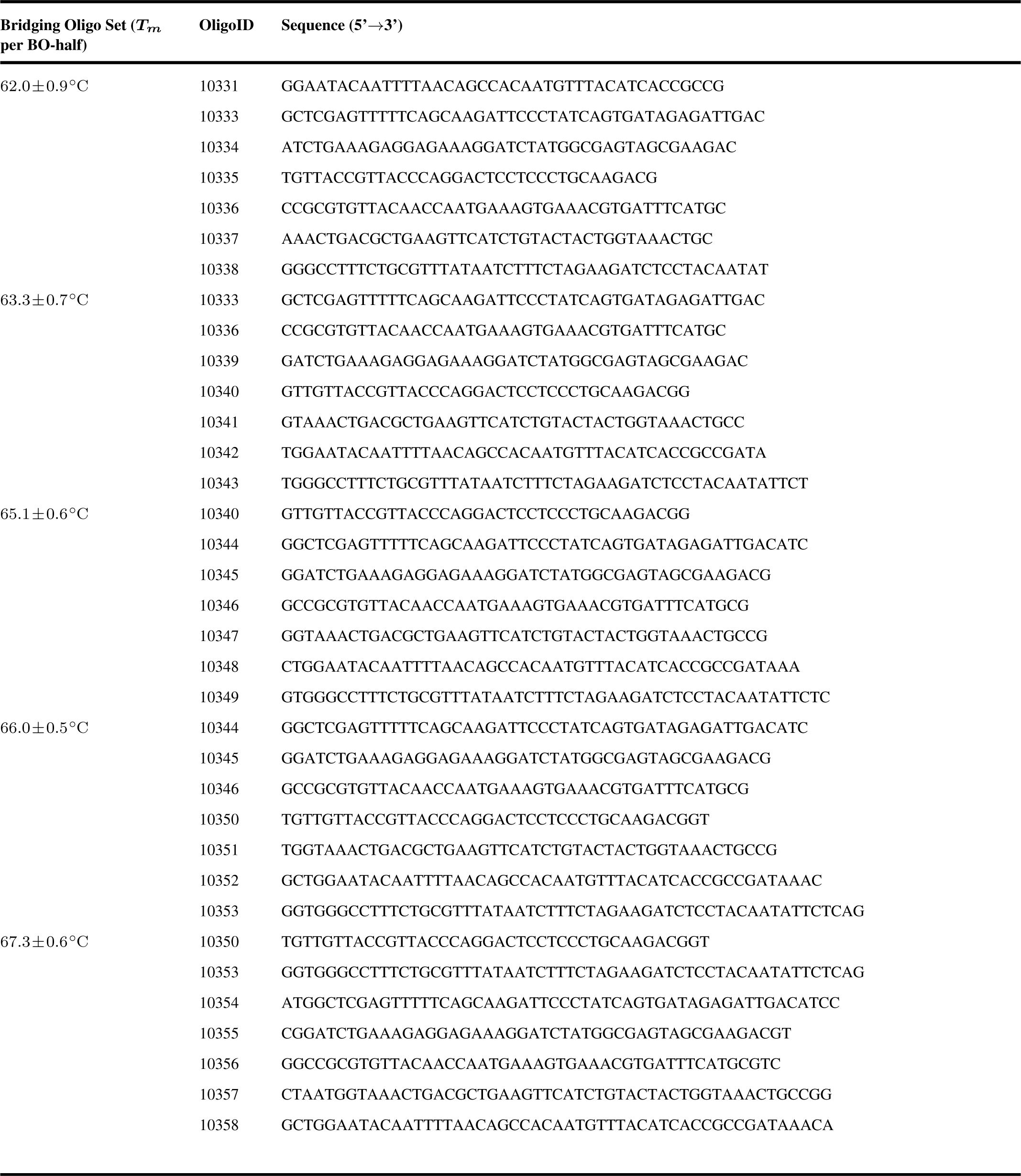
Bridging oligo sets with temperatures in the range of 62.0 °C to 67 °C. The results of the LCRs are shown in Figure 3C+D. All melting temperatures (*Tm*s) presented here are calculated for each BO-half using the formula of SantaLucia (20) for the *Tm*-calculation and the salt correction. *Tm*: melting temperature of a BO-half.

### Validation experiments

**Supplementary Figure 18.**
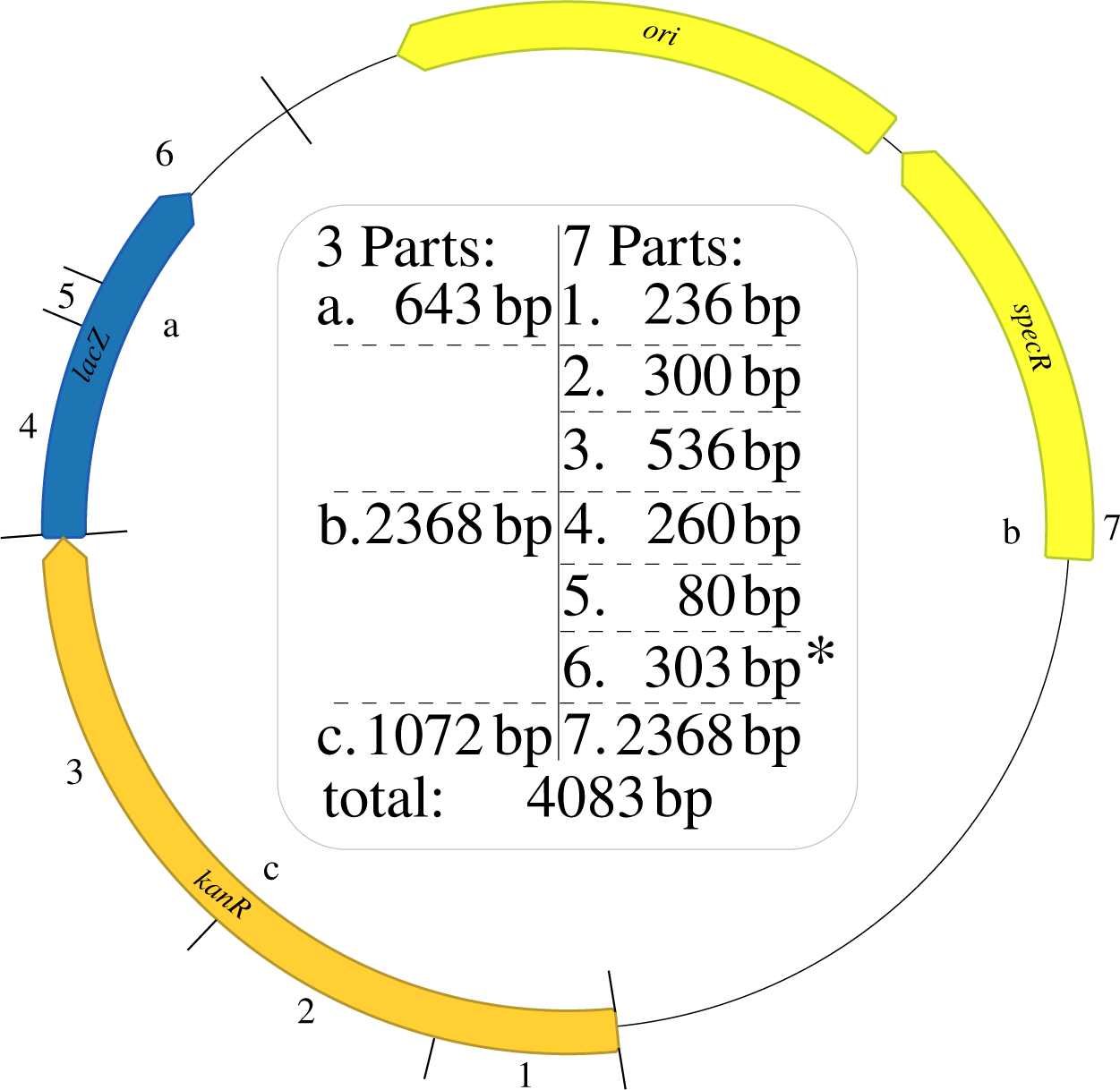
Plasmid 1 (“VP1”) for the validation experiments of the improved LCR conditions (Figure 4, Table 1). The plasmid was split in three and seven parts. Only if all parts were assembled the cells were able to replicate, were resistant to spectinomycin and turned blue by using the blue-white screening. Part 7 was PCR-amplified from the already existing plasmid build for scar-free deletions in *Bacillus subtilis* (41). The *lacZ* gene was inserted to enable a blue-white screening and was derived from the TALEN Kit 2.0 from Addgene (42). In comparison to the original flanking regions of the *lacZ* some mutations were present due to subcloning. *: Part 6 was amplified from the existing validation plasmid 2 (“VP2”) shown in Supplementary Figure 19. The *kanR* gene was derived from (43). Before PCR-amplification, all templates were digested by restriction enzymes. Afterwards, all PCR-products were additionally digested by a DpnI-digestion. *lacZ*: *β*-galactosidase, *ori*: origin of replication, *specR*: spectinomycin resistance gene.

**Supplementary Figure 19.**
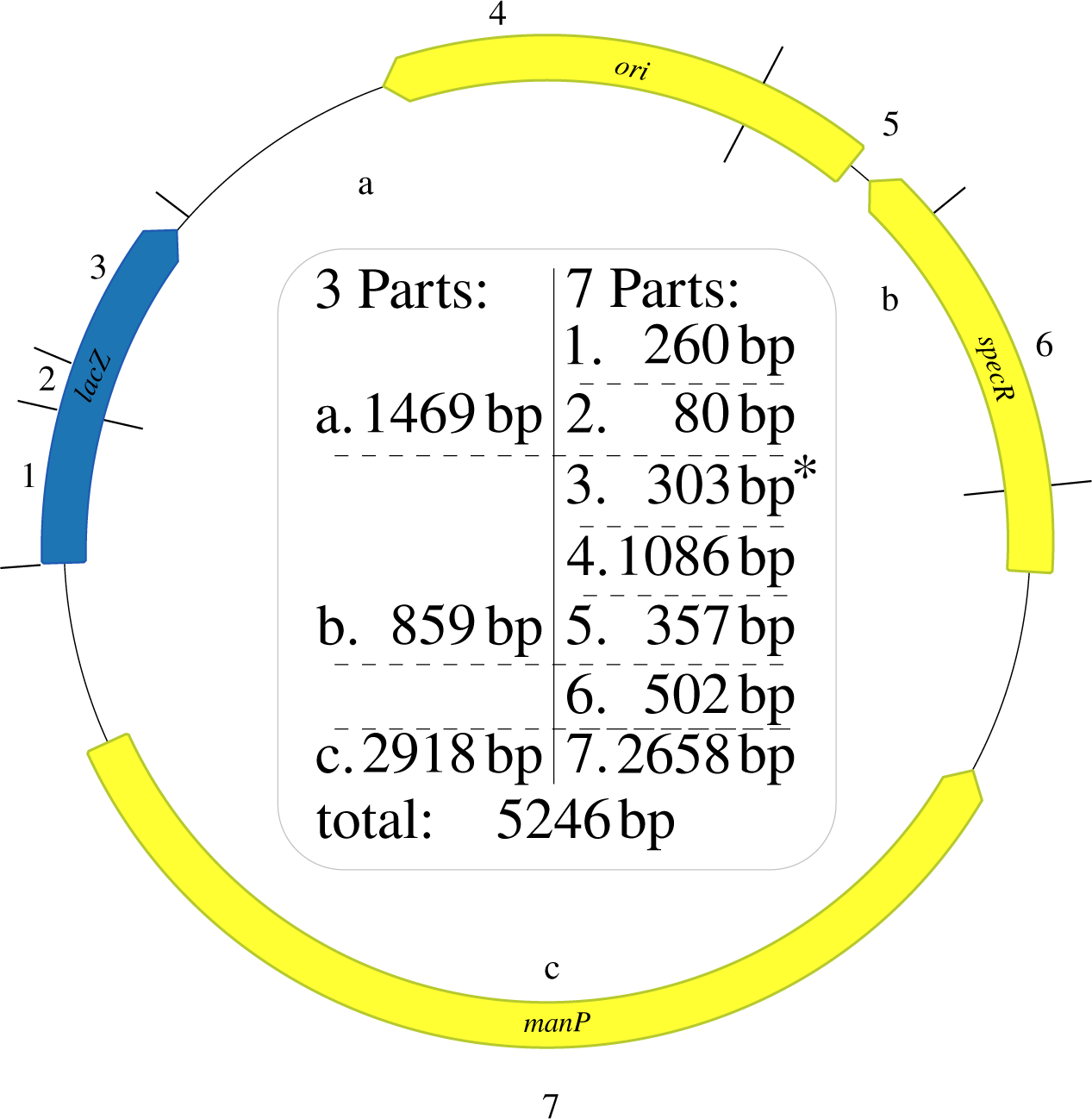
Plasmid 2 (“VP2”) for the validation experiments of the improved LCR conditions (Figure 4, Table 1). The plasmid was split in three and seven parts. Only if all parts were assembled the cells were able to replicate, were resistant to spectinomycin and kanamycin and turned blue by using blue-white screening. Parts 1-3 were derived by PCR-amplification of a plasmid from Wenzel et al. (41) and. Part 3 (*) begins in the *lacZ* of the plasmid derived from TALEN Kit 2.0 from Addgene (42) and ends in the plasmid derived from Wenzel et al. (41). Parts 4-7 were PCR-amplified from the already existing plasmid build for scar-free deletions in *Bacillus subtilis* (41). Before PCR-amplification, all templates were digested by restriction enzymes. Afterwards, all PCR-products were digested by DpnI. *lacZ*: *β*-galactosidase, *manP*: D-mannose permease (for *Bacillus* spp.), *ori*: origin of replication), *specR*: spectinomycin resistance gene.

**Supplementary Figure 20.**
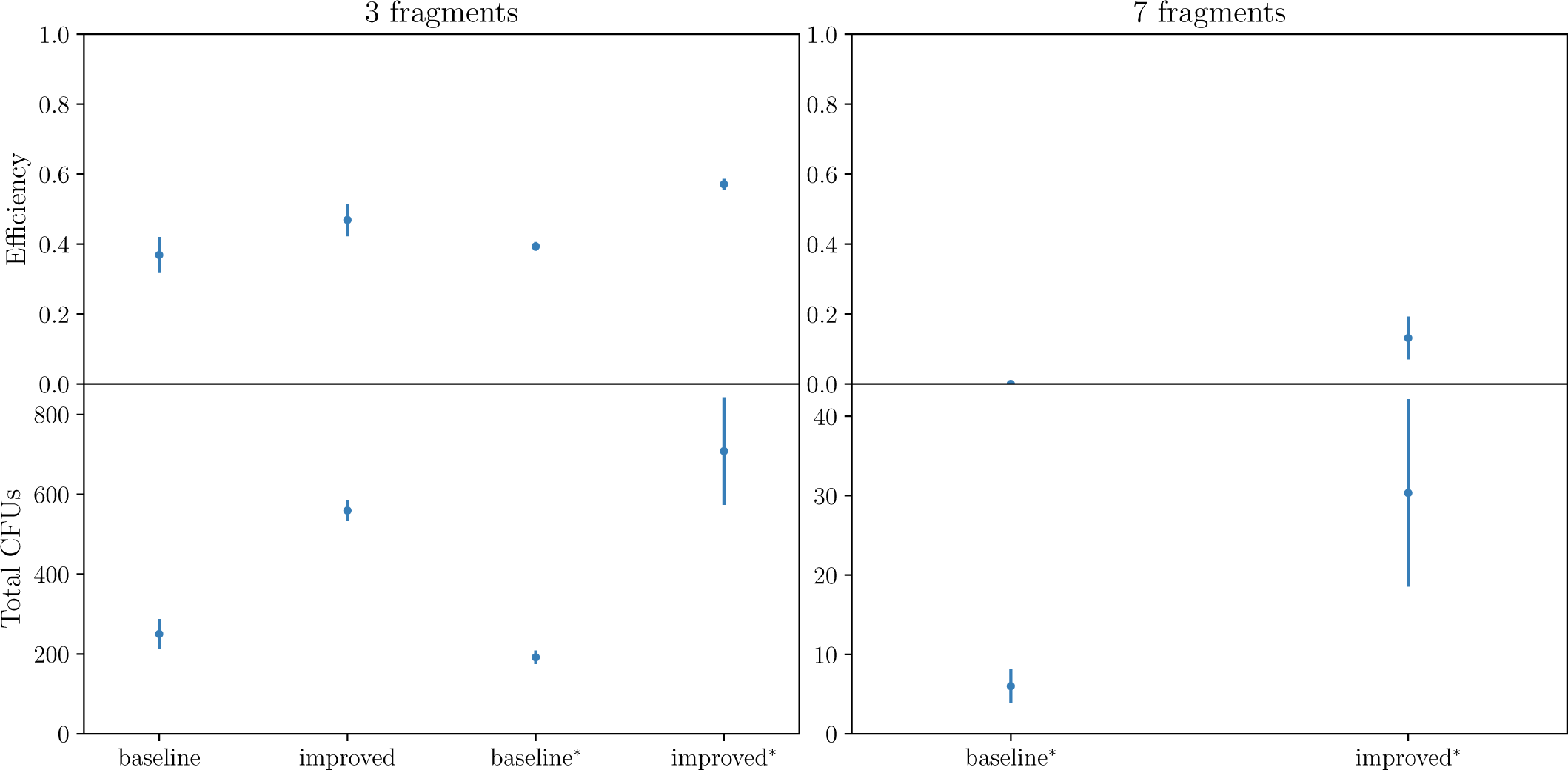
Results of the baseline and improved LCR-protocol for the toy-model plasmid (“TP”). For the three-part split, the same LCRs were transformed in chemical and in ^*\**^ electrocompetent cells to prove that the results are independent of the transformation method. For the seven-part LCR only electroporation resulted in CFUs. For all LCRs, 3 µL were transformed in 30 µL cells. All LCRs were performed as triplicates. BO: bridging oligo, CFU: colony forming unit, DMSO: dimethyl sulfoxide, *T*_*m*_: melting temperature of one BO-half.

**Supplementary Figure 21.**
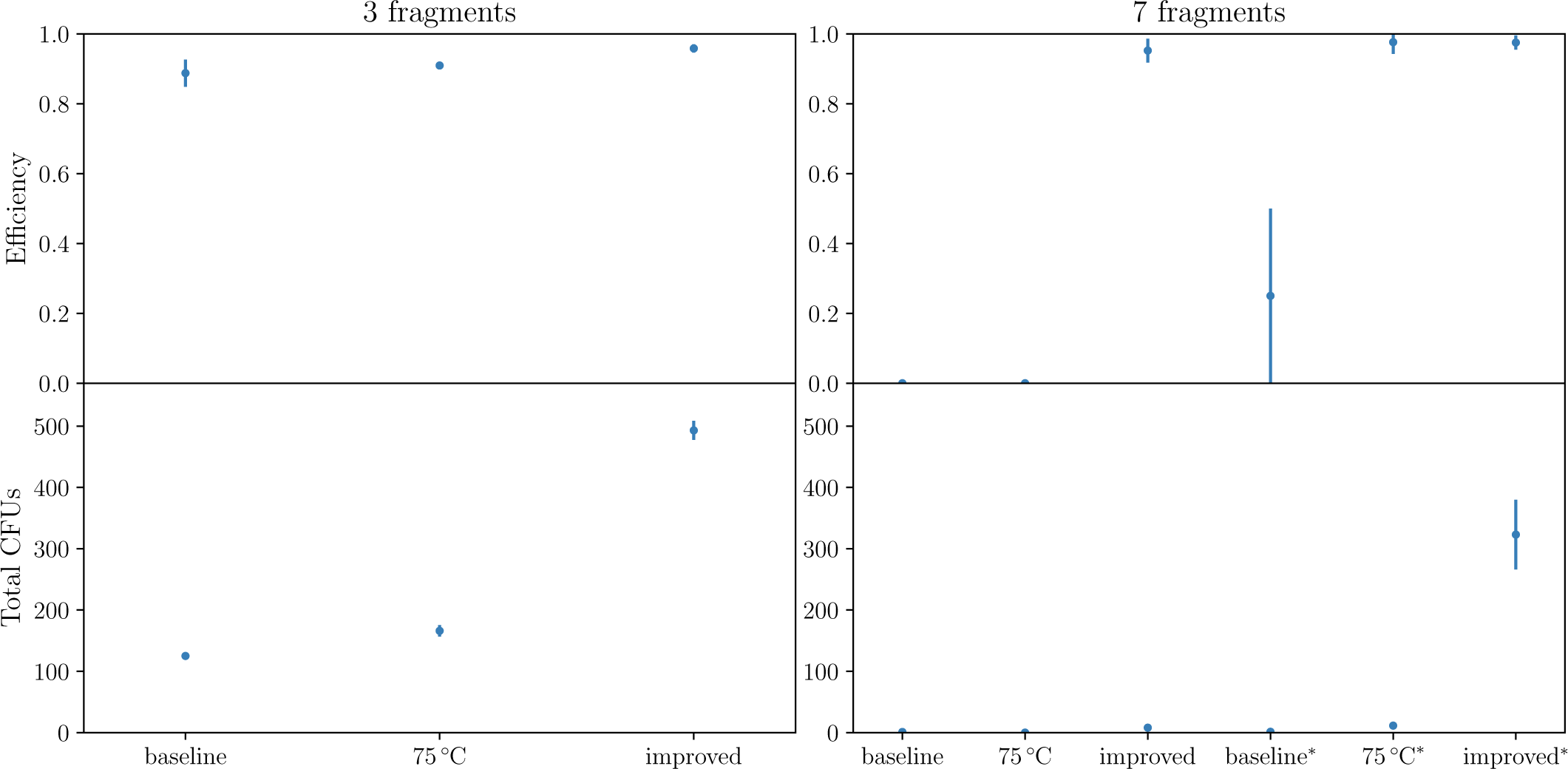
Results of the baseline, 75 °C and improved LCR-protocol for the validation plasmid 1 (“VP1”). For the seven-part split the same LCRs were transformed in chemical cells and electrocompetent cells (indicated by an *) due to no CFUs for the baseline and 75 °C protocol when chemical transformation was used. For all LCRs, 3 µL were transformed in 30 µL cells. As a negative control, the LCR-mix without BOs and without Ampligase^®^was used and resulted in no colonies (not shown). All LCRs were performed as triplicates. CFU: colony forming unit.

**Supplementary Figure 22.**
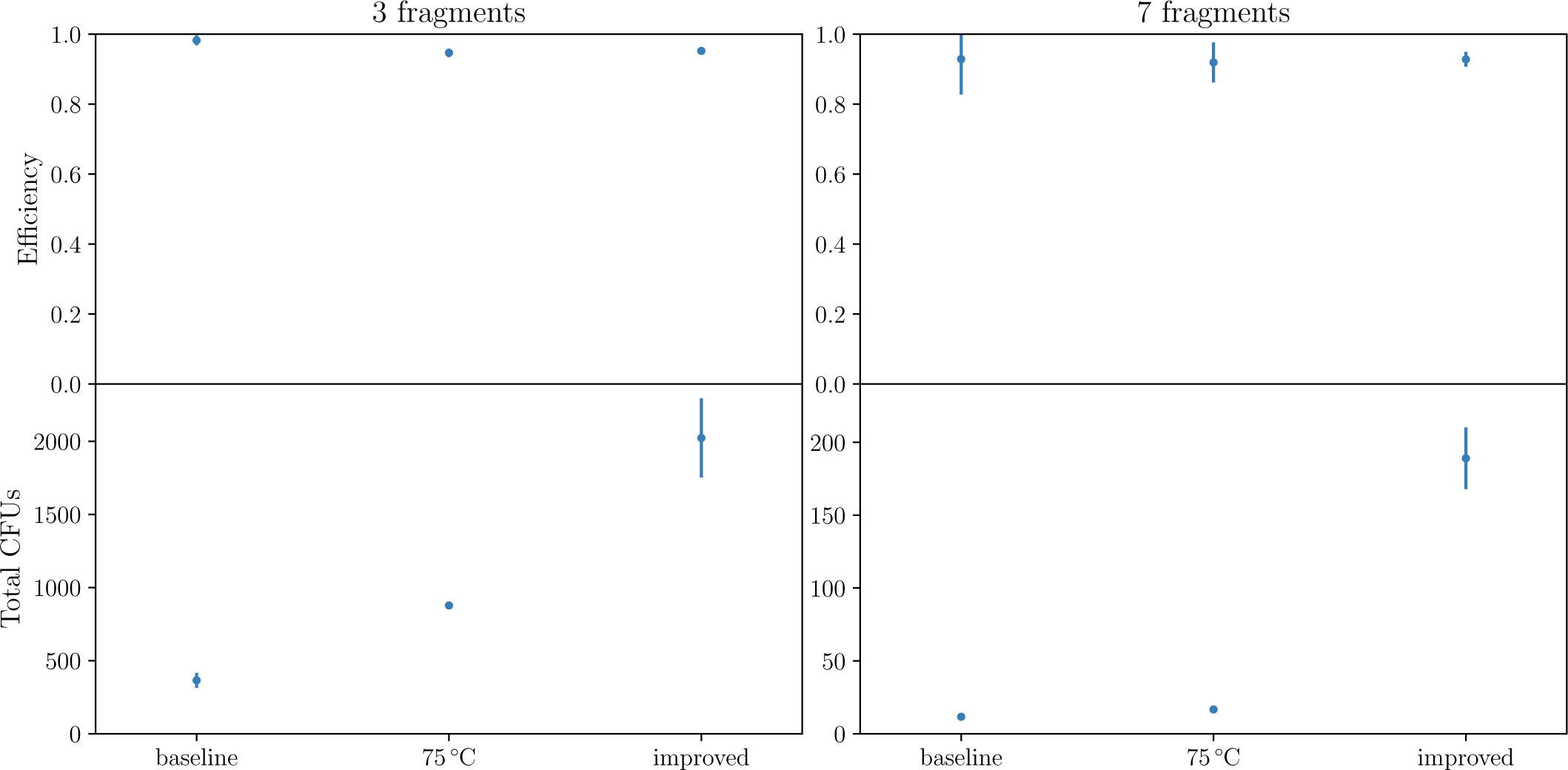
Results of the baseline, 75 °C and improved LCR-protocol for the validation plasmid 2 (“VP2”). All LCRs were transformed in chemical competent cells. For this, 3 µL were transformed in 30 µL cells. As a negative control, the LCR mix without BOs and without Ampligase^®^were used and resulted in no colonies (not shown). All LCRs were performed as triplicates. BO: bridging oligo, CFU: colony forming unit.

## Step-by-step protocol for the ligase cycling reaction

### Reagents

– amplification of DNA fragments:
  * proof-reading polymerase, no T/A overhangs
  * phosphorylated primers for the amplification of DNA fragments (use T4-polynucleotide kinase (T4-PNK; New England Biolabs, Ipswich, USA) and T4-PNK buffer (10×) or order as synthetically modified oligos
– LCR:
  * Ampligase^®^ thermostable DNA (Lucigen, Wisconsin, USA) ligase (Epicentre) and 10×-buffer
  * *aq. dest.*
  * competent cells

### Amplification of DNA fragments

– use <u>phosphorylated amplification primers</u> for the amplification of DNA fragments (inserts and backbone)
– phosphorylation of the amplification primers by T4-PNK (on ice):

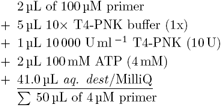
– incubate for 1 h at 37 °C and 20 min at 65 °C
– use 6.25 µL of each phosphorylated amplification primer in a 50 µL PCR (primer concentration in PCR: 500 nM)^1,2^
– <u>Recommended</u>: DpnI digestion of all parts before the purification
– store parts at 20 °C; prepare aliquots for multiple usage of the same part (loss of function due to repeated freeze-thaw cycles [1, 2])

### Bridging Oligo Design

For the LCRs the addition of DMSO/betaine is NOT recommended. The target-*T*_*m*_ of the BO design depends on the algorithms you are applying (1):

– SantaLucia 1998 (*T*_*m*_ calculation and salt correction [3])
  * target *T*_*m*_ ≈ 67.8 °C

or

– SantaLucia 1998 (*T*_*m*_ calculation [3]) and Owczarzy 2008 (salt correction [4])
  * target *T*_*m*_ ≈ 65.2 °C

or

– `Geneious` (www.geneious.com, [5]) utilizing `Primer3` [6]:
  * a Geneious-plugin for the design of bridging oligos is available at www.gitlab.com/kabischlab.de/lcr-publication-synthetic-biology to use Geneious without the plugin:
  * restrictions in adjusting parameters for DNA-part concentration (only “Oligo” can be adjusted)
  * to get correct melting temperatures use 114 nM for the input of “Oligo”
  * then use SantaLucia 1998 [3] for the *T*_*m*_ calculation and salt correction (Owczarzy 2008 [4] is not available for the calculations in `Geneious` versions≤ 11.0.5)
  * target *T*_*m*_ ≈ 67.8 °C

**Figure 1:**
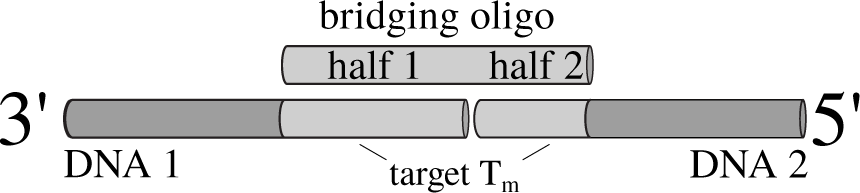
Scheme of the bridging oligo design. Half 1: 100% complementary to the 5-end of the first DNA part. Half 2: 100% complementary to the 3-end of the second DNA part.

### Ligation

– use inserts:vector ratio of 10:1 (use 0.3 nM of the backbone) to reduce the background (religation of the backbone occurs due to the phosphorylation of all parts)
– for 25 µl (in brackets: final concentration): DNA inserts (3 nM of each fragment)

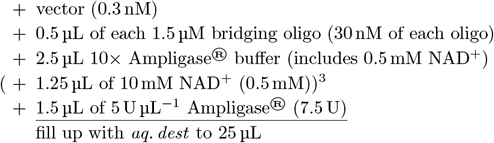
– use low-profile tubes or low-profile PCR plates (96-well) in a cycler for volumes of less than 10 µL
– for more than 25 cycles of small volumes (≤10 µL): use 384-well plates
– start cycling (Figure 2)

**Figure 2:**
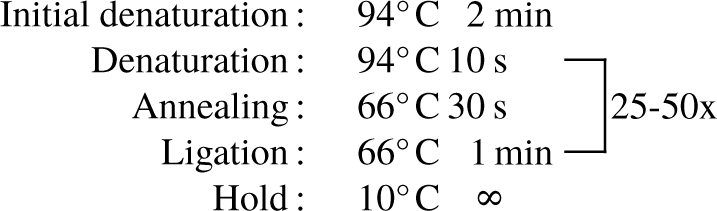
Cycling parameters for the LCR.

– store at −20 °C or directly use for transformation of chemically- or electrocompetent cells or use LCR-mix as template in a PCR (1 µl of 1:100 dilution works in general)

### Notes

1. This is the recommended primer concentration for the Q5^®^ High-Fidelity Polymerase (New England Biolabs, Ipswich, USA). Probably, the concentration has to be adjusted for your polymerase. For this please ask your supplier.
2. For some PCRs the sum of the forward and reverse primers 12.5 µL in a total PCR-volume of 50 µL can result in no product. In those cases, a dialysis of the phosphorylated primers for 20 min with *aq. dest.* and a floating membrane (Merck-Millipore™Membrane Filter, 0.025 µm; order-#: VSWP02500) is helpful.
3. Self-made 10× -buffer without NAD^+^ is recommended. NAD^+^ is sensitive to freeze-thaw cycles, long-term storage and light exposure. Prepare a stock solution of NAD^+^ and store in aliquots at −80 °C up to 6 months. 10× Ampligase^®^ buffer: 200 mM TRIS-HCl (pH 8.3), 250 mM KCl, 100 mM MgCl_2_, 0.1% Triton X-100.

---

**Supplementary Table 6:**
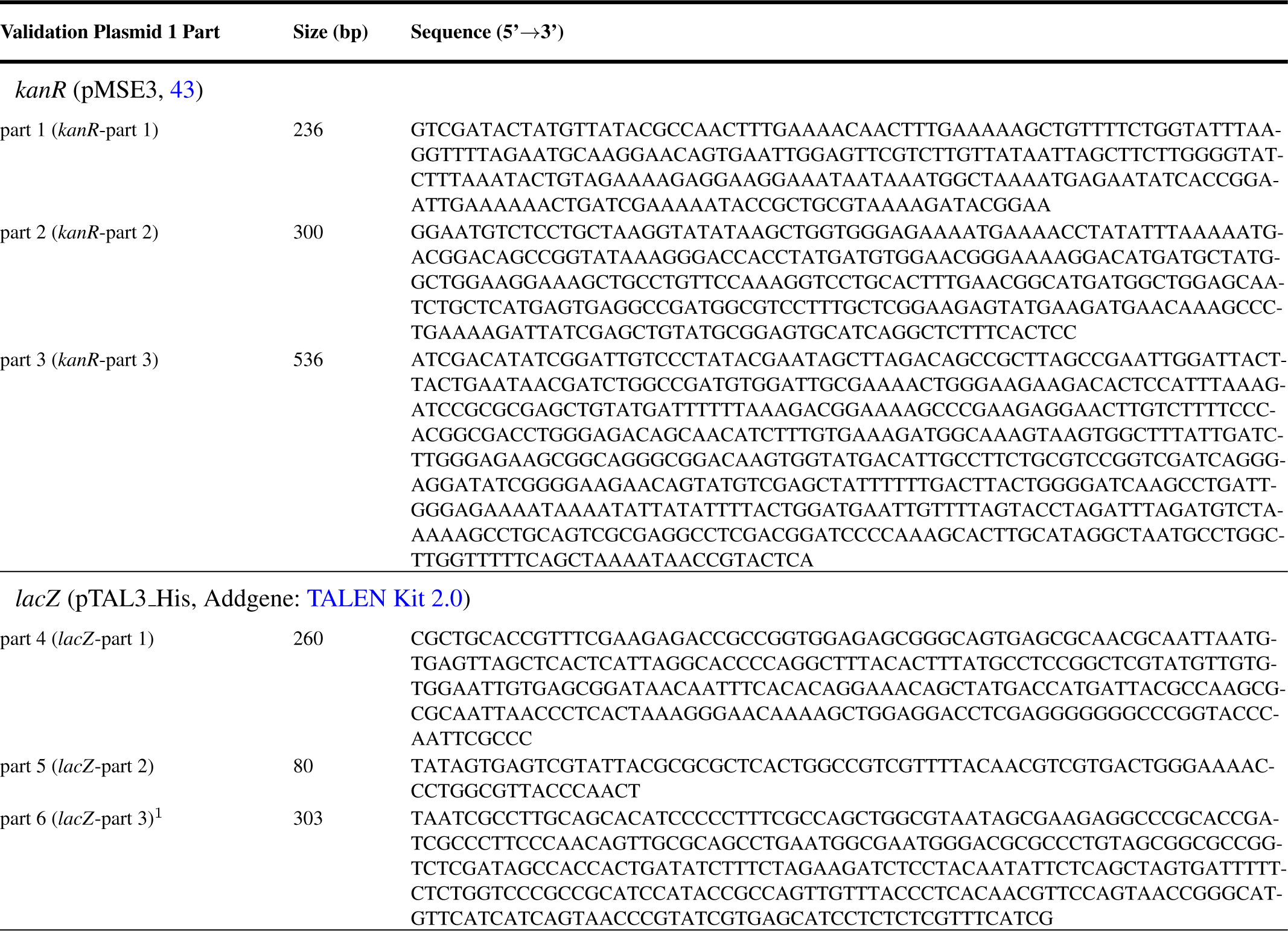
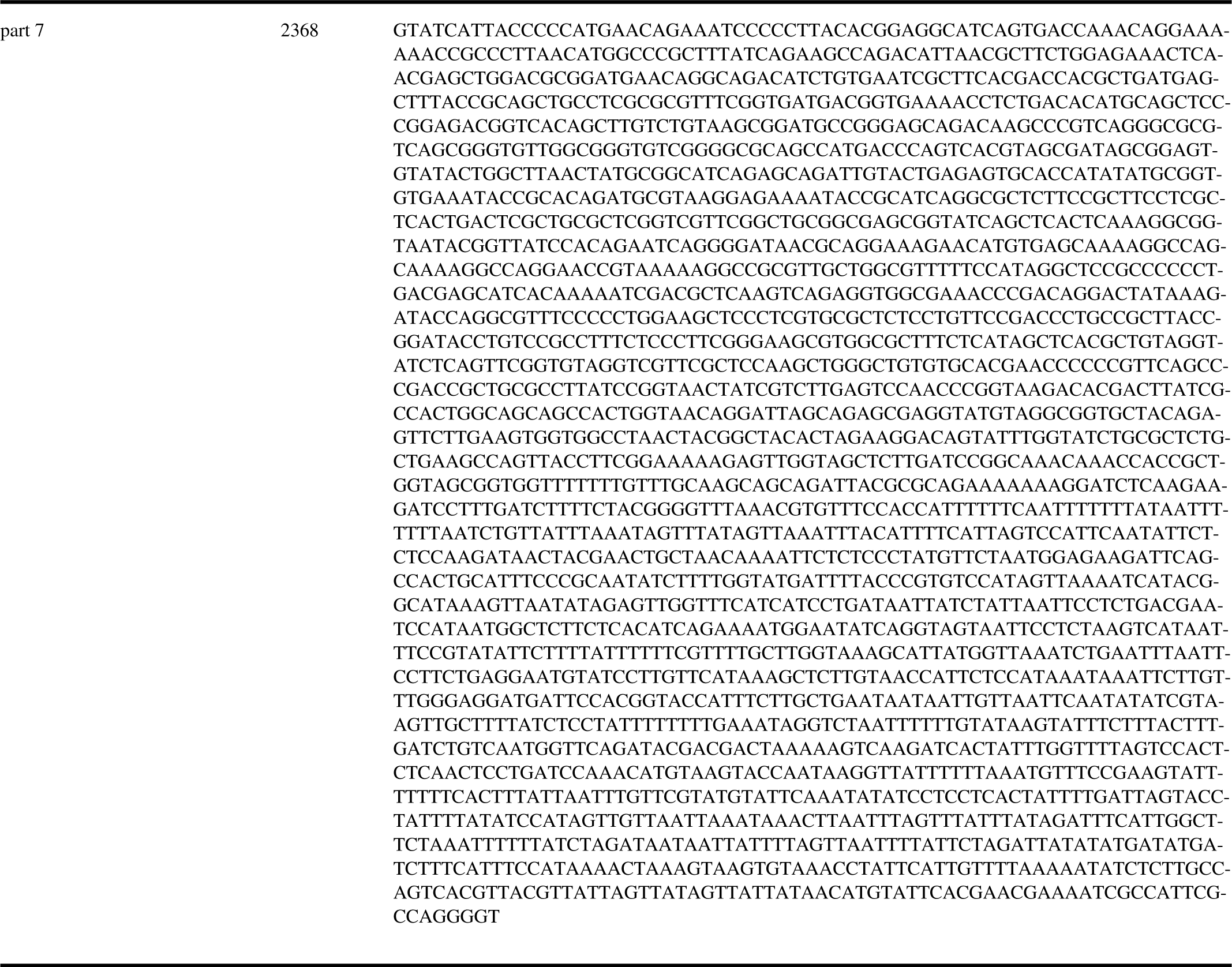
Sequences of the seven-part split of the validation plasmid 1 (Supplementary Figure 18). For the three-part split parts 1-3, 4-6 and 7 were used. The bridging oligos were designed by concatenating subsequences of these parts. It was not necessary to use the reverse complement for the design because the experiments contained both strands of these parts. ^1^ Part 6 begins in the *lacZ* of a plasmid derived from the TALEN Kit 2.0 from Addgene (42) and ends in the plasmid derived from Wenzel et al. (41). *kanR*: kanamycin resistance gene, *lacZ*: *β*-galactosidase.

**Supplementary Table 7:**
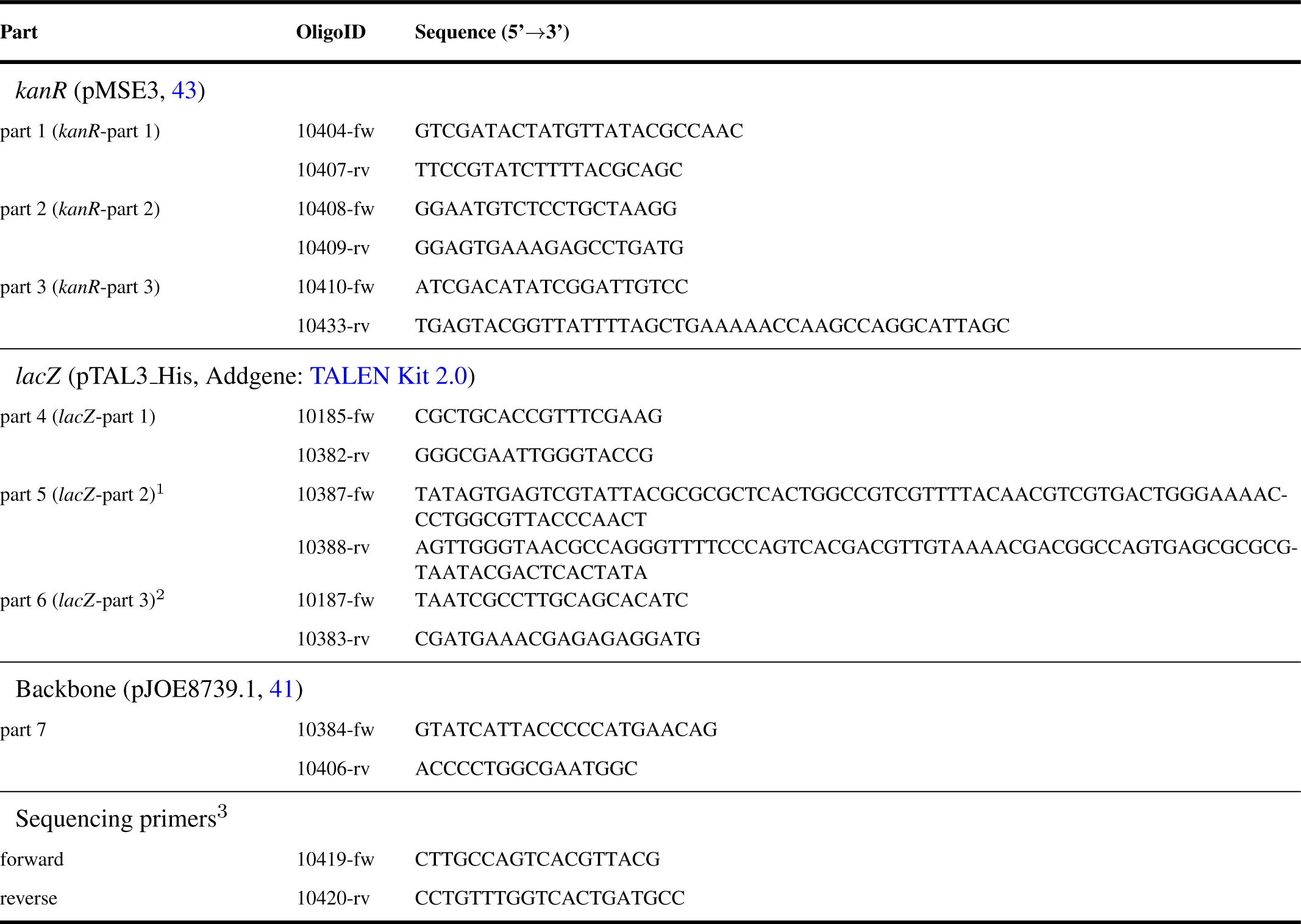
Oligonucleotides for the amplification of all validation plasmid 1 parts (shown in Supplementary Figure 18 and Supplementary Table 6). For the amplification of the parts of the three-part split the oligonucleotides of parts 1 (fw)+3 (rv), 4 (fw)+6 (rv) and 7 (fw and rv) were used. ^1^ The complete part was ordered as two oligonucleotides, which were phosphorylated and annealed to double stranded DNA before the LCR (described in the methods). ^2^ Part 6 begins in the *lacZ* of the plasmid derived from the TALEN Kit 2.0 from Addgene (42) and ends in the plasmid derived from Wenzel et al. (41). ^3^ For sequencing from the vector into the inserts. fw: forward direction, *kanR*: kanamycin resistance gene, *lacZ*: *β*-galactosidase, rv: reverse direction.

**Supplementary Table 8:**
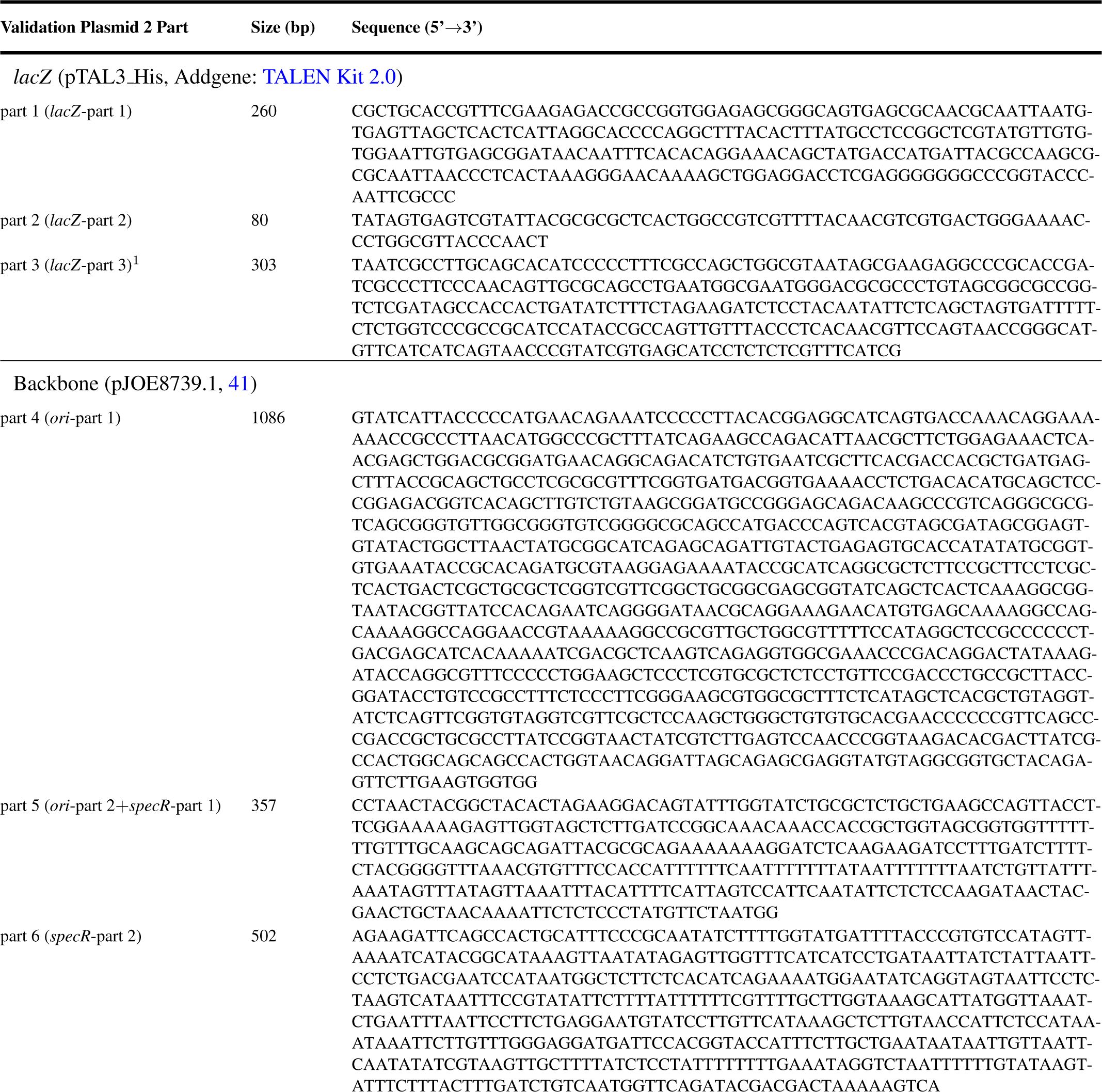
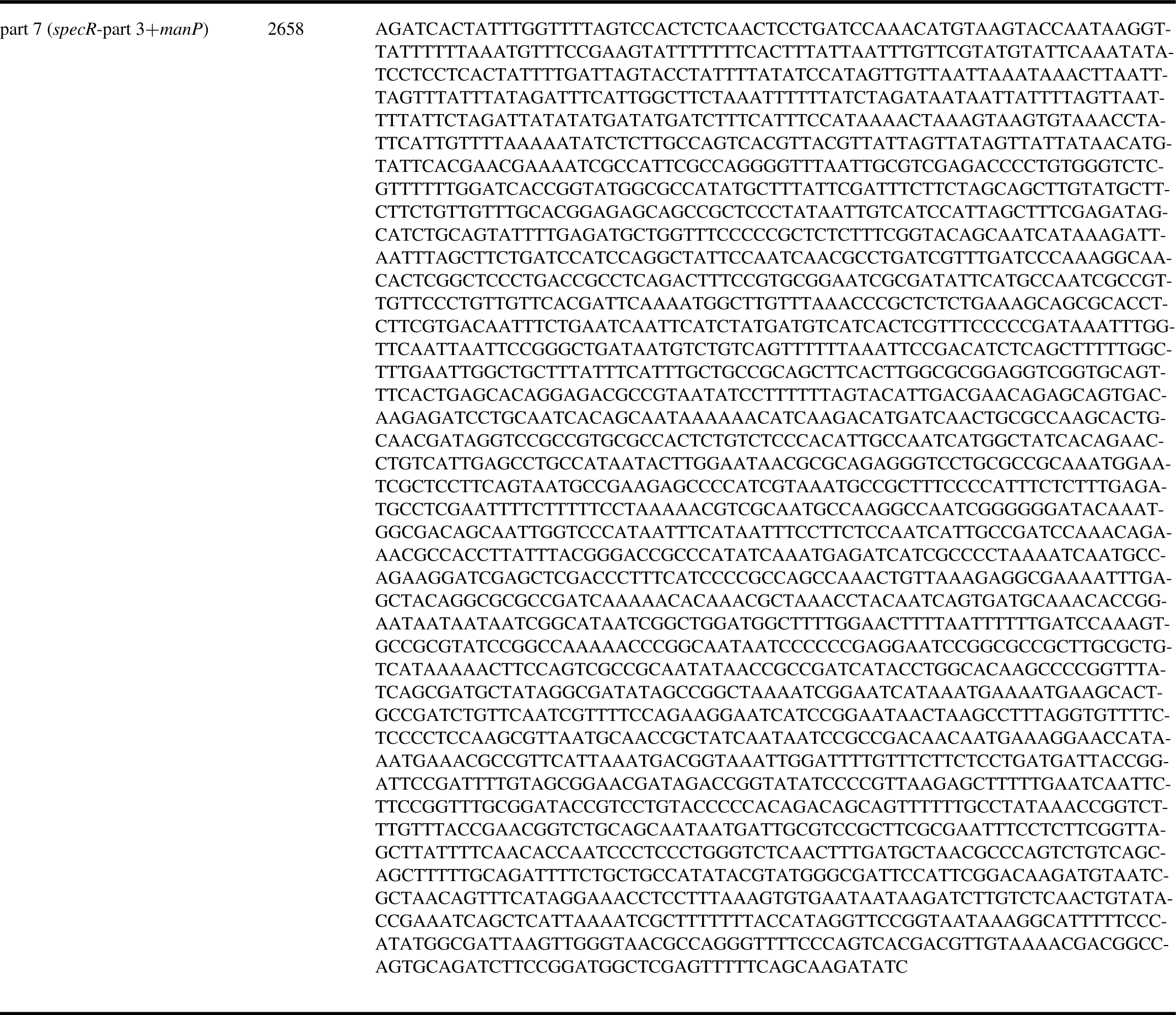
Sequences of the seven-part split of the validation plasmid 2 (Supplementary Figure 19). For the three-part split parts 2-4, 5+6 and 7+1 were used. The bridging oligos were designed by concatenating subsequences of these parts. It was not necessary to use the reverse complement for the design because the experiments contained both strands of these parts. ^1^ Part 3 begins in the *lacZ* of the plasmid derived from the TALEN Kit 2.0 from Addgene (42) and ends in the plasmid derived from Wenzel et al. (41). *kanR*: kanamycin resistance gene, *lacZ*: *β*-galactosidase.

**Supplementary Table 9:**
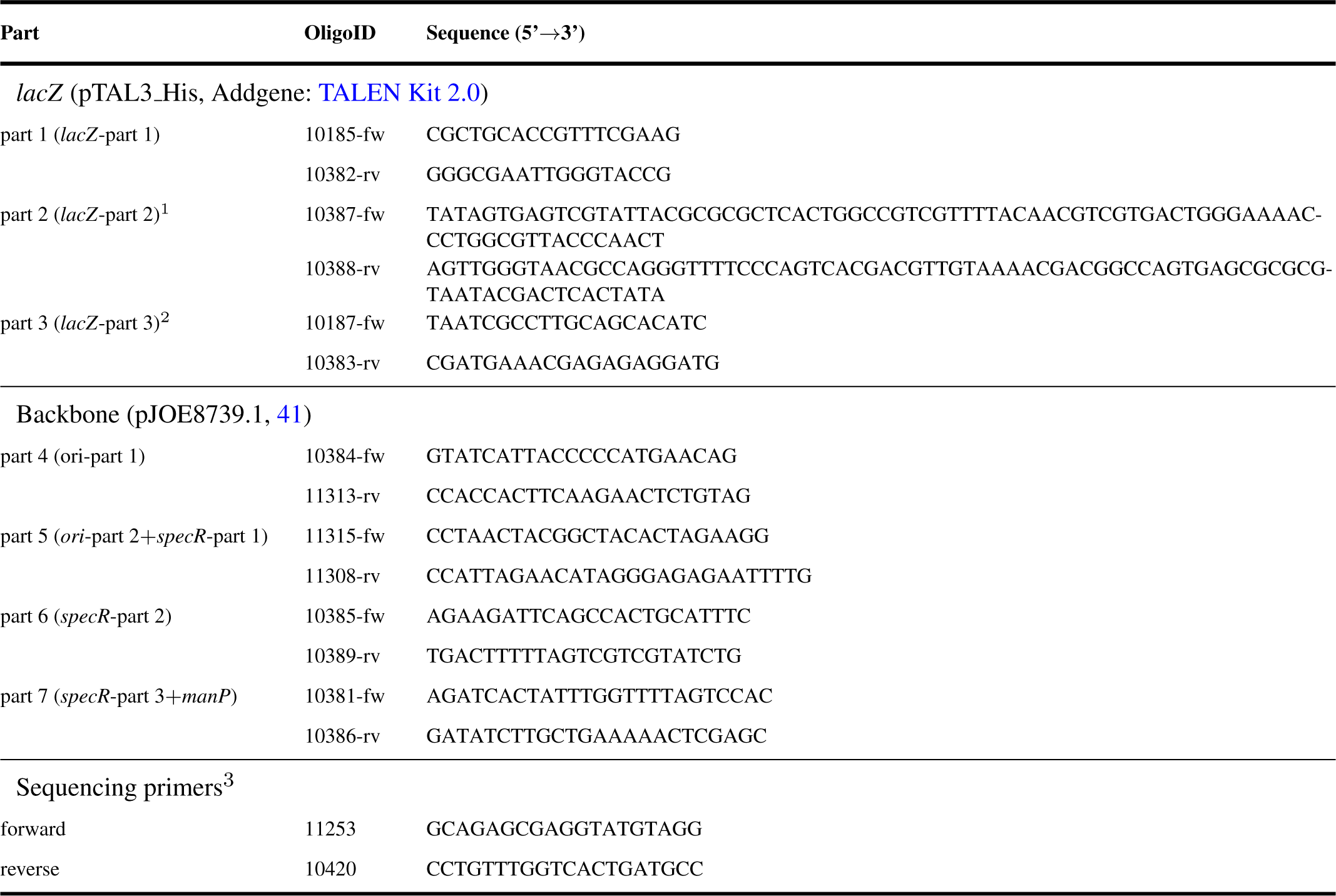
Oligonucleotides for the amplification of all validation plasmid 2 parts (shown in Supplementary Figure 19 and Supplementary Table 9). For the amplification of the parts of the three-part split the oligonucleotides of parts 2 (fw)+4 (rv), 5 (fw)+6 (rv) and 7 (fw)+1 (rv) were used. ^1^ The complete part was ordered as two oligonucleotides, which were phosphorylated and annealed to double stranded DNA before the LCR (described in the methods). ^2^ Part 3 begins in the *lacZ* of the plasmid derived from the TALEN Kit 2.0 from Addgene (42) and ends in the plasmid derived from Wenzel et al. (41). ^3^ For sequencing from the vector into the inserts. fw: forward direction, *ori*: origin of replication, *lacZ*: *β*-galactosidase, rv: reverse direction, *specR*: spectinomycin resistance gene.

**Supplementary Table 10:**
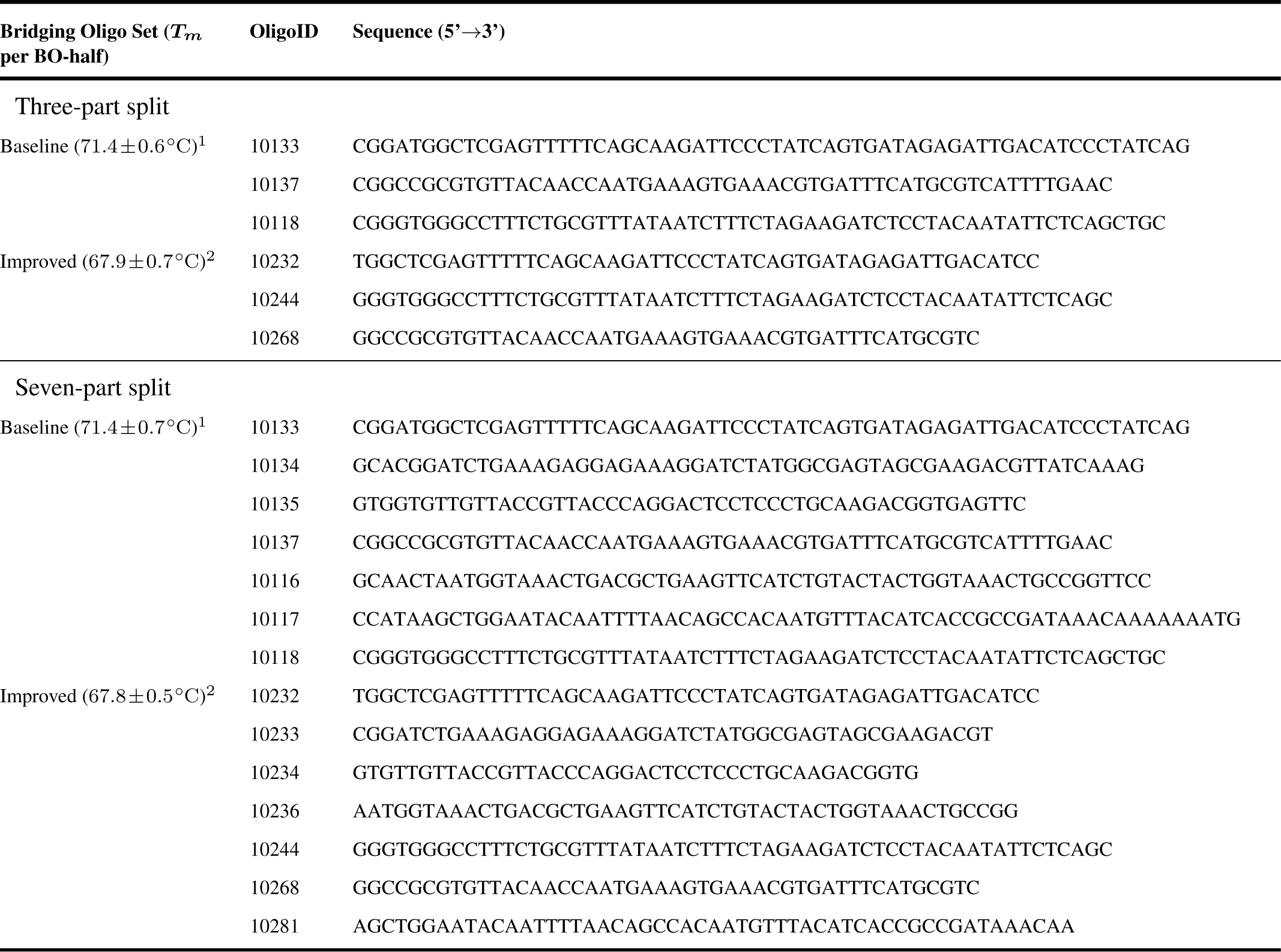
Bridging oligonucleotides for the assembly of the three-part and seven-part toy-plasmid consisting of *mRFP1, sfGFP* and pJET1.2/blunt by using the baseline and improved protocol (results and protocols shown in Figure 4). The BO-set for the baseline LCR consists of BOs of the manual set shown in Supplementary Table 3. The BO-set for the improved LCR consists of BOs of the “67.8 °C”-set of the gradient-LCR (Supplementary Table 4). All melting temperatures (*T*_*m*_s) presented here are calculated for each BO-half using the formula of SantaLucia (20) for the *T*_*m*_-calculation and the salt correction. ^1^ Bridging oligos with a target T_*m*_ of 70.0 °C for each half, with 8 %v*/*v DMSO and 0.45 M betaine and the experimental annealing temperature of 55 °C. ^2^ Bridging oligos with a target T_*m*_ of 67.8 °C for each half, <u>without</u> DMSO and betaine and the experimental annealing temperature of 66 °C. *T*_*m*_: melting temperature of a BO-half.

**Supplementary Table 11:**
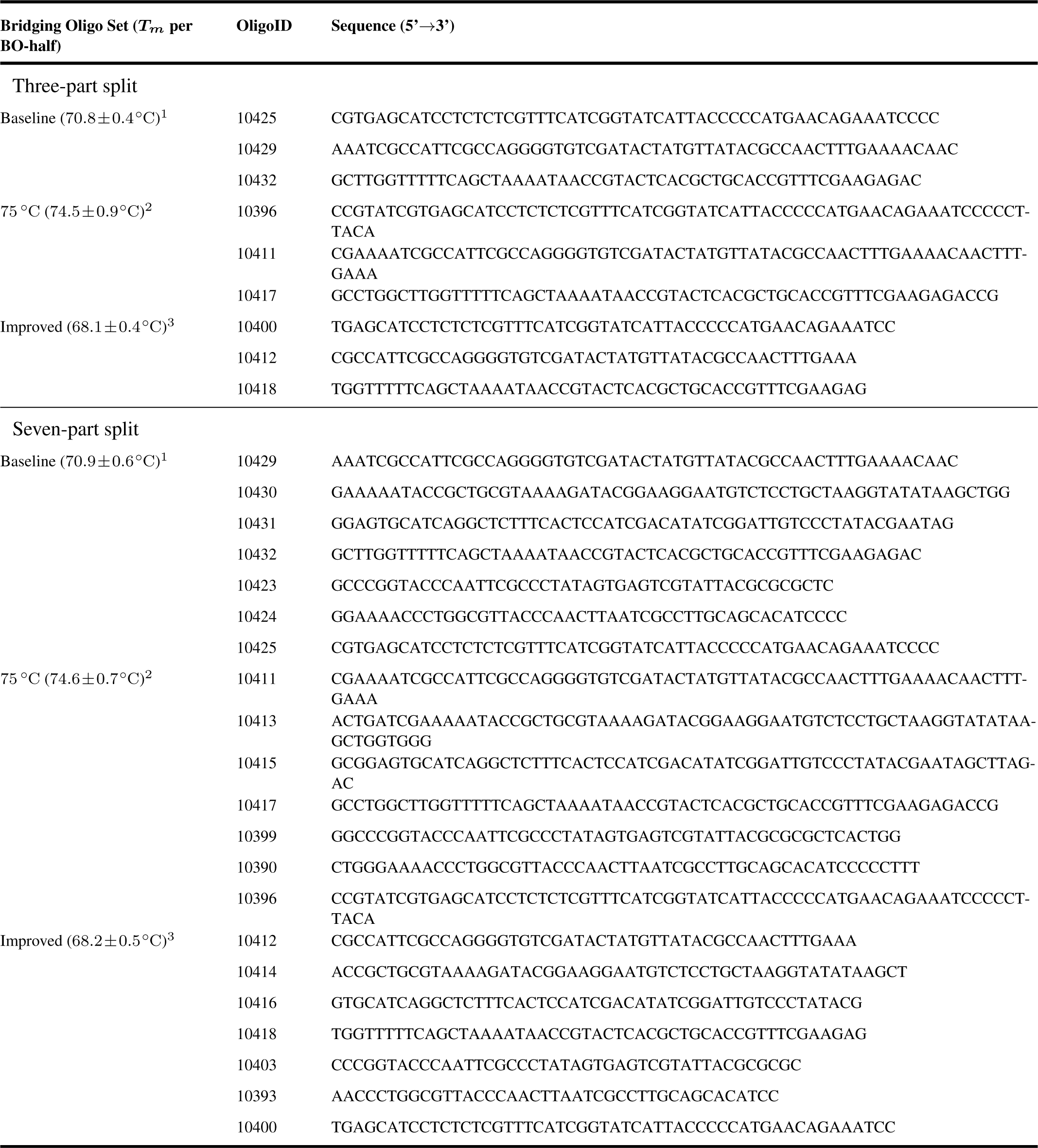
Bridging oligos for the assembly of the validation plasmid 1 (Supplementary Figure 18; results shown in Figure 4). All melting temperatures (*T*_*m*_s) presented here are calculated for each BO-half using the formula of SantaLucia (20) for the *T*_*m*_-calculation and the salt correction. ^1^ Bridging oligos with a target T_*m*_ of 70.0 °C for each half, with 8 %v*/*v DMSO and 0.45 M betaine and the experimental annealing temperature of 55 °C. ^2^ Bridging oligos with a target T_*m*_ of 74.8 °C for each half, with 8 %v*/*v DMSO and 0.45 M betaine and the experimental annealing temperature of 55 °C. ^3^ Bridging oligos with a target T_*m*_ of 67.8 °C for each half, <u>without</u> DMSO and betaine and the experimental annealing temperature of 66 °C. DMSO: dimethyl sulfoxide, *T*_*m*_: melting temperature of a BO-half, w/o: without.

**Supplementary Table 12:**
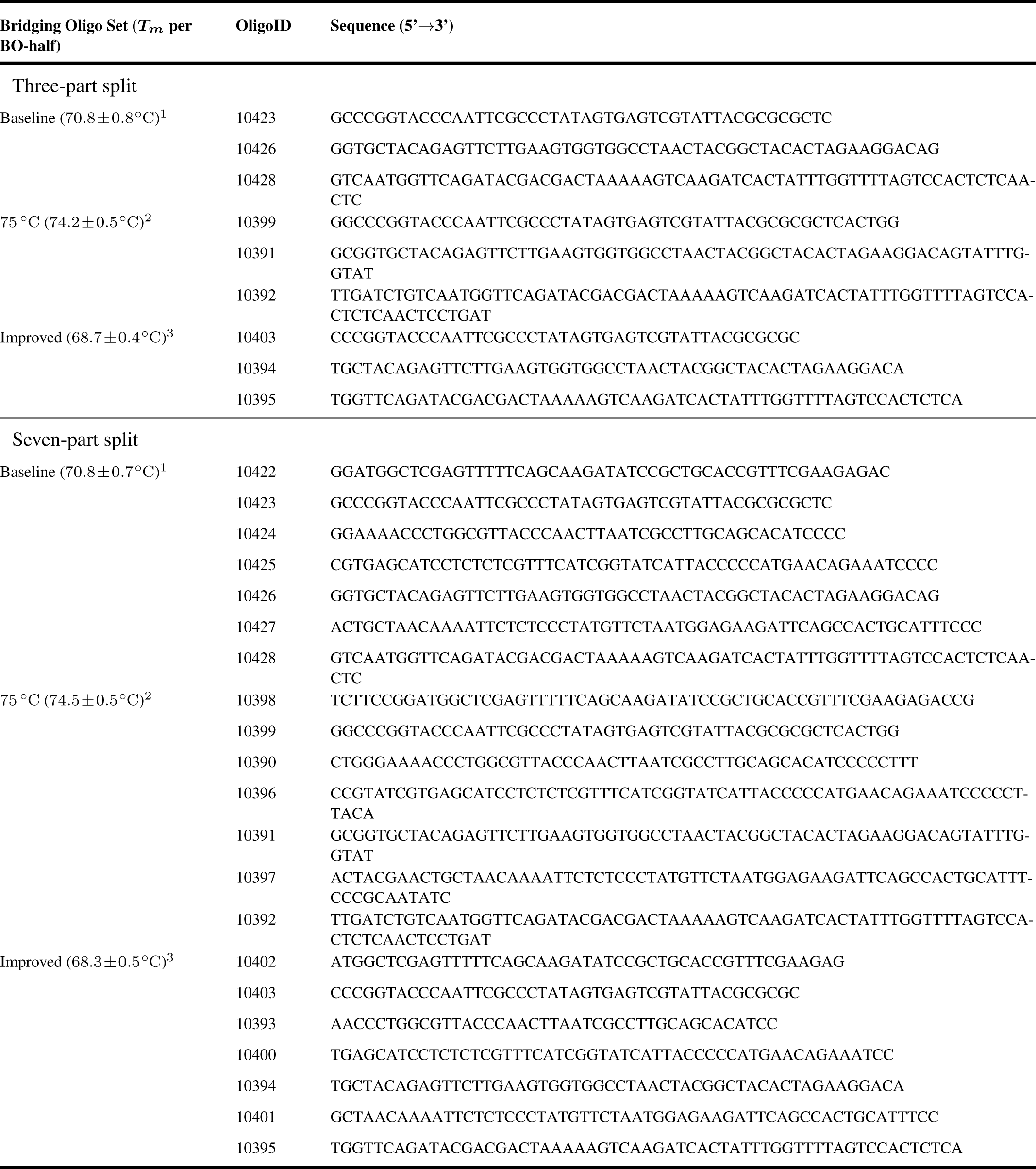
Bridging oligos for the assembly of the validation plasmid 2 (Supplementary Figure 19; results shown in Figure 4). All melting temperatures (*T*_*m*_s) presented here are calculated for each BO-half using the formula of SantaLucia (20) for the *T*_*m*_-calculation and the salt correction. ^1^ Bridging oligos with a target T_*m*_ of 70.0 °C for each half, with 8 %v*/*v DMSO and 0.45 M betaine and the experimental annealing temperature of 55 °C. ^2^ Bridging oligos with a target T_*m*_ of 74.8 °C for each half, with 8 %v*/*v DMSO and 0.45 M betaine and the experimental annealing temperature of 55 °C. ^3^ Bridging oligos with a target T_*m*_ of 67.8 °C for each half, <u>without</u> DMSO and betaine and the experimental annealing temperature of 66 °C. DMSO: dimethyl sulfoxide, *T*_*m*_: melting temperature of a BO-half.

## GENBANK FILES

### Toy-model plasmid

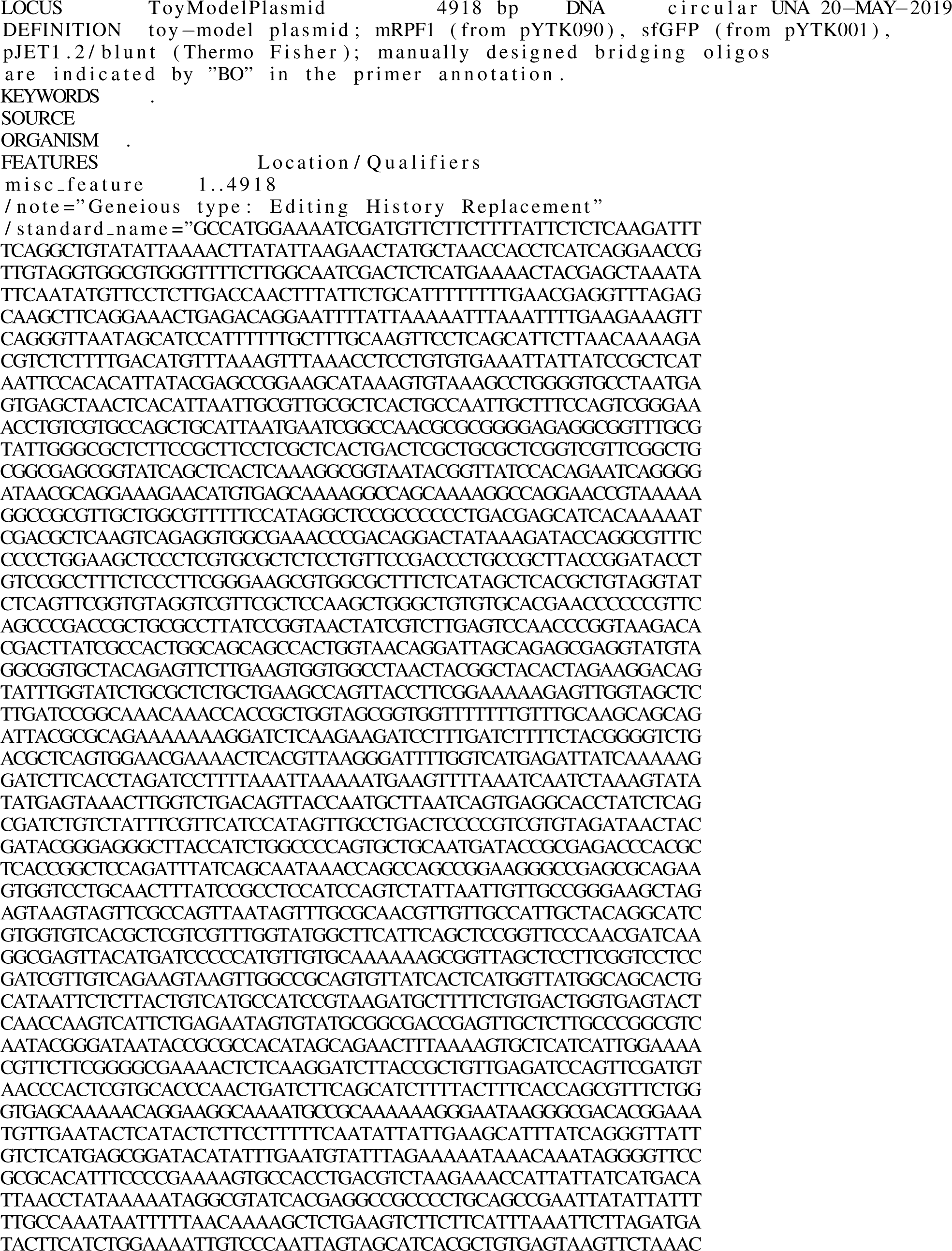

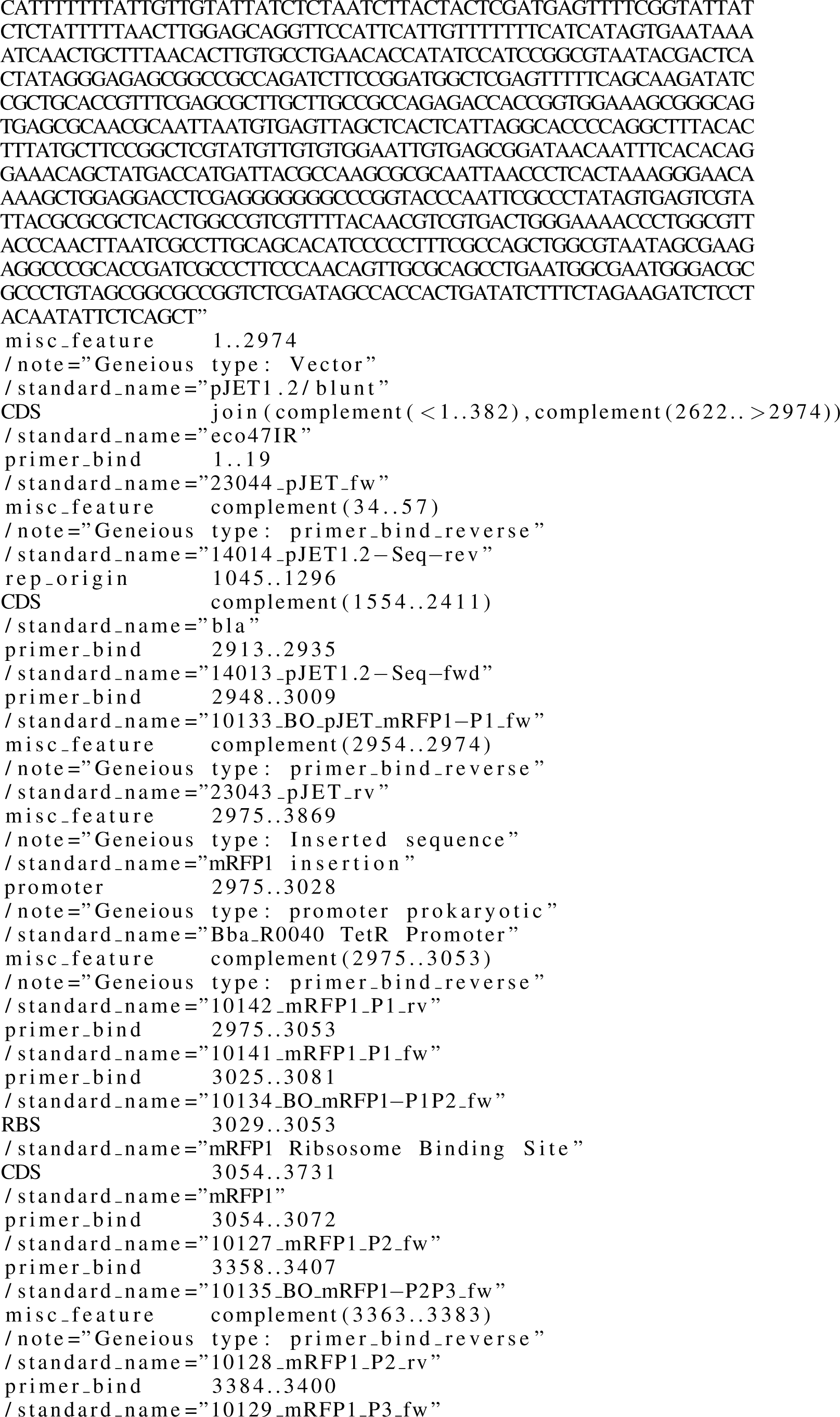

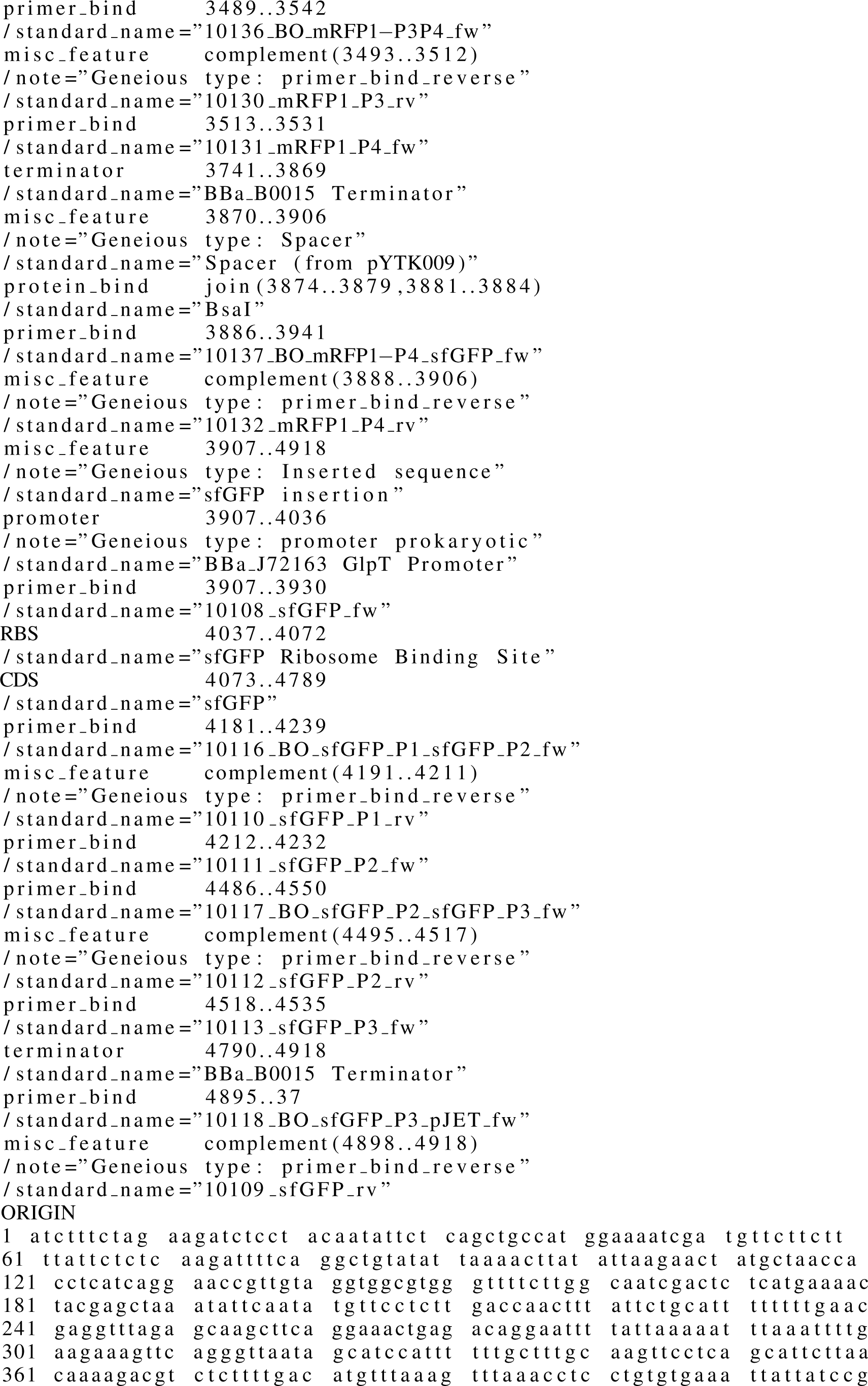

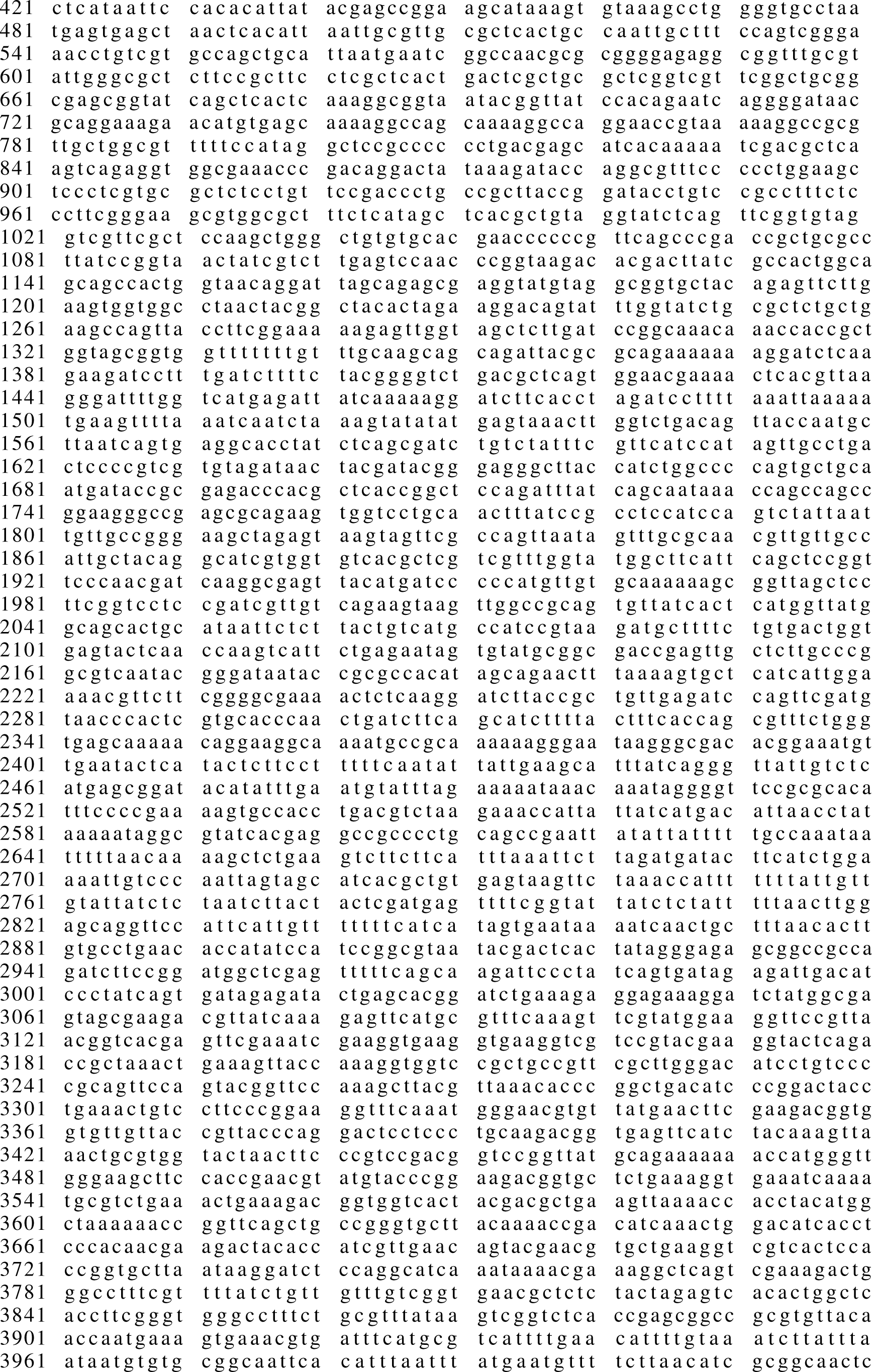

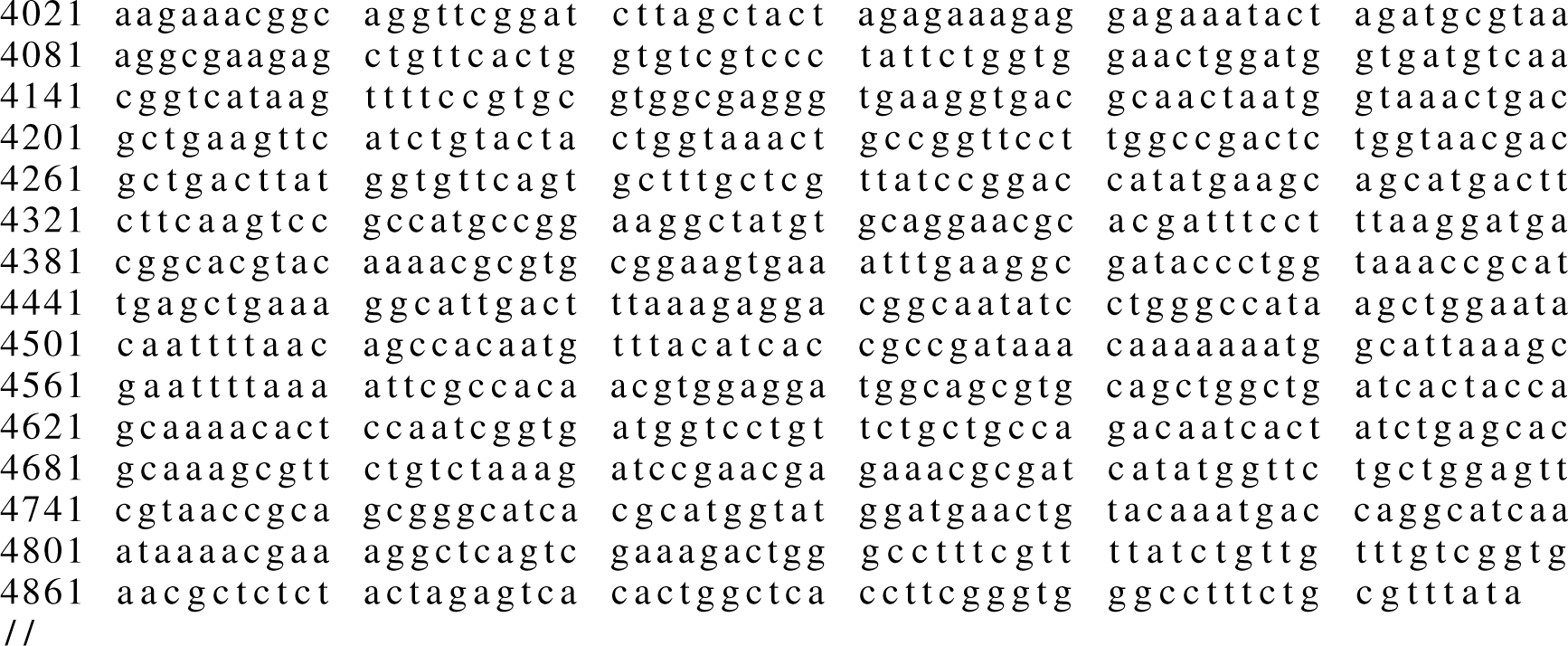

### Validation plasmid 1

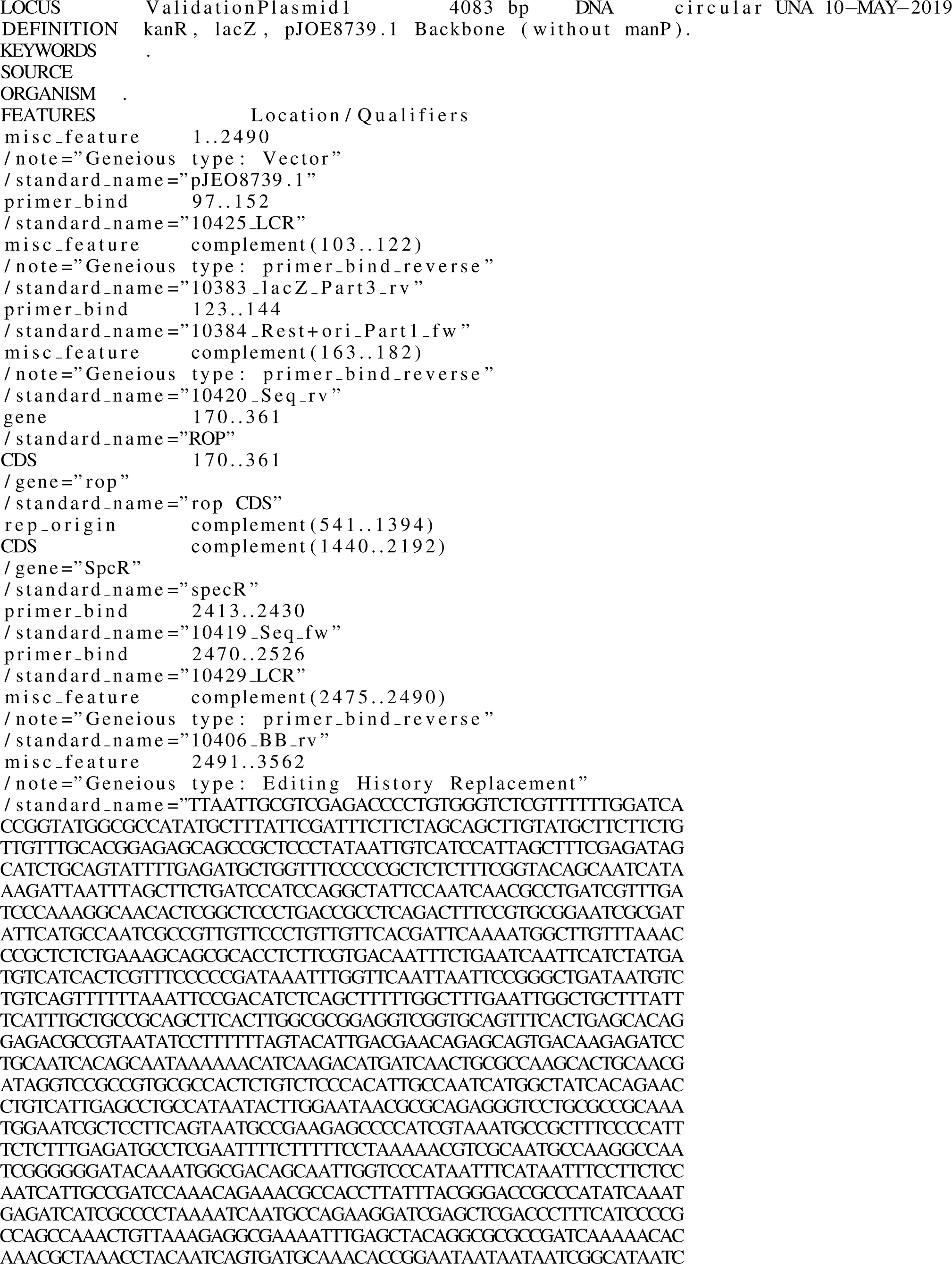

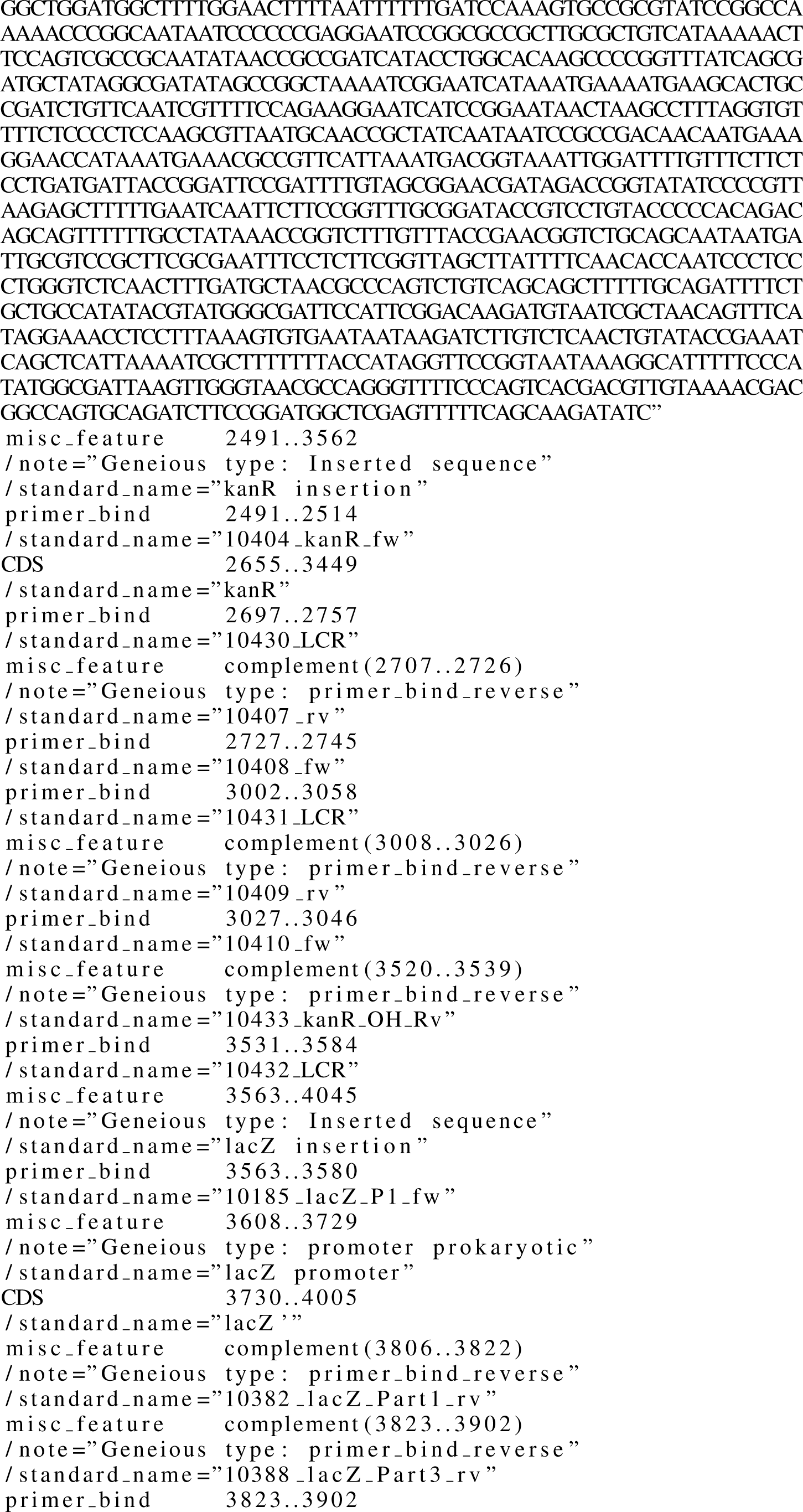

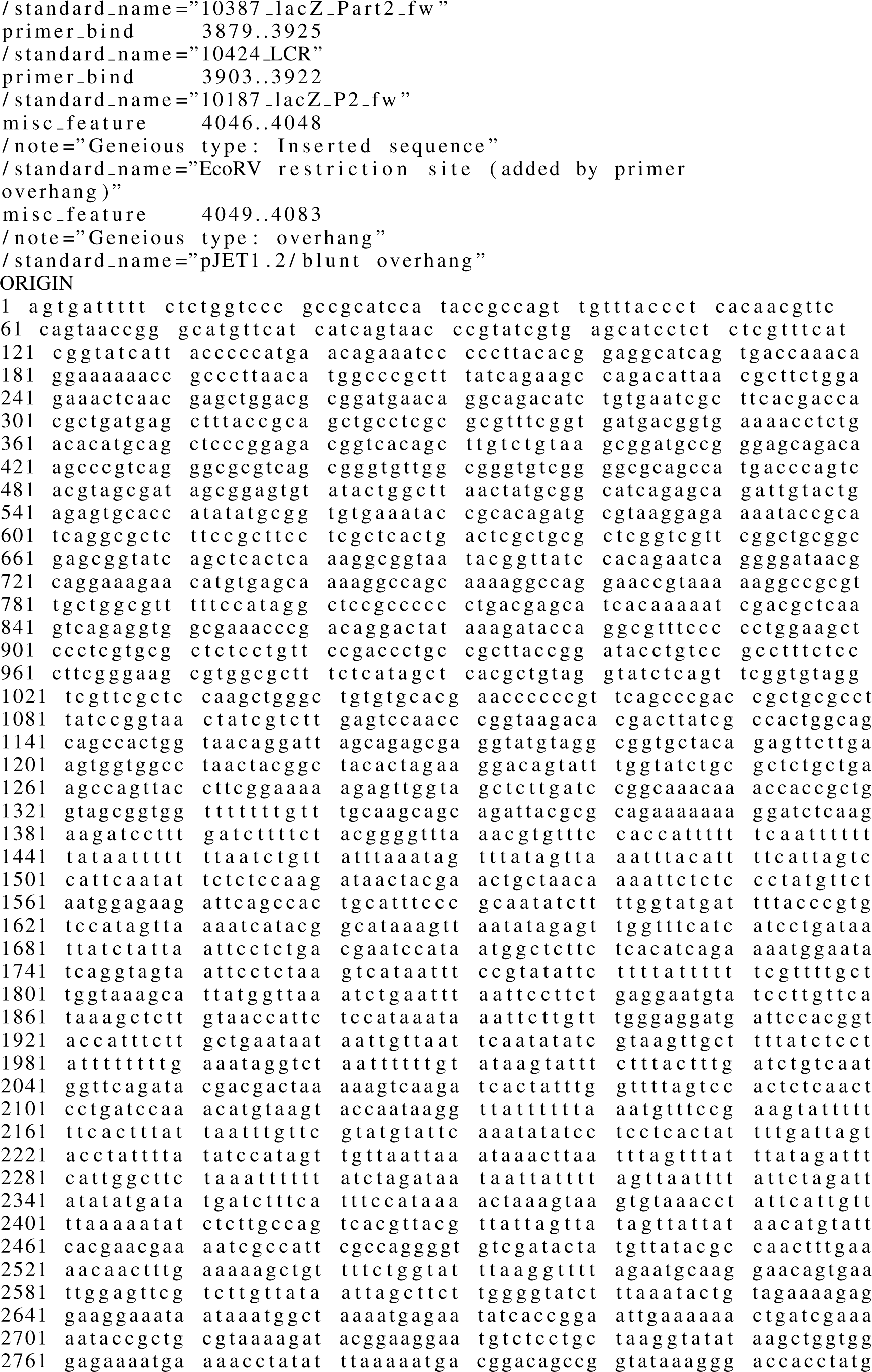

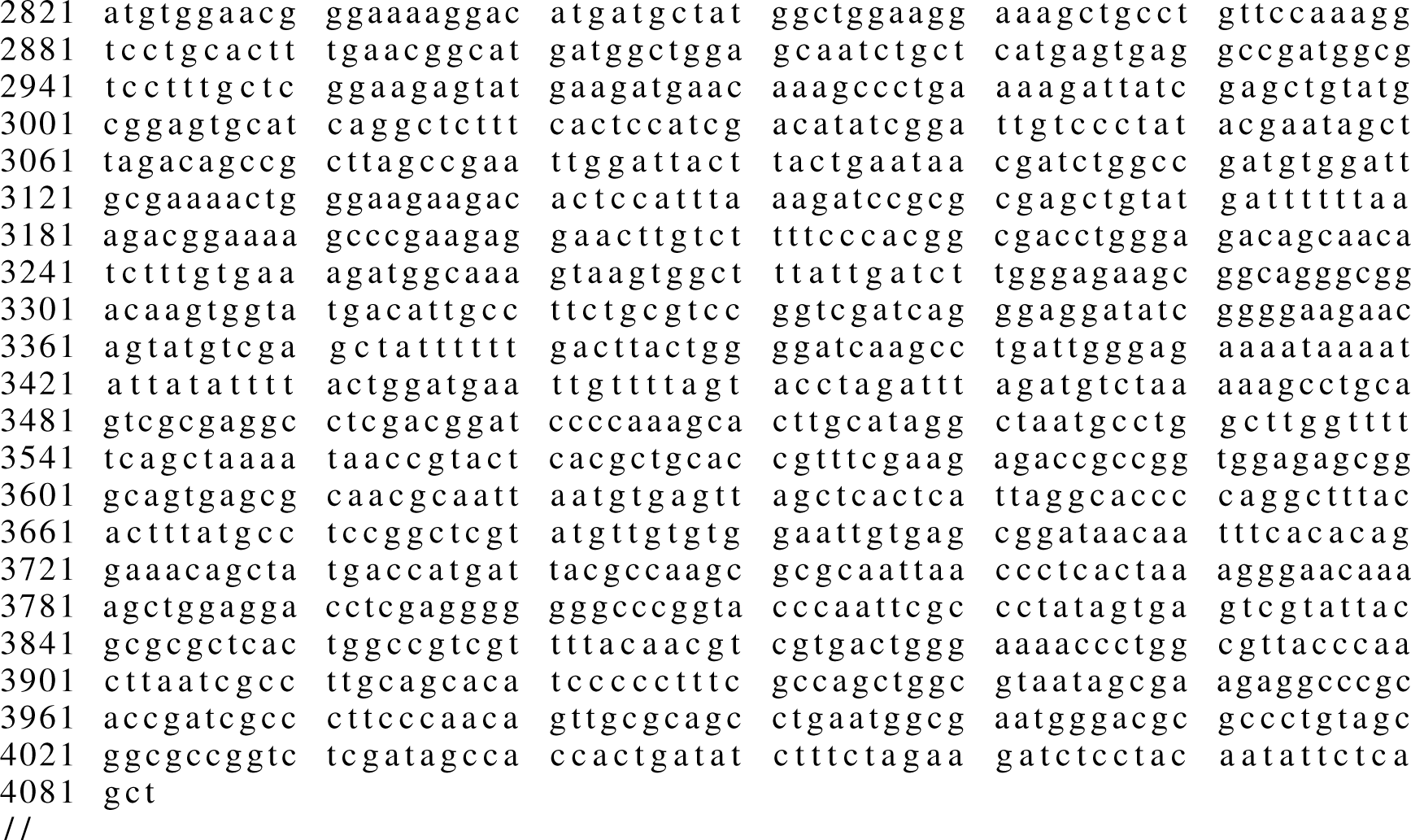

### Validation plasmid 2

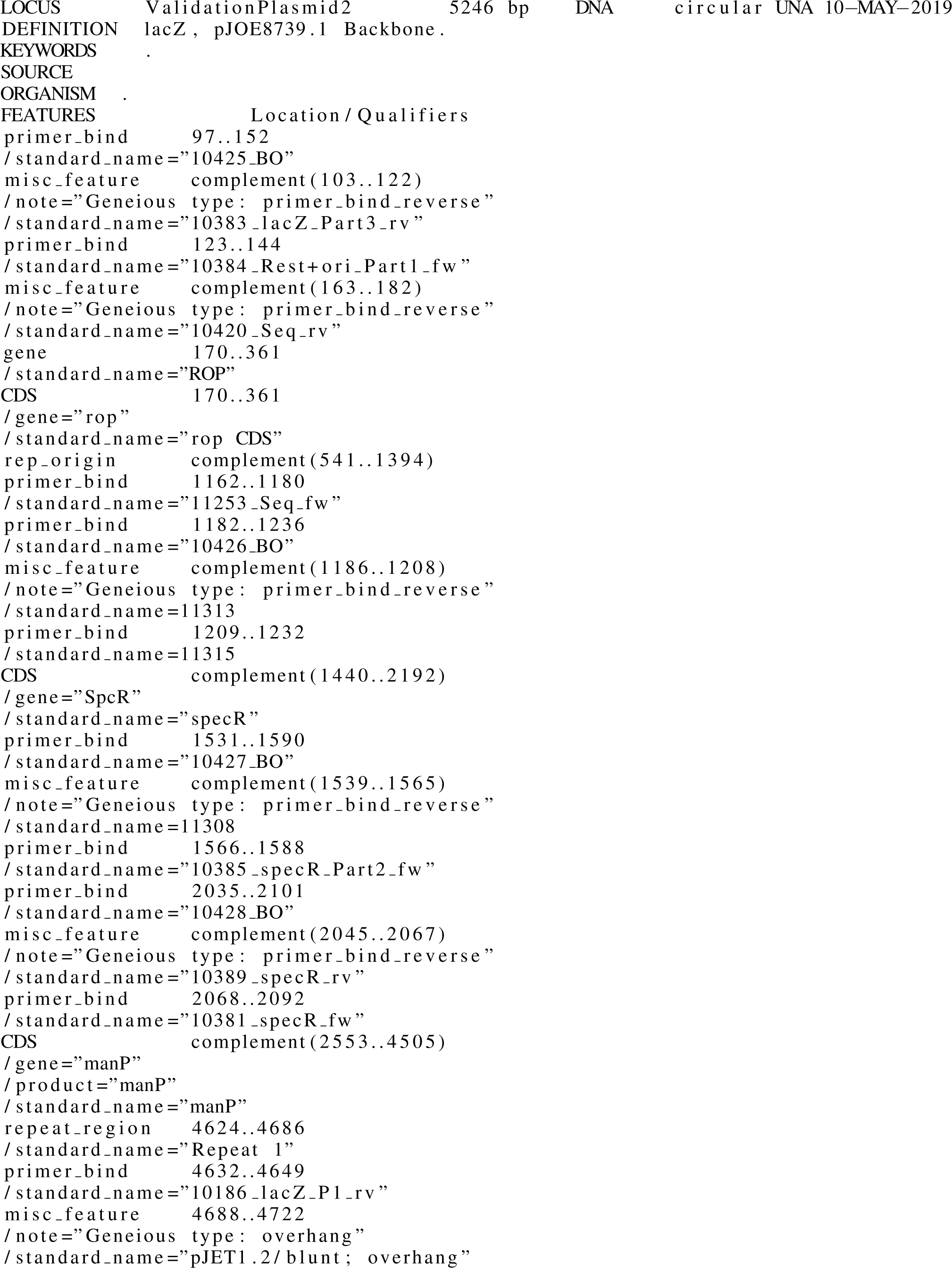

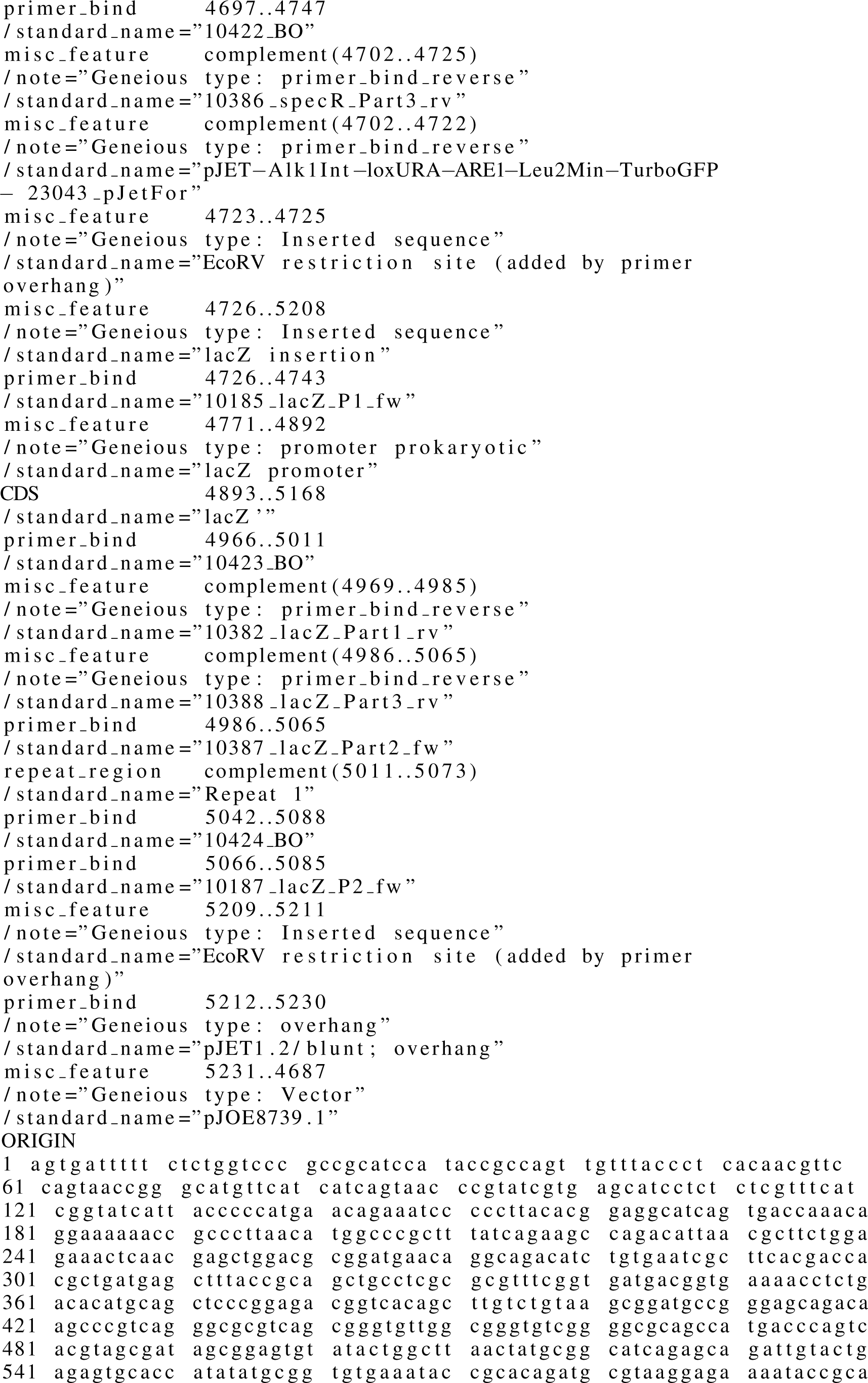

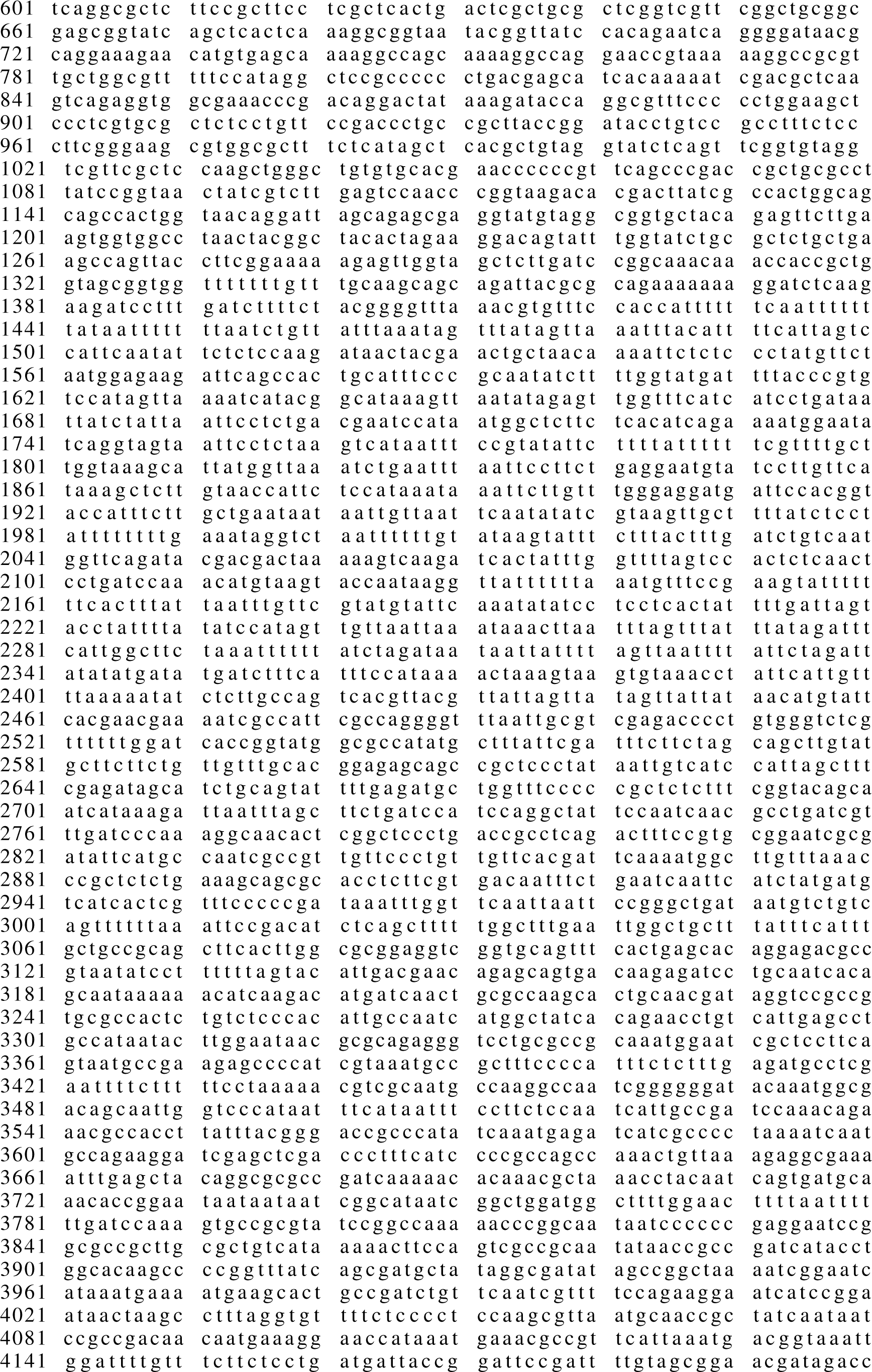

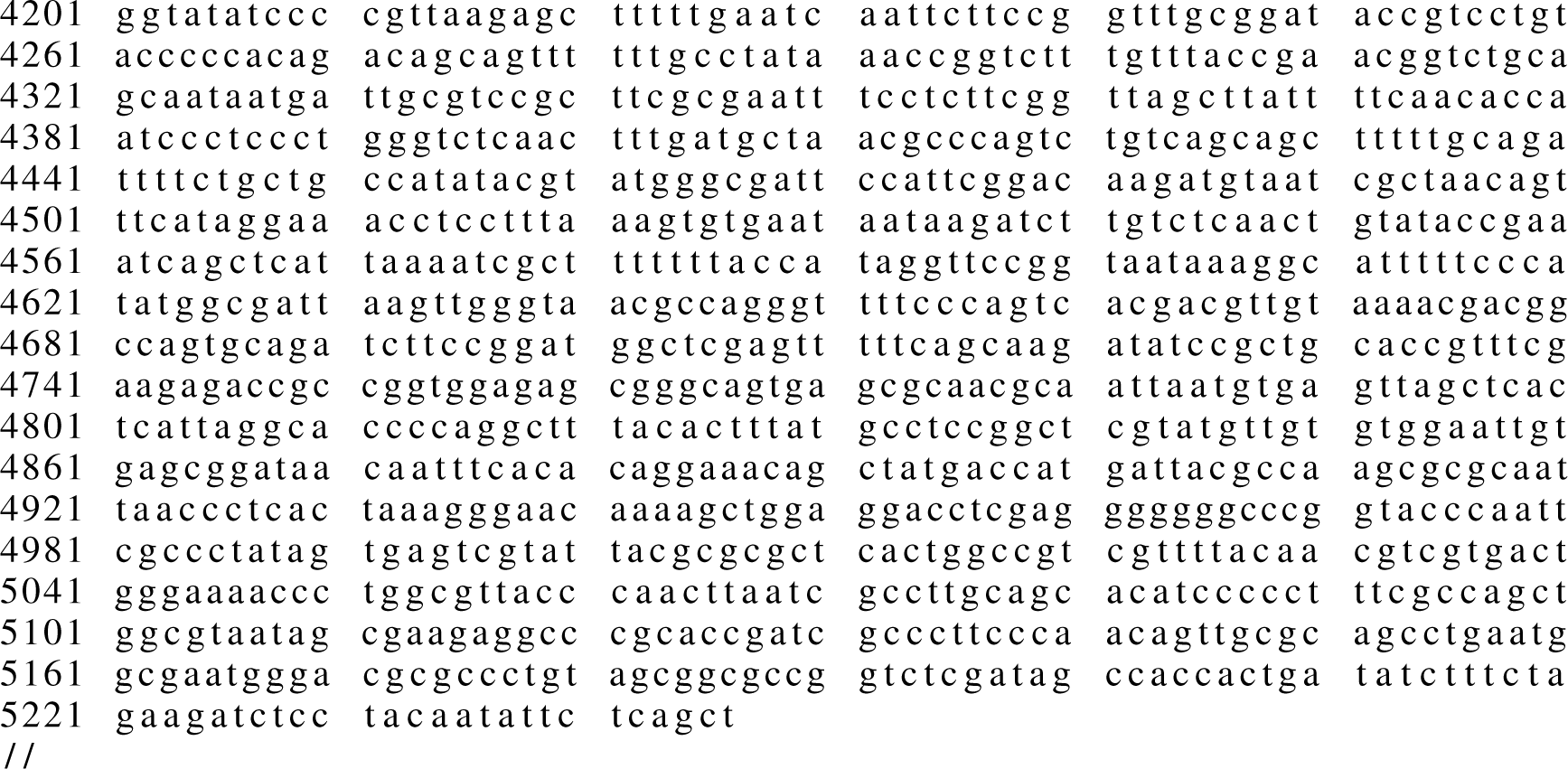

